# The mechanism of SARS-CoV-2 nucleocapsid protein recognition by the human 14-3-3 proteins

**DOI:** 10.1101/2020.12.26.424450

**Authors:** Kristina V. Tugaeva, Dorothy E. D. P. Hawkins, Jake L. R. Smith, Oliver W. Bayfield, De-Sheng Ker, Andrey A. Sysoev, Oleg I. Klychnikov, Alfred A. Antson, Nikolai N. Sluchanko

## Abstract

The coronavirus nucleocapsid protein (N) controls viral genome packaging and contains numerous phosphorylation sites located within unstructured regions. Binding of phosphorylated SARS-CoV N to the host 14-3-3 protein in the cytoplasm was reported to regulate nucleocytoplasmic N shuttling. All seven isoforms of the human 14-3-3 are abundantly present in tissues vulnerable to SARS-CoV-2, where N can constitute up to ~1% of expressed proteins during infection. Although the association between 14-3-3 and SARS-CoV-2 N proteins can represent one of the key host-pathogen interactions, its molecular mechanism and the specific critical phosphosites are unknown. Here, we show that phosphorylated SARS-CoV-2 N protein (pN) dimers, reconstituted via bacterial co-expression with protein kinase A, directly associate, in a phosphorylation-dependent manner, with the dimeric 14-3-3 protein, but not with its monomeric mutant. We demonstrate that pN is recognized by all seven human 14-3-3 isoforms with various efficiencies and deduce the apparent K_D_ to selected isoforms, showing that these are in a low micromolar range. Serial truncations pinpointed a critical phosphorylation site to Ser197, which is conserved among related zoonotic coronaviruses and located within the functionally important, SR-rich region of N. The relatively tight 14-3-3/pN association can regulate nucleocytoplasmic shuttling and other functions of N via occlusion of the SR-rich region, while hijacking cellular pathways by 14-3-3 sequestration. As such, the assembly may represent a valuable target for therapeutic intervention.

**Highlights:** SARS-CoV-2 nucleocapsid protein (N) binds to all seven human 14-3-3 isoforms. This association with 14-3-3 strictly depends on phosphorylation of N. The two proteins interact in 2:2 stoichiometry and with the Kd in a μM range. Affinity of interaction depends on the specific 14-3-3 isoform. Conserved Ser197-phosphopeptide of N is critical for the interaction.

## Introduction

The new coronavirus-induced disease, COVID19, has caused a worldwide health crisis with more than 90 million confirmed cases and 1.9 million deaths as of January 2021 [1]. The pathogen responsible, Severe Acute Respiratory Syndrome Coronavirus 2 (SARS-CoV-2), is highly similar to the causative agent of the SARS outbreak in 2002-2003 (SARS-CoV) and, to a lesser extent, to the Middle East Respiratory Syndrome Coronavirus (MERS-CoV) [2, 3]. Each is vastly more pathogenic and deadly than human coronaviruses HCoV-OC43, HCoV-NL63, HCoV-229E, and HCoV-HKU1 which cause seasonal respiratory diseases [4]. Like SARS-CoV and HCoV-NL63, SARS-CoV-2 uses angiotensin-converting enzyme 2 (ACE2) as entry receptor [5]. The ACE2 expression roughly correlates with the evidenced SARS-CoV-2 presence in different tissue types, which explains the multi-organ character of the disease [6] (Fig. 1).

**Fig. 1.**
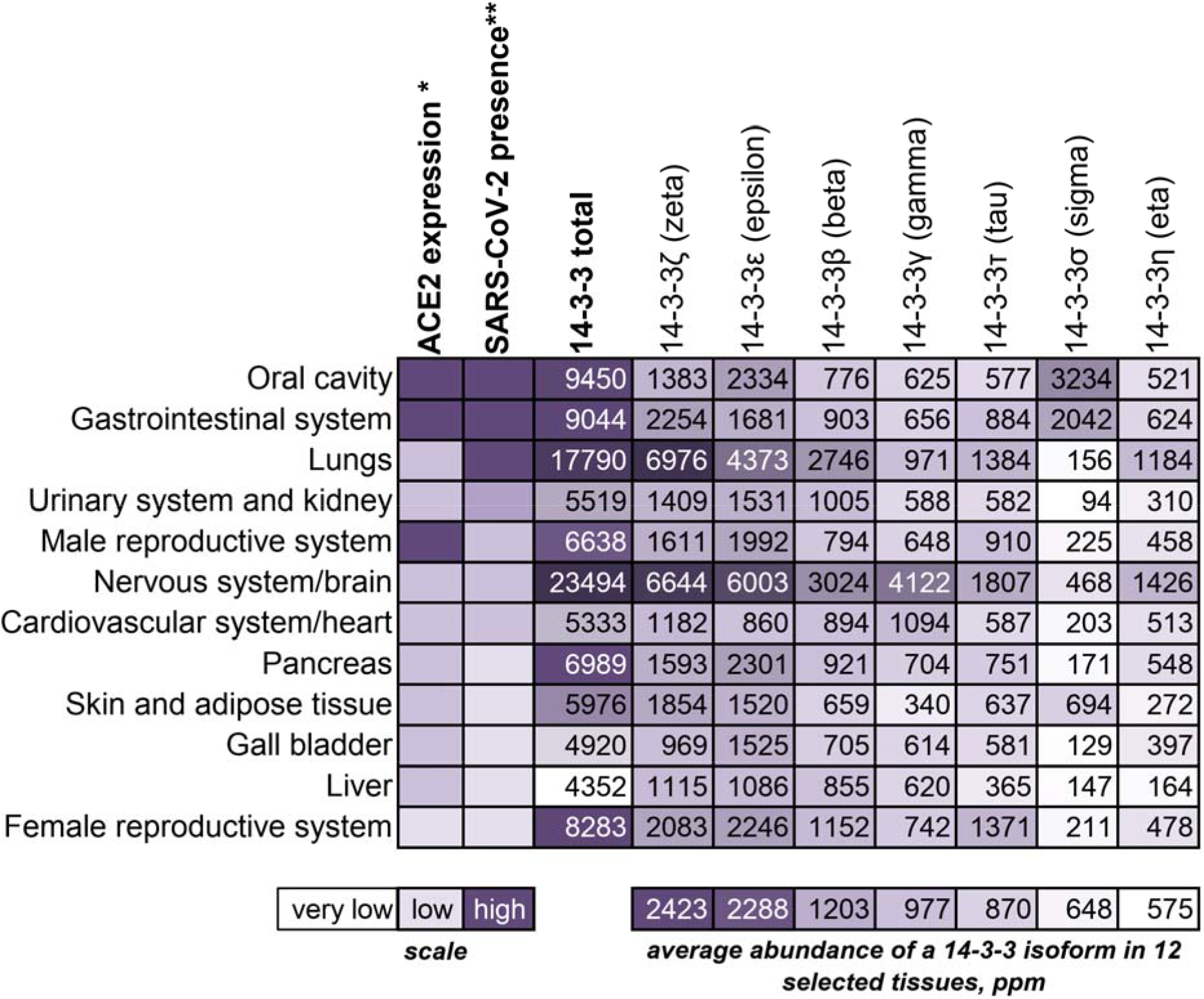
14-3-3 proteins are highly abundant in human tissues with SARS-CoV-2 presence. Correlation of ACE2 expression levels (*) and SARS-CoV-2 reported presence (**) in various tissues of COVID19 patients based on the data from [6], shown with abundances (indicated in ppm, part per million, i.e., one molecule of a given protein per 1 million of all proteins from a given tissue) of the seven human 14-3-3 isoforms, extracted from the PAXdb database [40]. Different tissues are shown in the order corresponding to the SARS-CoV-2 presence, starting, at the top, from the highest virus presence [6]. The shown relative scale of ACE2 expression is also taken from [6]. The total abundance of the seven human 14-3-3 isoforms in a given tissue and the average abundance of an isoform in 12 selected tissues are also indicated. The latter values were used for ordering the data for 14-3-3 isoforms, left to right, from the highest average abundance (14-3-3ζ; 2423 ppm, or ~0.24%) to the lowest average abundance (14-3-3η; 575 ppm, or ~0.06%). Note that the average abundance of all seven 14-3-3 proteins in three tissues with the highest SARS-CoV-2 presence (oral cavity, gastrointestinal tract, lungs) reaches 1.21% of all proteins.

In contrast to multiple promising COVID19 vaccine clinical trials [7–9], treatment of the disease is currently limited by the absence of approved efficient drugs [10]. The failure of several leading drug candidates in 2020 warrants the search for novel therapeutic targets including not only viral enzymes, but also heterocomplexes involving viral and host cell proteins. Unravelling mechanisms of interaction between the host and pathogen proteins may provide the platform for such progress.

The positive-sense single-stranded RNA genome of SARS-CoV-2 coronavirus encodes approximately 30 proteins which enable cell penetration, replication, viral gene transcription and genome assembly amongst other functions [11]. The 46-kDa SARS-CoV-2 Nucleocapsid (N) protein is 89.1% identical to SARS-CoV N. Genomic analysis of human coronaviruses indicated that N might be the major factor conferring the enhanced pathogenicity to SARS-CoV-2 [4]. N represents the most abundant viral protein in the infected cell [12–14], with each assembled virion containing approximately one thousand molecules of N [15]. Given that each infected cell can contain up to 10^5^ virions (infectious, defective and incomplete overall) [14], the number of N molecules in an infected cell can reach 10^8^, accounting for ~1% of a total number of cellular proteins (~10^10^ [16]).

The N protein interacts with viral genomic RNA, the membrane (M) protein and self-associates to provide for the efficient virion assembly [17–19]. It consists of two structured domains and three unstructured regions, including a functionally important central Ser/Arg-rich region (Fig. 2A) [20–22]. Such organization allows for a vast conformational change, which in combination with positively charged surfaces [23], facilitates nucleic acid binding [24]. Indeed, the crystal structure of the N-terminal domain (NTD) reveals an RNA binding groove [25–27], while crystal structures of the C-terminal domain (CTD) show a highly interlaced dimer with additional nucleic acid binding capacity [28, 29]. The N protein shows unusual properties in the presence of RNA, displaying concentration-dependent liquid-liquid phase separation [22, 23, 30, 31] that is pertinent to the viral genome packaging mechanism [32, 33]. In human cells, the assembly of condensates is down-regulated by phosphorylation of the SR-rich region [30, 34]. SARS-CoV-2 N protein is a major target of phosphorylation by host cell protein kinases, with 22 phosphosites identified in vivo throughout the protein (Supplementary table 1 and [13, 35]). Functions of N and viral replication can be regulated by a complex, hierarchical phosphorylation of the SR-rich region of SARS-CoV-2 N by a cascade of protein kinases [36]. Nevertheless, the potential functional role of N phosphorylation at each specific site is not understood.

**Fig. 2.**
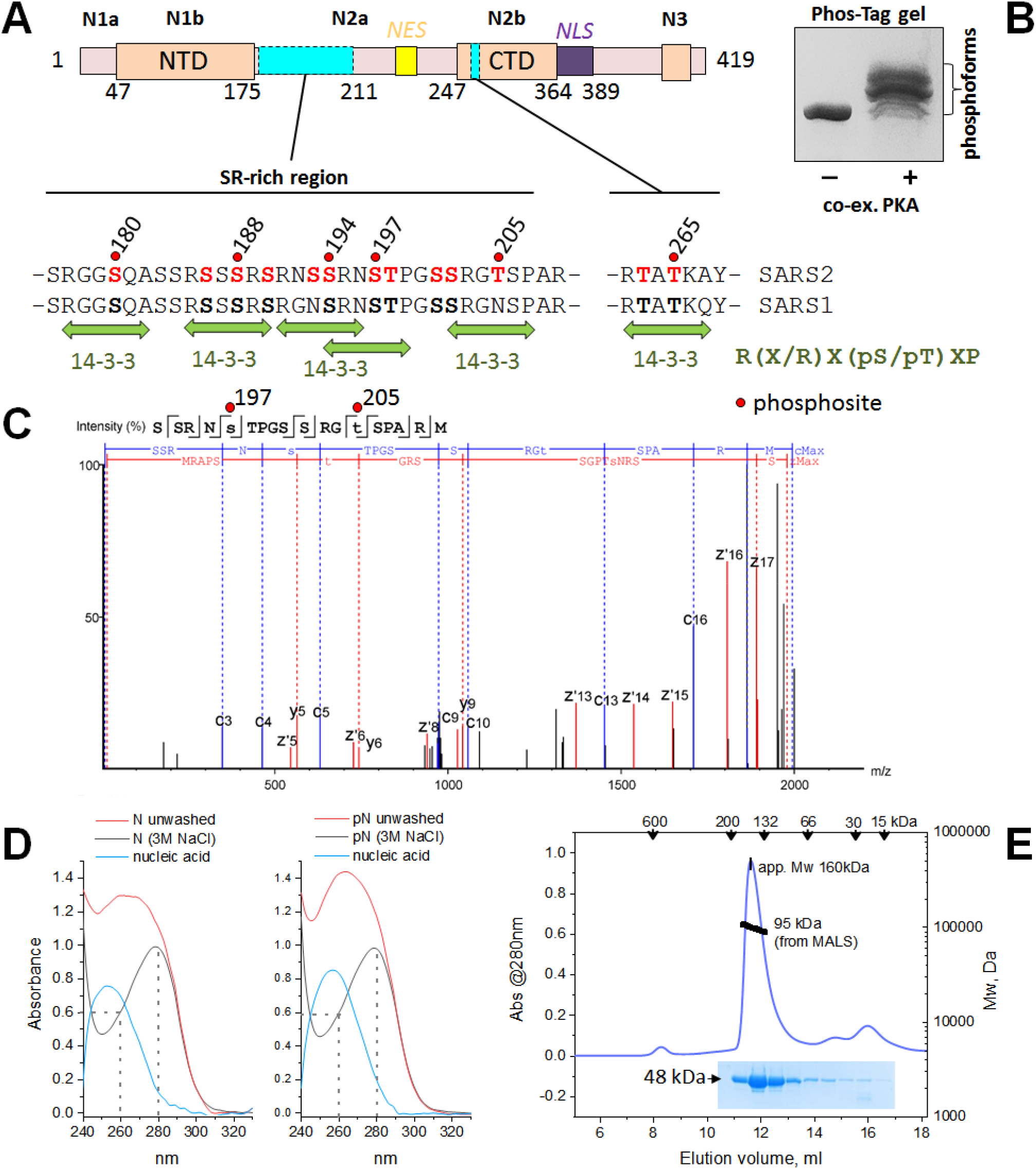
Characterization of the SARS-CoV-2 N phosphoprotein. A. Schematic representation of the N sequence and its main features. The subdomains are named above, predicted Nuclear Export Signal (NES, yellow), Nuclear Localization Signal (NLS, violet), N- and C-terminal domains (NTD, CTD) and two main phosphorylation loci (cyan rectangles) are marked. Sequences of the SR-rich region and short phosphorylatable section of the CTD are shown aligned between SARS-CoV and SARS-CoV-2 N proteins, highlighting multiple conserved phosphorylation sites (bold font indicates phosphosites present in both N proteins and experimentally confirmed for SARS-CoV-2 N). Some phosphosites (red spheres and labeled) are predicted as suboptimal 14-3-3 binding sites (green arrows). B. A Phos-tag gel showing that bacterial co-expression of the full-length N with PKA yields polyphosphorylated protein. C. A fragmentation spectrum of the representative phosphopeptide carrying phosphorylation at Ser197 and Thr205. Z- (red) and c-series (blue) of ETD fragmentation are shown. Error did not exceed 5 ppm. D. Absorbance spectra show that both recombinant unphosphorylated and PKA-phosphorylated N proteins elute from the Ni-affinity column bound to random *E.coli* nucleic acid. On-column washing with 3 M NaCl (50 column volumes) eliminates bound nucleic acid. E. Analysis of the oligomeric state of pN using size-exclusion chromatography on a Superdex 200 Increase 10/300 column at 200 mM NaCl, with multi-angle light scattering detection (SEC-MALS) and SDS-PAGE of the eluted fractions. Flow rate was 0.8 ml/min. Apparent Mw determined from column calibration based on protein standards (arrows above) is compared to the absolute mass determined from SEC-MALS. The experiments were carried out three times, and the most typical results are shown.

Using immunofluorescence, immunoprecipitation, siRNA silencing and kinase inhibition, it has been shown that SARS-CoV N protein shuttles between the nucleus and the cytoplasm in COS-1 cells [37]. This process is regulated by N protein phosphorylation by several protein kinases including glycogen synthase kinase-3, protein kinase A, casein kinase II and cyclin-dependent kinase [37–39]. Consequently, phosphorylated N associates with 14-3-3 proteins in the cytoplasm [37]. Notably, treatment with a kinase inhibitor cocktail eliminated the N/14-3-3 interaction, whereas inhibition of 14-3-3θ expression by siRNA led to accumulation of N protein in the nucleus [37]. These data suggest that 14-3-3 proteins directly shuttle SARS-CoV N protein in a phosphorylation-dependent manner: a role which may be universal for N proteins of all coronaviruses, including SARS-CoV-2. However, the molecular mechanism of the 14-3-3/N interaction remains ill-defined.

14-3-3 proteins are amongst the top 1% of highest-expressed human proteins in many tissues, with particular abundance in tissues vulnerable to SARS-CoV-2 infection including the lungs, gastrointestinal system and brain [40, 41] (Fig. 1). 14-3-3 proteins recognize hundreds of phosphorylated partner proteins involved in a magnitude of cellular processes ranging from apoptosis to cytoskeleton rearrangements [42, 43]. Human 14-3-3 proteins are present in most of the tissues as seven conserved “isoforms” (β, γ, ε, ζ, η, σ, τ/θ) (Fig. 1), with all-helical topology, forming dimers possessing two identical antiparallel phosphopeptide-binding grooves, located at ~35 Å distance from each other [44]. By recognizing phosphorylated Ser/Thr residues within the structurally flexible (R/K)X_2-3_(pS/pT)X(P/G) consensus motif [44, 45], 14-3-3 binding is known to regulate the stability of partner proteins, their intracellular localization and interaction with other factors [46]. In addition to their high abundance in many tissues susceptible to SARS-CoV-2 infection (Fig. 1), 14-3-3 proteins were reported as one of nine key host proteins during SARS-CoV-2 infection [47], indicating their potential association with viral proteins.

In this work, we dissected the molecular mechanism of the interaction between SARS-CoV-2 N and human 14-3-3 proteins. SARS-CoV-2 N protein containing several phosphosites reported to occur during infection, was produced using the efficient *Escherichia coli* system that proved successful for the study of polyphosphorylated proteins [48]. We have observed the direct phosphorylation-dependent association between polyphosphorylated SARS-CoV-2 N and all seven human 14-3-3 isoforms and determined the affinity and stoichiometry of the interaction. Series of truncated mutants of N localized the key 14-3-3-binding site to a single phosphopeptide residing in the functionally important SR-rich region of N. These findings suggest a topology model for the heterotetrameric 14-3-3/pN assembly occluding the SR-region, which presents a feasible target for further characterization and therapeutic intervention.

## Results

### 1. Characterization of the polyphosphorylated N protein obtained by co-expression with PKA in *E.coli*

Host-expressed SARS-CoV-2 N protein represents a phosphoprotein, harboring multiple phosphorylation sites scattered throughout its sequence. The most densely phosphorylated locus is the SR-rich region (Fig. 2A, Supplementary table 1, Supplementary data file 1 and [13, 35]). Remarkably, this region is conserved in N proteins of several coronaviruses [39, 49], including SARS-CoV (Fig. 2A). Although a number of protein kinases have been implicated in SARS-CoV N phosphorylation [36, 37, 39, 50], the precise enzymes responsible for identified phosphosites and the functional outcomes are largely unknown. Of note, many of the reported phosphosites within the SARS-CoV-2 N protein are predicted to be phosphorylated by protein kinase A (PKA), (Supplementary table 1). Hence, PKA was used for production of phosphorylated SARS-CoV-2 N in *E.coli* [48], using the same approach that was successfully applied for production of several phosphorylated eukaryotic proteins [48, 51–54] including the polyphosphorylated human tau competent for specific 14-3-3 binding [48].

Indeed, co-expression with PKA yielded a heavily phosphorylated SARS-CoV-2 N (Fig. 2B) containing more than 20 phosphosites according to LC-MS analysis (Supplementary table 1 and Supplementary data file 1). Especially dense phosphorylation occurred within the SR-rich region, involving recently reported in vivo sites Ser23, Thr24, Ser180, Ser194, Ser197, Thr198, Ser201, Ser202, Thr205 and Thr391 [13, 35] (Fig. 1A, C and Supplementary data file 1), implicating the success of the PKA co-expression at emulating native phosphorylation. Interestingly, due to the frequent occurrence of Arg residues, it was possible to characterize the polyphosphorylation of the SR-rich region only with the use of an alternative protease such as chymotrypsin, in addition to datasets obtained separately with trypsin (4 independent experiments overall, see Supplementary data file 1). Due to high conservation between SARS-CoV and SARS-CoV-2 N proteins, many phosphosites identified in the PKA-co-expressed N are likely shared by SARS-CoV N (Fig. 2A). Importantly, many identified phosphosites lie within the regions predicted to be disordered, and contribute to predicted 14-3-3-binding motifs, albeit deviating from the optimal 14-3-3-binding sequence RXX(pS/pT)X(P/G) [44] (Fig. 2A and Supplementary table 1).

Of note, the bacterially expressed SARS-CoV-2 N protein avidly binds random *E.coli* nucleic acid, which results in a high 260/280 nm absorbance ratio in the eluate from the nickel-affinity chromatography column. This is unchanged by polyphosphorylation (Fig. 2D) and nucleic acid remained bound even after further purification using heparin chromatography (data not shown). To quantitatively remove nucleic acid from N preparations, we used continuous on-column washing of the His-tagged protein with 3 M salt. This yielded clean protein preparations with the 260/280 nm absorbance ratio of 0.6 (Fig. 2D) of the unphosphorylated and polyphosphorylated N (pN) with high electrophoretic homogeneity (Fig. 2E), enabling thorough investigation of the 14-3-3 binding mechanism.

SEC-MALS suggested that the nucleic acid-free protein was a ~95 kDa dimer (Fig. 2E), based on the calculated Mw of the His-tagged N monomer of 48 kDa, regardless of its phosphorylation status. A significant overestimation of the apparent Mw of pN from column calibration, ~160 kDa for the 95 kDa dimeric species, indicates presence of elements with expanded loose conformation, in agreement with the presence of unstructured regions. This necessitates the use of SEC-MALS for absolute Mw determination that is independent of the assumptions on density and shape.

The skewed Mw distribution across the SEC peak indicated the propensity of the SARS-CoV-2 N to oligomerization (Fig. 2E), which was instigated by the addition of nucleic acid (Supplementary Fig. 1A and B). Using tRNA isolated from *E.coli* DH5LJ cells, SEC and agarose gel electrophoresis, we showed that our nucleic acid-free, polyphosphorylated N preparation is capable of avid binding of noncognate nucleic acid (Supplementary Fig. 1A-C).

### 2. Polyphosphorylated SARS-CoV-2 N and human 14-3-3γ form a tight complex with defined stoichiometry

Next, we compared the ability of native full-length SARS-CoV-2 N, both unphosphorylated and polyphosphorylated (N.1-419 and pN.1-419, respectively), to be recognized by a human 14-3-3 protein. For the initial analysis we chose 14-3-3γ as one of the strongest phosphopeptide binders among the 14-3-3 family [55].

SEC-MALS (Fig. 3A) shows that 14-3-3γ elutes as a dimer with Mw of ~55.2 kDa (calculated monomer Mw 28.3 kDa, see also Fig. 3B). The position and amplitude of this peak did not change in the presence of the N.1-419 dimer with Mw of 94.5 kDa (calculated monomer Mw 48 kDa) where the SEC profile shows two distinct peaks. This is corroborated by SDS-PAGE analysis, suggesting no interaction between unphosphorylated N (pI >10) and 14-3-3 (pI ~4.5) despite the large difference in their pI values (Fig. 3C,D). Thus, the presence of 200 mM NaCl was sufficient to prevent nonspecific interactions between the two proteins.

**Fig. 3.**
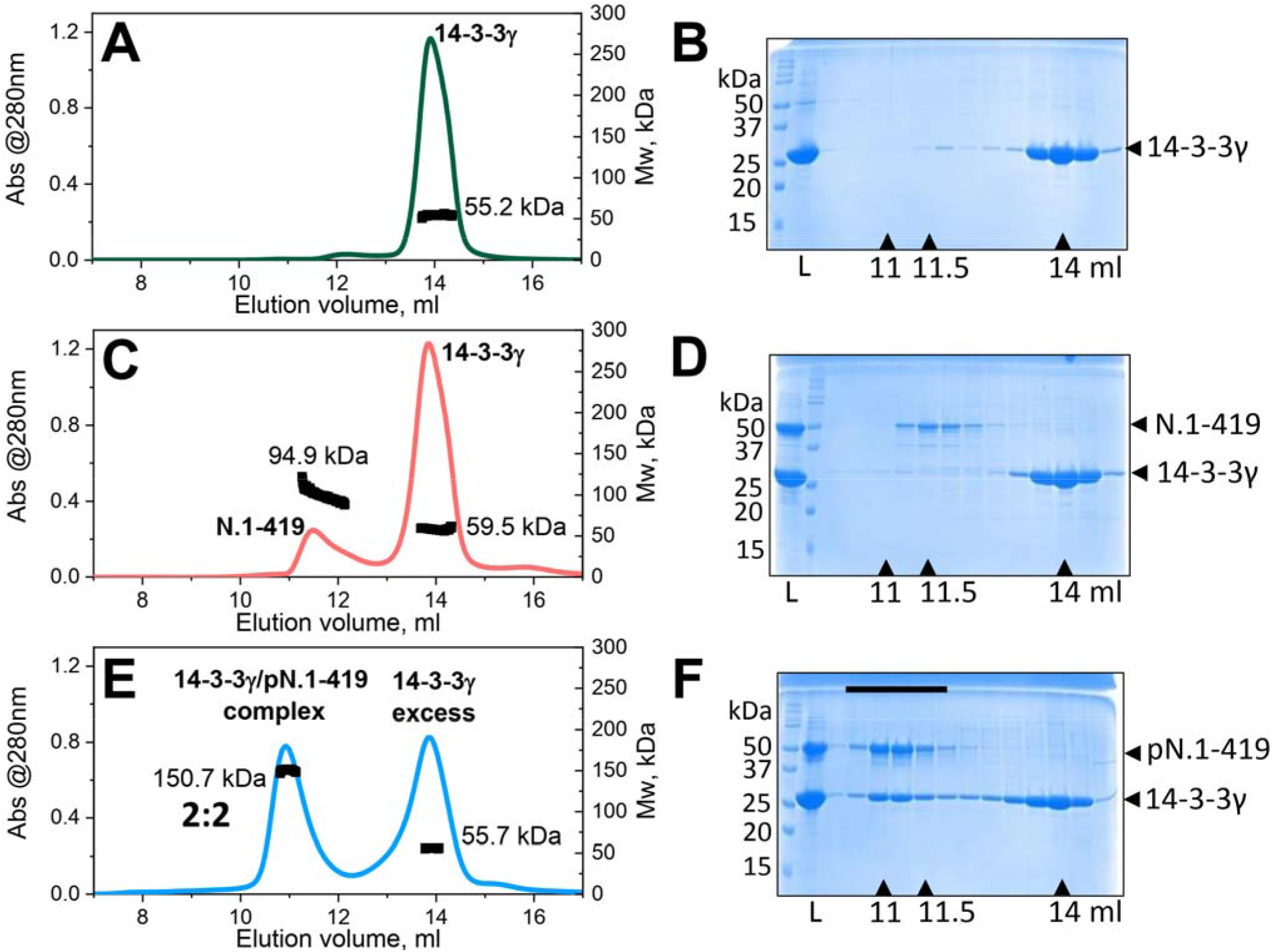
Interaction of SARS-CoV-2 N with human 14-3-3γ. Interaction between SARS-CoV-2 N protein (50 μM) and human 14-3-3γ (160 μM) was analyzed by SEC-MALS and SDS-PAGE of the eluted fractions. SEC-MALS/SDS-PAGE of 14-3-3γ alone (A,B), with unphosphorylated N (C,D) and with phosphorylated N (E,F) are presented. Mw distributions obtained using MALS are shown on each peak. “L” shows contents of the samples loaded on a column, elution volumes corresponding to the maxima of the peaks are indicated by arrows below the gels. Mw markers are shown to the left of each gel. A black bar on panel F indicates the fractions of the complex between 14-3-3γ and pN. A Superdex 200 Increase 10/300 column equilibrated with 20 mM Tris-HCl buffer pH 7.6 containing 200 mM NaCl and 3 mM NaN_3_ was operated at a flow rate of 0.8 ml/min. The experiment was performed three times and the most typical results are shown.

In sharp contrast, co-expression of SARS-CoV-2 N with PKA, and subsequent polyphosphorylation (Fig. 2), allows for tight complex formation between pN.1-419 and human 14-3-3γ. This is evident from the peak shift and corresponding increased Mw from ~95 to 150.7 kDa (Fig. 3E), perfectly matching the addition of the 14-3-3 dimer mass (calculated dimer Mw 56.6 kDa) to the pN.1-419 dimer (calculated dimer Mw 96 kDa). The presence of both proteins in the complex was confirmed by SDS-PAGE (Fig. 3F). Collectively these data pointed toward the equimolar binding upon saturation. Given the dimeric state of both proteins in their individual states (Fig. 2D and Fig. 3) and the Mw of the 14-3-3γ/pN.1-419 complex, the most likely stoichiometry is 2:2. The ratio of the proteins does not change across the peak of the complex (Fig. 3F), implying that they form a relatively stable complex with the well-defined stoichiometry.

### 3. SARS-CoV-2 N interacts with all human 14-3-3 isoforms

We then questioned whether the interaction with pN is preserved for other human 14-3-3 isoforms. Analytical SEC clearly showed that the phosphorylated SARS-CoV-2 N can be recognized by all seven human 14-3-3 isoforms, regardless of the presence of a His-tag or disordered C-terminal tails on the corresponding 14-3-3 constructs (Fig. 4A). However, the efficiency of complex formation differed for each isoform. Judging by the repartition of 14-3-3 between free and the pN-bound peaks, the apparent efficiency of pN binding was higher for 14-3-3γ, 14-3-3η, 14-3-3ζ and 14-3-3ε, and much lower for 14-3-3β, 14-3-3τ and 14-3-3σ, in a roughly descending order (Fig. 4A). The interaction also appeared dependent on the oligomeric state of 14-3-3, since the monomeric mutant form of 14-3-3ζ, 14-3-3ζm-S58E [56] (apparent Mw 29 kDa) showed virtually no interaction relative to the wild-type dimeric 14-3-3ζ counterpart (apparent Mw 58 kDa), (Fig. 4B).

**Fig. 4.**
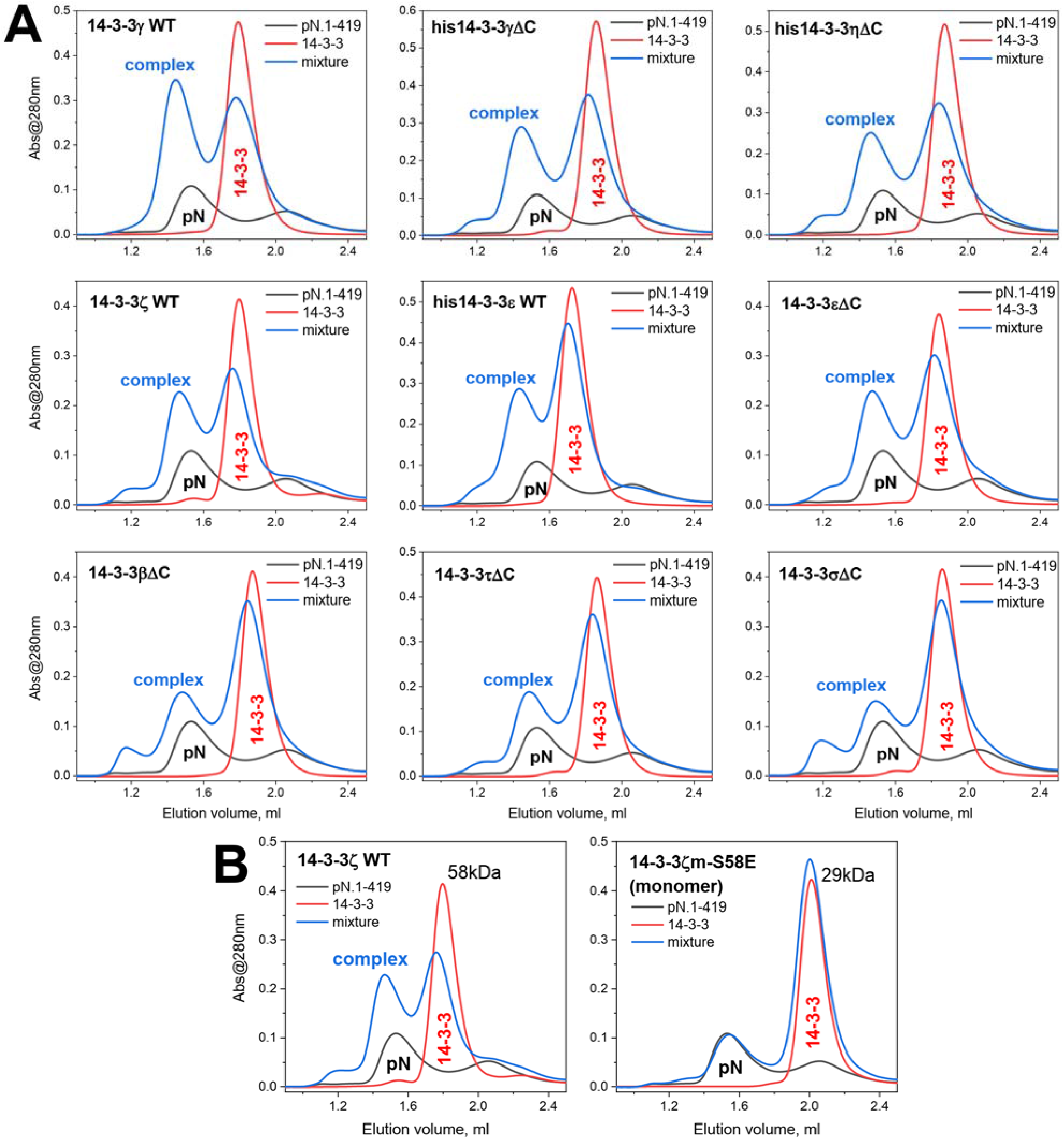
SARS-CoV-2 N interacts with all seven dimeric human 14-3-3 isoforms, but not with the monomeric mutant of 14-3-3ζ. A. Interaction between phosphorylated N protein (46.6 μM) and human isoforms 14-3-3γ (69.4 μM), his-tagged 14-3-3γΔC (73.9 μM), his-tagged 14-3-3ηΔC (73.8 μM), 14-3-3ζ (77.6 μM), his-tagged 14-3-3ε (76.2 μM), 14-3-3εΔC (60.3 μM), 14-3-3βΔC (69.4 μM), 14-3-3τΔC (71.7 μM), 14-3-3σΔC (75.4 μM) studied by analytical SEC on a Superdex 200 Increase 5/150 column at 200 mM NaCl. B. Phosphorylated N protein (46.6 μM) interacts with the wild-type dimeric 14-3-3ζ (77.6 μM) but not with its monomeric mutant 14-3-3ζm-S58E (74.9 μM). Apparent Mw indicated in kDa were determined from column calibration.

This apparent variation in binding efficiency (Fig. 4A) justified more detailed assessment of the binding parameters between pN.1-419 and selected 14-3-3 isoforms.

### 4. Affinity of the phosphorylated SARS-CoV-2 N towards selected human 14-3-3 isoforms

In light of the relative positions of the two proteins, separately and complexed, on SEC profiles (Fig. 3 and 4), we used analytical SEC to track titration of a fixed concentration of pN.1-419 (around 10 μM) against increasing quantities (0-100 μM) of either of two selected full-length human 14-3-3 isoforms, γ and ε (Fig. 5A and B). A saturation binding curve showed the maximal concentration of bound 14-3-3γ to asymptotically approach 10 μM (Fig. 5C), supporting 2:2 stoichiometry. The apparent K_D_ of the 14-3-3γ/pN.1-419 complex was estimated as 1.5 ± 0.3 μM. A similar binding mechanism could be observed for 14-3-3ε, however in this case we could achieve pN.1-419 saturation only at much higher 14-3-3ε concentrations (Fig. 5B and C), and the resulting apparent K_D_ was ~7 times higher than for 14-3-3γ (Fig. 5C). Nevertheless, once again the stoichiometry was close to 2:2. These findings strongly disfavor the earlier hypothesis that 14-3-3 binding affects dimerization status of N [57].

**Fig. 5.**
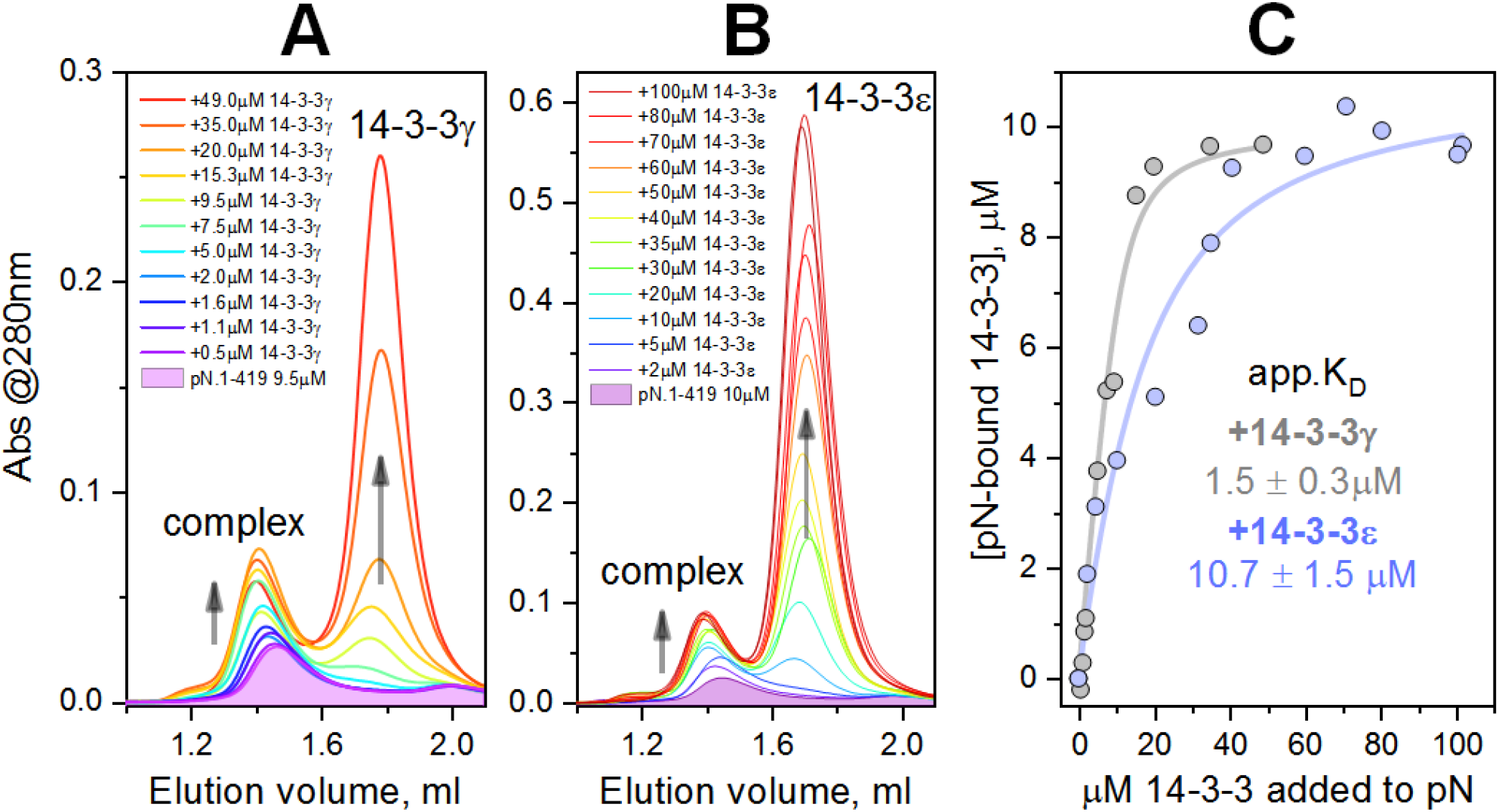
Binding affinity. pN at a fixed concentration (~10 μM) was titrated against either of the two human 14-3-3 isoforms and monitored by analytical SEC with serial sample loading. A. Titration of pN with 14-3-3γ. B. Titration of pN with 14-3-3ε. Changes of the elution profiles associated with the increasing 14-3-3 concentration are shown by arrows. C. Binding curves used for apparent K_D_ determination. Note that, regardless of the observed differences in the affinities, the maximum 14-3-3 concentration complexed with pN equals the concentration of pN (~10 μM) for both 14-3-3 isoforms. Typical results from two independent experiments are shown.

We further asked what are the specific regions of SARS-CoV-2 N that are responsible for interaction with human 14-3-3.

### 5. The N-terminal part of SARS-CoV-2 N is responsible for 14-3-3 binding

Among multiple phosphosites identified in our pN.1-419 preparations (Supplementary table 1), the two most interesting regions are located in the intrinsically disordered or loop segments, normally favored by 14-3-3 proteins [45]. The first represents the C-terminally located RTA[pT^265^]KAY site, which is predicted by the 14-3-3-Pred webserver [58] as the 14-3-3-binding site. The second, SR-rich region features multiple experimentally confirmed phosphosites including several suboptimal predicted 14-3-3-binding sites (Fig. 2A). To narrow down the 14-3-3-binding locus we used several constructs representing its N- and C-terminal parts (N.1-211 and N.212-419, respectively). The individual CTD included the C-terminal phosphosite around Thr265 (N.247-364), and the longer N-terminal construct extended toward the C-terminus to include the predicted NES sequence (N.1-238) (Fig. 6A). This contains Asp225 and a cluster of Leu residues which together resemble the unphosphorylated 14-3-3-binding segments from ExoS/T [59] and therefore could be important for 14-3-3 binding.

**Fig. 6.**
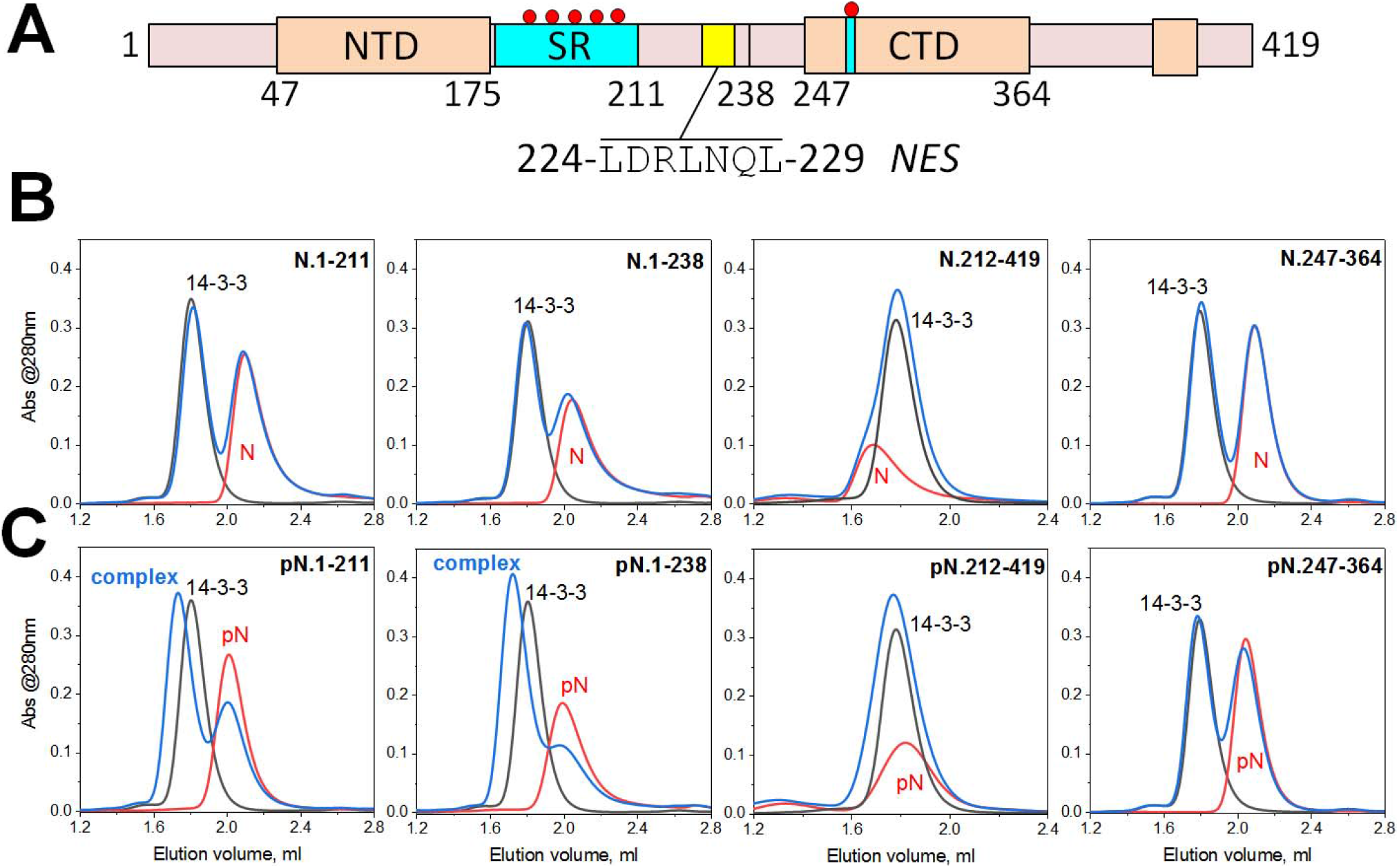
Interaction of human 14-3-3γ with individual SARS-CoV-2 N protein domains. A. Schematic representation of the N protein sequence with the main domains/regions highlighted. Yellow indicates NES, small red circles designate phosphosites within the two corresponding phosphorylatable loci. B. SEC profiles of individual 14-3-3γ (black traces), individual N (red traces), and 14-3-3 with N (blue traces) obtained using a Superdex 200 Increase 5/150 column at 200 mM NaCl in running buffer and operated at a 0.45 ml/min flow rate. The upper row corresponds to unphosphorylated N fragments, the bottom row corresponds to their phosphorylated counterparts, as indicated on each panel. In all cases, 50 μM of N was mixed with 50 μM of 14-3-3 and loaded in an equal volume of 50 μl. In the case of N.247-364 and pN.247-364, 90 μM of N was used. Note that the well-defined complexes with 14-3-3 were observed only for pN.1-211 and pN.1-238 fragments. The experiment was repeated twice and the most typical results are presented.

As for the wild-type protein, the truncated SARS-CoV-2 N constructs were cloned and expressed in the absence or presence of PKA to produce unphosphorylated or polyphosphorylated proteins. PKA could be easily detected in the eluate from the Ni-affinity column and typically led to an upward shift on SDS-PAGE and a shift on the SEC profile (Supplementary Fig. 2). Each indicates efficient phosphorylation. Perhaps with the exception of N.212-419, phosphorylation did not affect the oligomeric state of the N constructs, and the observed shifts could be accounted for by the affected flexible regions (Fig. 4 and Supplementary Fig. 2).

SEC-MALS (Supplementary Fig. 2) indicates the N.247-364 construct (calculated monomer Mw 15.3 kDa), previously crystallized and kindly provided by Prof. Baumeister’s laboratory, exists as a ~30 kDa dimer. This is in perfect agreement with earlier published data [28]. A dimeric state (Mw of 47.3 kDa) was also confirmed for the longer C-terminal construct, N.212-419 (Supplementary Fig. 3, calculated monomer Mw 23.3 kDa), however, in this case a skewed Mw distribution indicative of the polydispersity and tendency for oligomerization was observed (Supplementary Fig. 3).

In contrast, N-terminal constructs including the RNA-binding domain, such as N.1-211, are monomeric (Supplementary Fig. 3). Thus, our data align with the low-resolution structural model of SARS-CoV and SARS-CoV-2 N proteins, in which the CTD (residues 247-364), largely responsible for dimerization, and NTD, involved in RNA-binding, are only loosely associated [24, 28, 29].

Of the constructs analyzed, the N-terminal constructs N.1-211 and N.1-238 both interacted with 14-3-3γ by forming distinct complexes on SEC profiles, and this interaction was strictly phosphorylation-dependent (Fig. 6). Given the similarity of the elution profiles for pN.1-211 and pN.1-238, it may be concluded that the presence of NES in the latter is dispensable for the 14-3-3 binding.

Under similar conditions, only a very weak interaction between the dimeric pN.212-419 construct and 14-3-3γ could be observed, whereas the phosphorylated dimeric CTD (pN.247-364) displayed virtually no binding (Fig. 6). Neither unphosphorylated construct interacted with 14-3-3. Thus, the Thr265 phosphosite can be broadly excluded as the critical binding site. Separate phosphosites outside the CTD, for instance, in the last ~30 C-terminal residues (Supplementary table 1) likely account for residual binding of the pN.212-419 construct. Only its SEC profile showed a significant positional peak shift with phosphorylation (Fig. 6B and C), indicating a potential change in the oligomeric state. It is tempting to speculate that such phosphorylation outside the CTD could affect higher order oligomerization associated with the so-called N3 C-terminal segment (Fig. 2A) [20, 21, 49, 60].

According to SEC-MALS, the 14-3-3γ dimer (Mw 57 kDa) interacts with the pN.1-238 monomer (Mw of 31 kDa) by forming a ~82 kDa complex (Fig. 7A) with an apparent 2:1 stoichiometry. It is remarkable that despite a moderate molar excess of pN.1-238 a 2:2 complex (one 14-3-3 dimer with two pN monomers) is not observed. The well-defined 2:2 stoichiometry of the 14-3-3γ complex with the full-length pN (Fig. 3) suggests that the dimeric pN is anchored using two equivalent, key 14-3-3-binding sites, each located in a separate subunit of N. It is tempting to speculate that, in the absence of the second subunit, the interaction involves the key phosphosite and an additional phosphosite which is separated by a sufficiently long linker (≥ 15 residues [61]), to secure occupation of both phosphopeptide-binding grooves of 14-3-3 (Fig. 7B). Such bidentate binding would prevent the recruitment of a second pN.1-238 monomer and is in line with the observed data. Similar 2:1 binding was observed qualitatively for the interaction of 14-3-3γ with pN.1-211 (Fig. 6C and data not shown).

**Fig. 7.**
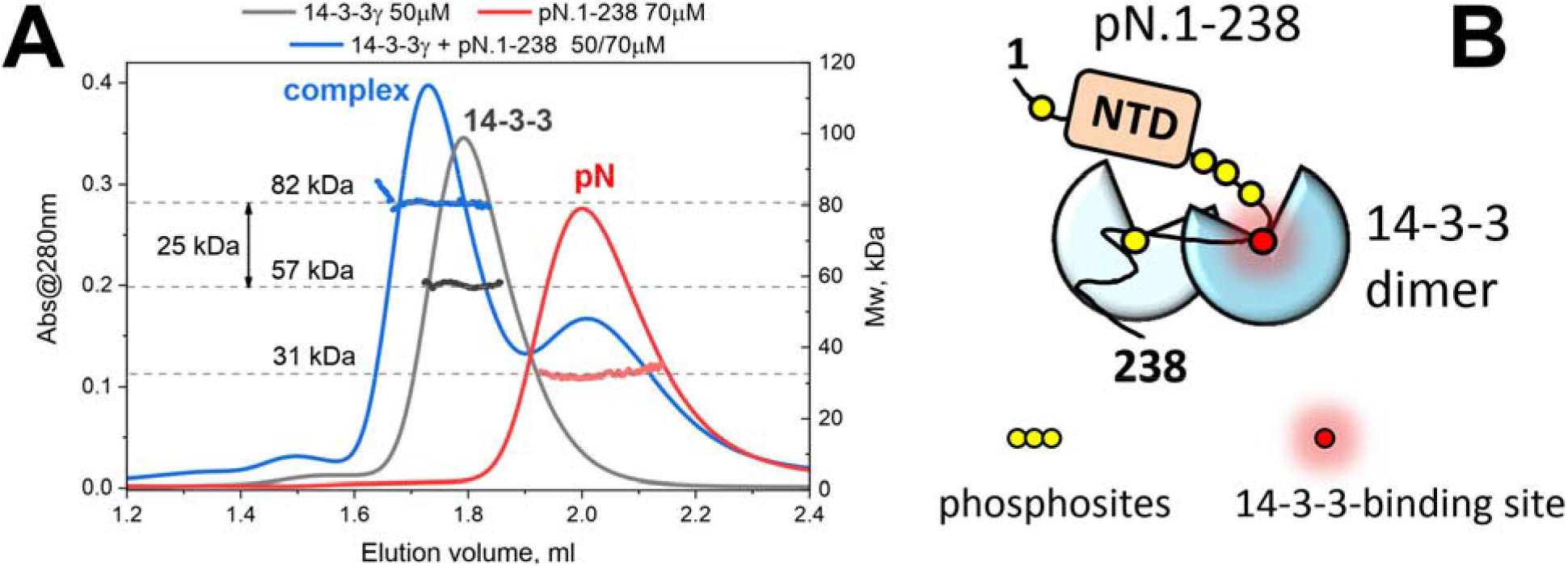
Interaction of human 14-3-3γ with the phosphorylated monomeric SARS-CoV-2 N construct 1-238. A. SEC-MALS analysis of individual pN, 14-3-3γ and their complex formed at the indicated molar excess of pN. Protein concentration, MALS-derived Mw distributions across the peaks and the respective average Mw values are indicated. Note that the difference shown by a two-headed arrow roughly corresponds to a pN.1-238 monomer mass. B. Schematic representation of the possible bidentate binding mode where the 14-3-3 dimer interacts with the tentative key 14-3-3-binding phosphosite and another sterically allowed phosphosite separated by a sufficiently long linker.

Our finding that the minimal N-terminal construct pN.1-211 exhibits firm binding to 14-3-3γ indicates that the key 14-3-3-binding phosphosite(s) is/are located exclusively within this region. Given the presence of numerous candidate 14-3-3-binding sites within its most C-terminal part, i.e., the SR-rich region, we further focused on the 1-211 sequence in search of the 14-3-3-binding phosphosite(s).

### 6. Localization of the main 14-3-3-binding site within the SR-rich region of SARS-CoV-2 N

Further N mutants were designed to disrupt the most probable 14-3-3-binding phosphosites. These are located in the intrinsically flexible phosphorylatable SR-rich segment centered at positions 197 and 205 (Fig. 8A). Both represent suboptimal 14-3-3-binding motifs **S**RN[pS^197^]T<u>**P**</u> and **S**RG[pT^205^]S<u>**P**</u> in lacking an Arg/Lys residue in position −3 (bold font) relative to the phosphorylation site (squared brackets). However, each also features a Pro residue in position +2 (bold underlined font), which is highly favorable for 14-3-3 binding [44] and absent from the other potential 14-3-3-binding phosphosites in the SR-rich region (Fig. 2A). These conflicting factors complicate predictions for the true 14-3-3 binding site. Moreover, even beyond the SR-rich region the NTD is predicted to host further possible 14-3-3-binding phosphosites, including the RRA[pT^91^]RR site, which is the highest-scoring in 14-3-3-Pred [58] prediction (Supplementary table 1).

**Fig. 8.**
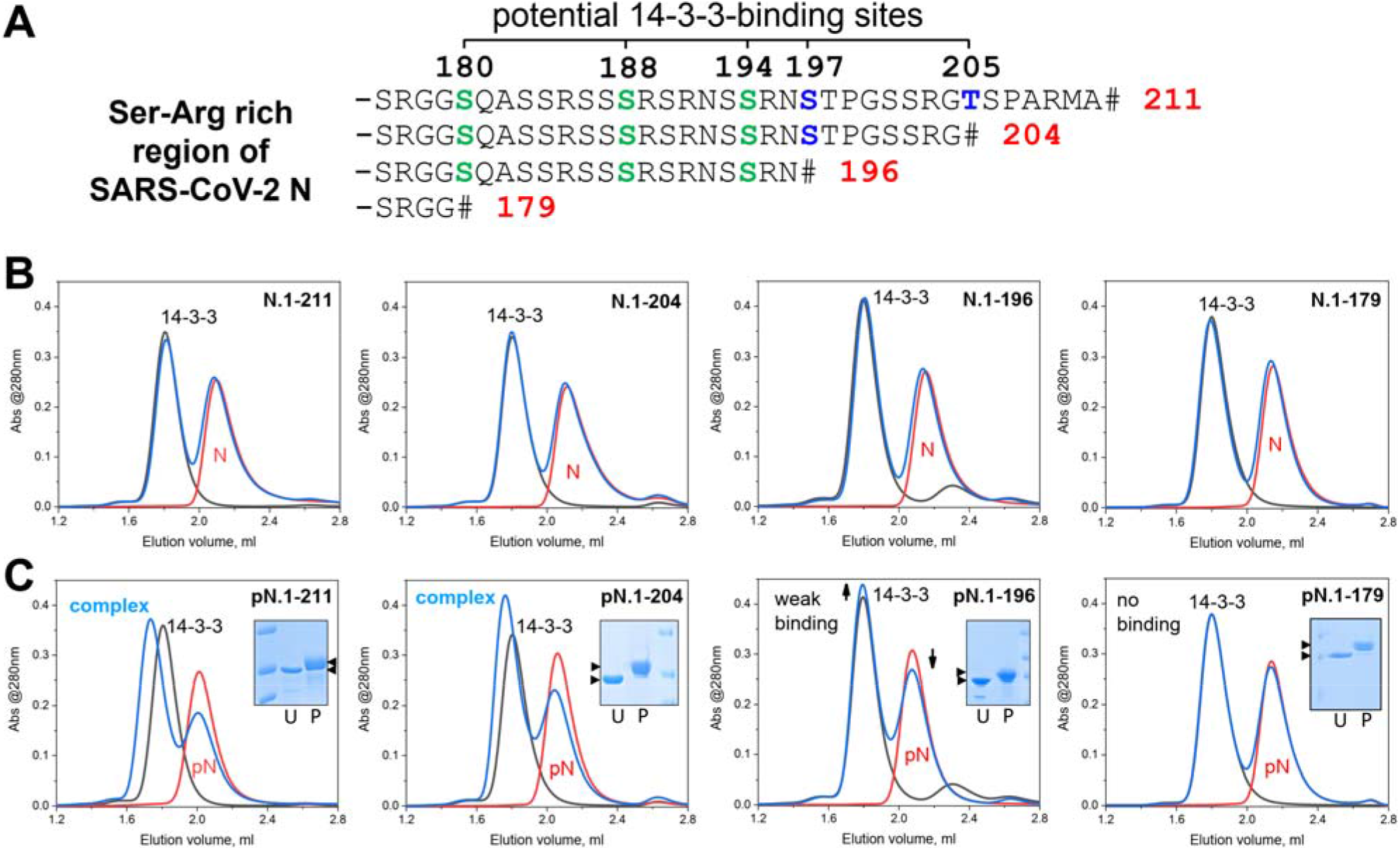
Localization of the 14-3-3-binding sites within the SR-rich region of the SARS-CoV-2 N protein. A. Sequence of the SR-rich region showing potential 14-3-3-binding sites (green and blue font) and the truncations designed to exclude one or two main 14-3-3-binding sites (blue font). # denotes the designed C-terminus. B, C. Analysis by SEC of the interaction between human 14-3-3γ and the truncated mutants of SARS-CoV-2 N in their unphosphorylated (B) or phosphorylated forms (C). Each graph contains SEC profiles for the individual 14-3-3 (black line), individual N or pN construct (red line), and the 14-3-3γ N/pN mixture (blue line) where 50 μM of each protein was used throughout. Inserts on panel C show the upward shift on SDS-PAGE as the result of phosphorylation (U, unphosphorylated; P, phosphorylated protein, arrowheads indicate the shift). Note that only the phosphorylated mutants interacted with 14-3-3γ and that only pN.1-211 and pN.1-204 formed a defined complex with 14-3-3γ. Column: Superdex 200 Increase 5/150, flow rate: 0.45 ml/min. The experiment was repeated twice and the most typical results are presented.

We conceived stepwise truncations to remove the most probable 14-3-3-binding phosphosites, aiming to identify the iteration at which binding (observed for the pN.1-211 construct) ceased. 14-3-3 can bind incomplete consensus motifs at the extreme C-terminus of some proteins [62], so truncations were designed to remove the critical phosphorylated residue. However, upstream residues of each candidate 14-3-3-binding site were preserved, in light of the sheer number of overlapping potential binding motifs in the SR-rich region (Fig. 8A).

The new truncated constructs of N, namely N.1-204, N.1-196 and N.1-179, were obtained in unphosphorylated and phosphorylated states, as before, and again washed with high salt to remove potentially disruptive nucleic acid. SEC-MALS confirmed the monomeric state of the N-terminal N mutants (the exemplary data are presented for N.1-196, Supplementary Fig. 3), consistent with the proposed architecture of the N protein [21, 24].

None of the truncated mutants interacted with 14-3-3γ in the unphosphorylated state (Fig. 8B). No binding to 14-3-3γ was detected with phosphorylated N.1-179 (Fig. 8C) and only very limited binding could be observed with phosphorylated N.1-196 (Fig. 8C). This strongly indicated that all phosphosites of the 1-196 segment (including at least three phosphosites within the SR-rich region, i.e., Ser180, Ser188 and Ser194, see Fig. 8A) are dispensable for 14-3-3 binding and at most could contribute only as auxiliary sites (as suggested by the scheme in Fig. 7B). This narrowed the 14-3-3-binding region within SARS-CoV-2 N down to 15 residues from 196 to 211, leaving only two possible sites centered at Ser197 and Thr205 (Fig. 8A).

By contrast, pN.1-204 showed a similar interaction with 14-3-3γ to that of pN.1-211 (Fig. 8C). Although we cannot fully exclude that Thr205 phosphosite contributes to 14-3-3 binding in the context of the full-length pN, it is clearly not critical for 14-3-3 recruitment, unlike Ser197. Moreover, in contrast to Thr205, Ser197 is preserved in most related coronavirus N proteins (see Fig. 2A, Fig. 9).

**Fig. 9.**
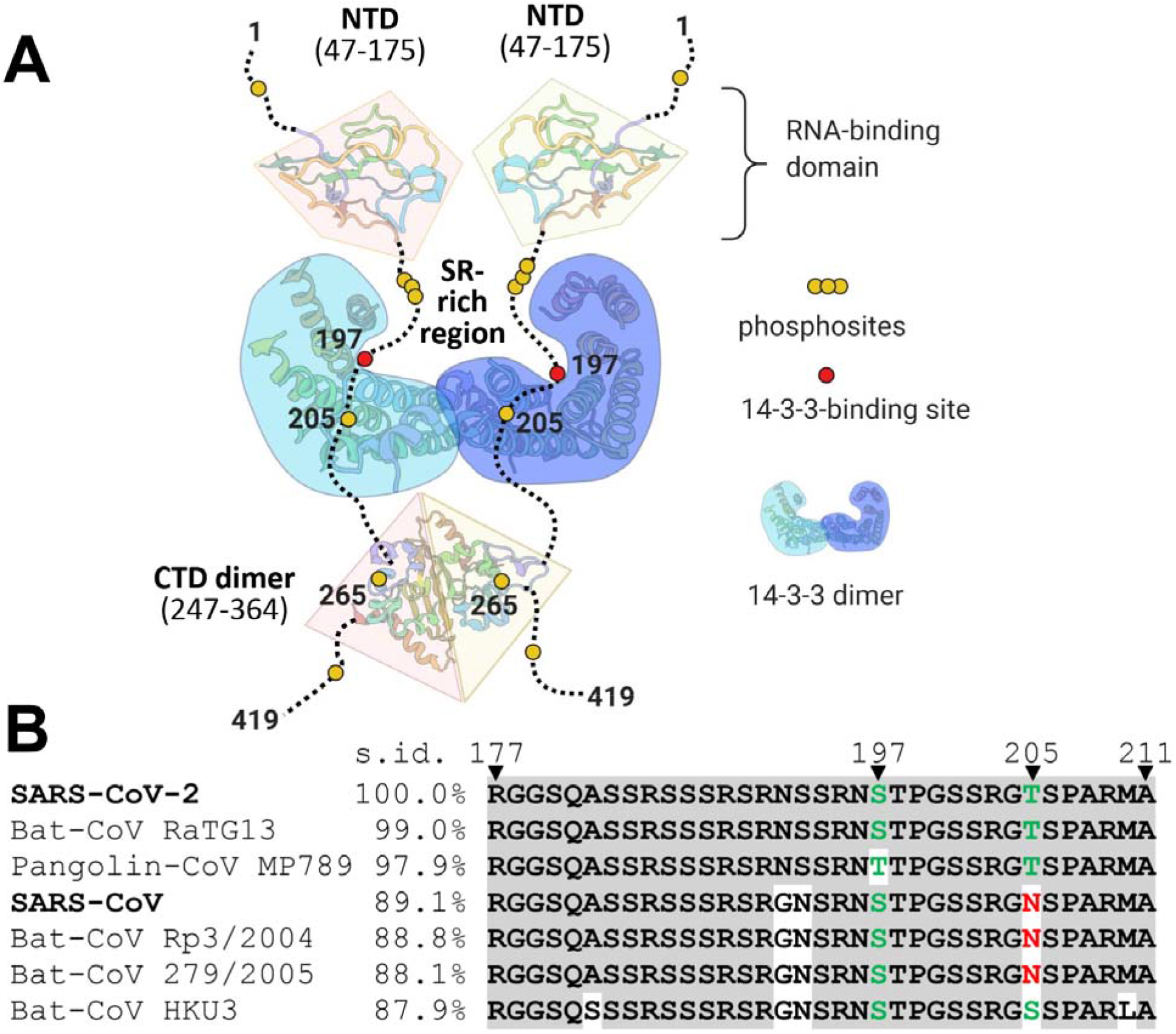
Association of SARS-CoV-2 N with human 14-3-3. A. A topology model for the complex of SARS-CoV-2 N protein dimer with the dimer of human 14-3-3 illustrating the occlusion of the SR-rich region. B. Local alignment of SR-rich regions of the most similar coronavirus N proteins in order of descending sequence identity (s.id.) determined using entire N protein sequences. Alignment was performed using Clustal omega and visualized using Mview (https://www.ebi.ac.uk/Tools/msa/mview/). Residues identical to those in the SARS-CoV-2 sequence are shadowed by grey, phosphorylatable residues in positions 197 and 205 are in green color, residues blocking phosphorylation in position 205 are in red.

## Discussion

In this work, we investigated the molecular association between the SARS-CoV-2 N protein and human phosphopeptide-binding proteins of the 14-3-3 family. The former is the most abundant viral protein [12, 13], the latter is a major protein-protein interaction hub involved in multiple cellular signaling cascades, expressed at high levels in many human tissues including those susceptible to SARS-CoV-2 infection (Fig. 1) [40]. SARS-CoV-2 N is heavily phosphorylated in infected cells [13, 35, 36], which poses a significant challenge for proteomic approaches: the densely phosphorylated SR-rich region alone, functionally implicated in numerous viral processes [34, 49, 63], hosts seven closely spaced Arg residues (Fig. 2A). These arginines restrict the length of tryptic phosphopeptides and decrease the probability of their unambiguous identification and phosphosite assignment [64]. The multiplicity of implicated protein kinases (including GSK-3, protein kinase C, casein kinase II and mitogen-activated protein kinase [36, 37, 50]) further hinders study of specific phosphorylations. Assuming that the mechanistic implication of a specific phosphorylation is independent of the acting kinase, we produced polyphosphorylated N protein (pN) via bacterial co-expression with a catalytic subunit of PKA. A combination of orthogonal cleavage enzymes and LC-MS phosphoproteomics mapped > 20 phosphosites (Supplementary table 1) including Ser23, Thr24, Ser180, Ser194, Ser197, Thr198, Ser201, Ser202, Thr205 and Thr391 reported recently at SARS-CoV-2 infection [13, 35]. At least six of the identified phosphosites are located in the unstructured regions and represent potential 14-3-3-binding sites (Fig. 2A).

Biochemical analysis confirmed that polyphosphorylated N is competent for binding to all seven human 14-3-3 isoforms (Fig. 3 and 4), but revealed remarkable variation in binding efficiency between them (Fig. 4). This was supported by the quantified affinities to two selected isoforms, 14-3-3γ (K_D_ of 1.5 μM) and 14-3-3ε (K_D_ of 10.7 μM) (Fig. 5). Our observations are in line with the recent finding that 14-3-3γ and 14-3-3η systematically bind phosphopeptides with higher affinities than 14-3-3ε and 14-3-3σ [55]. The low micromolar-range K_D_ values compare well to those reported for other physiologically relevant partners of 14-3-3 [65–67], indicating a stable and specific interaction. Meanwhile, the well-defined 2:2 stoichiometry of the ~150-kDa 14-3-3/pN complex, supported by titration experiments and SEC-MALS analysis (Fig. 3 and 5), excludes the possibility that 14-3-3 binding disrupts pN dimerization [57]. It is reasonable to assume that the principally bivalent 14-3-3 dimer [44, 46] recognizes just one phosphosite in each pN subunit because a bidentate 14-3-3 binding to different phosphosites within a single pN subunit would inevitably alter the observed 2:2 stoichiometry. Identification of the single phosphosite responsible for 14-3-3 recruitment proved challenging, as none of the potential sites were a perfect match to the currently known optimal 14-3-3-binding motifs [65].

To restrict the search, we analyzed the interaction of various N constructs with 14-3-3γ. This eliminated the high-scoring potential 14-3-3-binding phosphosite RTApT^265^KAY, present in the C-terminal N fragments, as the true site of interaction despite its conservation in many related coronavirus N proteins (Supplementary table 2 and 3). The residual binding of pN.212-419 to 14-3-3γ suggested the existence of auxiliary 14-3-3-binding sites located outside the folded CTD (residues 247-364). Both pN.1-211 and pN.1-238 bound 14-3-3 with equivalent efficiency suggesting the binding lies between amino acid 1 and 211. Interestingly, the N-terminal constructs existed as monomers (Supplementary Fig. 3), which could potentially lower the binding affinity to 14-3-3 dimers in light of the 2:2 stoichiometry. Nonetheless, sufficient binding was clearly observed between the dimeric 14-3-3γ and the monomeric N-terminal constructs (Fig. 6 and 7).

Truncation of the SR-rich region streamlined the search for the 14-3-3-binding site to the 15-residue stretch of amino acids 196-211. This sequence hosts two principally similar potential 14-3-3-binding sites, RNpS^197^TP and RGpT^205^SP (Fig. 2A and C). Importantly, the proximity of these sites rules out bidentate binding to the 14-3-3 dimer: 14-3-3 binds in an antiparallel manner requiring a minimum of 13-15 residues between phosphosites on a single peptide [44, 68]. Thus, binding to Ser197 and Thr205 sites must be mutually exclusive.

The markedly different binding between pN.1-204 and pN.1-196 to 14-3-3 (Fig. 8) prompted us to propose Ser197 as the critical phosphosite. This finding aided the design of a topology model for the complex (Fig. 9A), in which the 14-3-3 dimer is anchored by two identical Ser197 phosphosites from the SR-rich region in the two equivalent pN chains. Noteworthily, the RN(pS/pT)^197^TP site is conserved in not only SARS-CoV and SARS-CoV-2 but also in N proteins from several bat and pangolin coronaviruses (Fig. 9B). Meanwhile, plausible phosphorylation of residue 205 is possible in a smaller subset of coronaviruses (Fig. 9B).

The model shown in Fig. 9A notably does not exclude the possibility of 14-3-3 binding to alternative phosphosites under significantly different phosphorylation conditions. Moreover, we speculate that hierarchical phosphorylation within the SR-rich region [36] would alter 14-3-3 binding with phosphorylation at adjacent Ser/Thr residues, likely to inhibit the interaction [69]. The SR-rich region is phosphorylated by both Pro-directed and non-Pro-directed protein kinases [36, 37, 39, 50]. Since 14-3-3 proteins typically reject peptides with a proline adjacent to the phosphorylated residue, the interplay between these aforementioned kinases could be regulatory. In theory, this could create a phosphorylation code and conditional binding of 14-3-3, as has been discussed recently for alternative polyphosphorylated 14-3-3 partners such as LRRK2, CFTR and tau protein [48, 67, 70, 71].

The recruitment of the 14-3-3 dimer is expected to occlude the SR-rich region of N by masking 10-20 residues surrounding the Ser197 phosphosite within the complex. Apart from the likely effects on the properties of N and its ability to phase separate and bind RNA, the 14-3-3 binding at the SR-rich region, triggered by its phosphorylation, can potentially interfere with N binding to the M protein, an event clearly relevant to the virion assembly [17]. In support of this hypothesis, the 14-3-3-occluded area reported here overlaps with the N region (residues 168-208) proposed to mediate its association with the M protein in SARS-CoV [19].

The presence of SR-rich regions in many viral N proteins suggests a more broad interaction of 14-3-3 with N proteins. Indeed, using 14-3-3-Pred prediction [58] a number of 14-3-3-binding phosphosites in N proteins from human (Supplementary table 2) and bat coronaviruses (Supplementary table 3) could be identified. Some display strong conservation of the 14-3-3-binding binding site around Ser197 (Fig. 9B), whereas others contain separate high-scoring potential 14-3-3-binding sites beyond the SR-rich region. Given the reasonable threat that other zoonotic coronaviruses may ultimately enter the human population [72, 73], the relevant N proteins are highly likely to undergo phosphorylation and 14-3-3 binding, as seen for SARS-CoV-2 N. This is particularly likely, given the high concentration of both proteins in the infected cells (see above).

Our findings underline the essential role of the SR-rich region in the biology of N proteins and host-virus interactions [49]. Unrelated proteins with similar domains also tend to show RNA-binding capability (e.g., the splicing factors) [74, 75] and are subject to multisite phosphorylation [76]. Such proteins are often associated with phase separation as a means to regulate membraneless compartmentalization within the cytoplasm. Likewise, SARS-CoV-2 N protein has been shown to undergo phase separation in vitro upon RNA addition: a phenomenon dependent on the concentration of salt, presence of divalent ions, phosphorylation state of N and on RNA sequence [30, 34, 77, 78]. Furthermore, the N protein has been shown to recruit the RNA-dependent RNA-polymerase complex, and granule-associated heterogeneous nuclear ribonucleoproteins forming phase separated granules which can aid SARS-CoV-2 replication [30, 34]. Thus 14-3-3 potentially influences phase separation by binding the SR-rich region and may also affect the accessibility of the NES sequence located nearby (Fig. 2A). This in turn may impact nucleocytoplasmic shuttling of N, as seen for SARS-CoV N [37].

Conclusively, 14-3-3 binding to the SR-rich region of N holds potential to regulate multiple cell host processes affected by N. 14-3-3 binding to pN may present a cell immune-like response to the viral infection aimed at arresting or neutralizing N activities [57]. On the other hand, in light of the abundance of N protein in the infected cell [12, 13], pN may instead arrest 14-3-3 proteins in the cytoplasm and indirectly disrupt cellular processes involving 14-3-3. For example, 14-3-3ε and 14-3-3η each play a role in the innate immune response via RIG-1 and MDA5 signaling respectively [79, 80]. The N protein:14-3-3 interaction would modulate these and other signaling pathways involving 14-3-3 proteins. As such, understanding the molecular mechanism of pN association with 14-3-3 proteins may inform the development of novel therapeutic approaches.

## Materials and methods

### Protein expression and purification

The tagless SARS-CoV-2 N gene coding for Uniprot ID P0DTC9 protein sequence was commercially synthesized (GeneWiz) and cloned into pET-SUMO bacterial expression vectors using *Nde*I and *Hind*III restriction endonuclease sites. To obtain constructs of N carrying the N-terminal His_6_-tag cleavable by HRV-3C protease (N.1-419, N.1-211, N.1-238, N.212-419), the N gene was PCR amplified using primers listed in Supplementary table 4 and cloned into the pET-YBSL-Lic-3C plasmid vector using an established ligation-independent cloning procedure [81]. After 3C cleavage each construct contained extra GPA residues at the N terminus. The N.247-364 construct carrying an N-terminal uncleavable His_6_-tag and corresponding to the previously crystallized folded CTD dimer [28], was kindly provided by Prof. W. Baumeister’s laboratory.

Truncated mutants of N.1-211, namely N.1-179, N.1-196 and N.1-204 were derived from the wild-type N plasmid using the standard T7 forward primer with varying reverse primers listed in Supplementary table 4. Each carried a *BamH*I site to enable consequent cloning into a pET28 vector using *Nco*I and *BamH*I sites. This preserved the N-terminal His_6_-tag cleavable by 3C protease. All constructs were validated by DNA sequencing.

Untagged full-length human 14-3-3γ (Uniprot ID P61981) and 14-3-3ζ (Uniprot ID P63104) cloned in a pET21 vector, and full-length human 14-3-3ε (Uniprot ID P62258) carrying an N-terminal His_6_-tag cleavable by Tobacco Etch Virus (TEV) protease, and cloned into a pRSET-A vector have been previously described [82–84]. Truncated versions of human 14-3-3η (Uniprot ID Q04917), 14-3-3β (Uniprot ID P31946), 14-3-3ε (Uniprot ID P62258), 14-3-3τ (Uniprot ID P27348), 14-3-3γ (Uniprot ID P61981) and 14-3-3σ (Uniprot ID P31947) devoid of the short disordered segment at the C terminus, cloned into pProExHtb vector and carrying TEV-cleavable His_6_-tags on their N-terminal ends were obtained as described previously [66, 85]. The monomeric mutant form of untagged full-length human 14-3-3ζ carrying monomerizing amino acid substitutions and pseudophosphorylation in the subunit interface (14-3-3ζm-S58E) was obtained as before [56].

14-3-3γ, 14-3-3ζ and 14-3-3ζm-S58E were expressed and purified using ammonium sulfate fractionation and column chromatography as previously described [82]. His_6_-tagged N.247-364 was also expressed and purified as before [28]. All other proteins carrying cleavable His_6_-tags were expressed in *E.coli* BL21(DE3) cells and purified using subtractive immobilized metal-affinity chromatography (IMAC) and gel-filtration. For the N protein and its various constructs the eluate from the first IMAC column, performed at 1 M NaCl, harbored a significant quantity of random bound nucleic acid (the 260/280 nm absorbance ratio of 1.7-2.0). This necessitated an additional long on-column washing step with 3 M NaCl (50 column volumes) to ensure the 260/280 nm absorbance ratio of ~0.6 and removal of all nucleic acid. Tagless N protein was purified after treatment of cell lysate with RNAse, using heparin and subsequent ion-exchange chromatography on a sulfopropyl Sepharose (elution gradients 100-1000 mM NaCl) followed by gel-filtration. Here, It proved challenging to entirely eliminate nucleic acid and the typical 260/280 nm absorbance ratio was 0.7-0.9.

Protein concentration was determined by spectrophotometry at 280 nm on a N80 Nanophotometer (Implen, Munich, Germany) using sequence-specific extinction coefficients calculated using the ProtParam tool in ExPASy (see Supplementary table 5).

### Isolation of E.coli tRNA

The DH5◻ cells were incubated in 30 ml of liquid medium (LB without antibiotics) for 16 h at 37 °C with maximum aeration and then harvested by centrifugation at 7000 g for 10 min. The pellet was gently resuspended in RNAse-free Tris-acetate buffer, followed by alkali-SDS lysis and neutralization by cold ammonium acetate. The suspension was incubated on ice for 5 min and centrifuged at 21000 g at 4 °C for 10 min. The resulting supernatant (8 ml) was incubated with 12.5 ml isopropanol for 15 min at 25 °C and centrifuged at 12100 g for 5 min. The pellet was resuspended in 800 μl of 2 M ammonium acetate, incubated for 5 min at 25 °C and centrifuged (12100 g, 10 min). RNA was precipitated from supernatant by 800 μl of isopropanol, incubated for 5 min at 25 °C and centrifuged (12100 g, 5 min). After supernatant removal the pellet was washed with 70% ice-cold ethanol, dried and dissolved in 100-200 μl milliQ-water.

### SARS-CoV-2 N protein co-expression with PKA

For phosphorylation in cells, SARS-CoV-2 N was bacterially co-expressed with a catalytic subunit of mouse protein kinase A (PKA), as described previously [48]. PKA was cloned into a low-copy pACYC vector [48] which ensured that the target protein was expressed in excess of kinase.

PKA and SARS-CoV-2 N were co-transformed into *E.coli* BL21(DE3) cells against Kanamycin and Chloramphenicol resistance. Cells were grown in LB to an OD_600_ reading of 0.6 before inducing with 500 μM of IPTG. After induction, cultivation was continued for 1.5 h, 3 h, 4 h and overnight at 37 ◻C, and 4 h was found to be sufficient to provide for the saturating interaction with 14-3-3. For all truncated constructs of N, overnight co-expression was used. For unphosphorylated controls we expressed proteins in the absence of PKA.

The cells with overexpressed proteins were harvested by centrifugation and resuspended in 20 mM Tris-HCl buffer pH 8.0 containing 1 M NaCl, 10 mM imidazole, 0.01 mM phenylmethylsulfonyl fluoride, as well as trace amounts of lysozyme to facilitate lysis. Phosphorylated proteins were purified using subtractive IMAC and gel-filtration, whereby the His_6_-tagged PKA was efficiently removed. Phosphorylated N and its constructs typically showed significant shifts on SDS-PAGE and PhosTag gels indicating phosphorylation.

### Identification of phosphosites within N

#### Sample treatment for proteomics analysis

For phosphopeptide mapping, the SARS-CoV-2 N protein co-expressed with PKA for 4 h at 30 ◻C was purified as above. An aliquot (35 μg) was subjected to enzymatic hydrolysis “in solution” either with trypsin (Sequencing Grade Modified Trypsin, Promega) or with chymotrypsin (Analytical Grade, Sigma-Aldrich). Briefly, the sample preparation was as follows. The sample was reduced with 2 mM Tris(2-carboxyethyl)phosphine (TCEP) and then alkylated with 4 mM S-Methyl methanethiosulfonate (MMTS); a protein:enzyme ratio was kept at 50:1 (w/w); digestion was performed overnight at 37 ◻C and pH 7.8 (for chymotrypsin, 10 mM Ca^2+^ was added to the reaction solution). The reaction was stopped by adjusting it to pH 2 with formic acid. The resulting peptides were purified on a custom micro-tip SPE column with Oasis HLB (Waters) as a stationary phase, using elution with an acetonitrile:water:formic acid mix (50:49.9:0.1 v/v/v%). The eluate of resulting peptides was dried out and stored at −30 ◻C prior to the LC-MS experiment.

#### LC-MS/MS experiment

Peptides were separated on a nano-flow chromatographic system with a flow rate of 440 nl/min (Ultimate 3000 Nano RSLC, Thermo Fisher Scientific) and ESI coupled to a mass-analyzer (Q Exactive Plus, Thermo Fisher Scientific). Briefly, the protocol was as follows. Dried peptide samples were rehydrated with 0.1% formic acid. An aliquot of rehydrated peptides (5 μl) was injected onto a precolumn (100 μm x 2 cm, Reprosil-Pur C18-AQ 1.9 μm, Dr. Maisch GmbH) and eluted on a nano-column (100 μm x 30 cm, Reprosil-Pur C18-AQ 1.9 μm, Dr. Maisch GmbH) with a linear gradient of mobile phase A (water:formic acid 99.9:0.1 v/v%) with mobile phase B (acetonitrile:water:formic acid 80:19.9:0.1 v/v/v%) from 2% till 80% B within 80 min. MS data were acquired by a data-dependent acquisition approach after a MS-scan of a 350−1500 m/z range. Top 12 peaks were subjected to high collision energy dissociation (HCD) or electron transfer dissociation (ETD). After a round of MS/MS, masses were dynamically excluded from further analysis for 35 s.

#### MS data analysis

For Mascot (MatrixScience) database search, raw LC-MS files were converted to general mgf format using MSConvert with its default settings. For peptide database search, a concatenated database including SARS-CoV-2 and general contaminants was constructed. The parameters of the search were: Enzyme – Trypsin with one miscleavage allowed, or in case of Chymotrypsin - no enzyme; MS tolerance – 5 ppm; MSMS – 0.2 Da; Fixed modifications – Methylthio(C); Variable modifications – Phospho(ST), Phospho(Y), Oxidation(M). For PEAKS (PEAKS Studio Xpro, Bioinformatics Solutions Inc.) analysis the parameters of a search were MS tolerance - 10 ppm, MSMS – 0.05 Da; Fixed modifications – Methylthio(C); Variable modifications – Phospho(STY), Oxidation(M), Ammonia-loss (N), Deamidation (NQ). For estimation of FDR, a target-decoy approach was used.

### Analytical size-exclusion chromatography (SEC)

The oligomeric state of proteins as well as protein-protein and protein-tRNA interactions were analyzed by loading 50 μl samples on a Superdex 200 Increase 5/150 column (GE Healthcare) operated at a 0.45 ml/min flow rate using a Varian ProStar 335 system (Varian Inc., Melbourne, Australia). When specified, a Superdex 200 Increase 10/300 column (GE Healthcare) operated at a 0.8 ml/min flow rate was used (100 μl loading). The columns were equilibrated by a 20 mM Tris-HCl buffer, pH 7.6, containing 200 mM NaCl and 3 mM NaN_3_ (SEC buffer) and calibrated by the following protein markers: BSA trimer (198 kDa), BSA dimer (132 kDa), BSA monomer (66 kDa), ovalbumin (43 kDa), α-lactalbumin (15 kDa). The profiles were followed by 280 nm and (optionally) at 260 nm absorbance. Diode array detector data were used to retrieve full-absorbance spectral information about the eluted samples including protein and nucleic acid. All SEC experiments were performed at least three times and the most typical results are presented.

To assess binding parameters for the 14-3-3/pN.1-419 interaction, we used serial loading of the samples containing a fixed pN concentration and increasing concentrations of 14-3-3 in a constant volume of 50 μl and a Superdex 200 Increase 5/150 column operated at 0.45 ml/min. To validate linear augmentation of the peak amplitude with increasing protein concentration, serial loading of 14-3-3 alone at different concentrations was used. Such linear dependence allowed conversion of the amplitude of the 14-3-3 peak into the molar concentration of its unbound form at each point of titration. Concentration of pN-bound 14-3-3 was determined as the difference between the total and free concentration. Binding curves representing the dependence of bound 14-3-3 on its total concentration were approximated using the quadratic equation to determine apparent K_D_ values. Graphing and fitting were performed in Origin 9.0 (OriginLab Corporation, Northampton, MA, USA).

### Size-exclusion chromatography coupled to multi-angle light scattering (SEC-MALS)

To determine the absolute masses of various N constructs and their complexes with 14-3-3, we coupled a SEC column to a ProStar 335 UV/Vis detector (Varian Inc., Melbourne, Australia) and a multi-angle laser light scattering detector miniDAWN (Wyatt Technologies). Either Superdex 200 Increase 10/300 (~24 ml, flow rate 0.8 ml/min) or Superdex 200 Increase 5/150 (~3 ml, flow rate 0.45 ml/min) columns (GE Healthcare) were used. The miniDAWN detector was calibrated relative to the scattering from toluene and, together with concentration estimates obtained from UV detector at 280 nm, was exploited for determining the Mw distribution of the eluted protein species. All processing was performed in ASTRA 8.0 software (Wyatt Technologies) taking dn/dc equal to 0.185 and using extinction coefficients listed in Supplementary table 5. Protein content in the eluted peaks was additionally analyzed by SDS-PAGE.

## Data availability

The source LC-MS data on SARS-CoV-2 N phosphoproteomics are available along with the paper as a Supplementary data file 1.

## Acknowledgements

N.N.S. is thankful to Prof. W. Baumeister and Dr. L. Zinzula (Martinsried, Germany) for providing the N.247-364 plasmid. MALDI and MALS were carried out at the Shared-Access Equipment Centre “Industrial Biotechnology” of the Federal Research Center “Fundamentals of Biotechnology” of the Russian Academy of Sciences. This work was supported by the Russian Science Foundation (grant 19-74-10031 to N.N.S.) and the Wellcome Trust (206377 award to A.A.A.). Protein-protein interactions were studied in the framework of the Program of the Russian Ministry of Science and Higher Education (K.V.T. and N.N.S.). O.I.K. acknowledges support by the Interdisciplinary Scientific and Educational School of Moscow University «Molecular Technologies of the Living Systems and Synthetic Biology». The authors declare no competing interests.

