## Supplementary Tables 1-5 Supplementary Figs 1-3 for "The mechanism of SARS-CoV-2 nucleocapsid protein recognition by the human 14-3-3 proteins"

**Supplementary Table 1.** Predicted and experimentally observed phosphorylation sites within SARS-CoV-2 N shown along with the predicted 14-3-3-binding sites.

| In vivo phosphosites reported by Davidson et al. (10.1186/s13073-020-00763-0) | In vivo phosphosites reported by Bouhaddou et al. (10.1016/j.cell.2020.06.034 ) | NetPhos 3.1 prediction of PKA sites (scores >0.2) | Scansite 4.0 prediction of PKA sites (minimum stringency) | Identified phosphosites (N co-expression with PKA) trypsin+LC-MS York (this work) | Identified phosphosites (N co-expression with PKA) trypsin+LC-MS and chymotrypsin+LC-MS Moscow (this work) | Identified phosphosites (N co-expression with PKA) trypsin+MALDI Moscow (this work) | Scansite 4.0 prediction of 1433 sites (minimum stringency) | 1433-Pred prediction of 1433-binding sites | 1433-Pred consensus score (color coded: from red - unlikely, to green - likely) | Structural context |
|---|---|---|---|---|---|---|---|---|---|---|
| S2 | | S2 | | | | | | | | |
| | | T16 | | T16 | T16 | | | | | |
| | | S21 | | S21 | | | | ITFGGP[S21]DSTG | -0.441 | |
| S23 | S23 | | | | S23 | | S23 | FGGPSD[S23]TGSN | -0.14 | |
| T24 | T24 | | | T24 | T24 | S21, S23 or T24* | | GGPSDS[T24]GSNQ | -0.711 | |
| | S26 | | | S26 | | | | PSDSTG[S26]NQNG | -0.556 | |
| | | S51 | | S51 | | | | LPNNTA[S51]WFTA | -0.248 | |
| T76 | T76 | | | T76 | | | | QGVPIN[T76]NSSP | -0.594 | |
| S78 | | | | | | | S78 | VPINTN[S78]SPDD | 0.359 | structurally blocked |
| S79 | S79 | | | | | | | PINTNS[S79]PDDQ | -0.421 | |
| | | T91 | T91 | | T91 | | T91 | GYYRRA[T91]RRIR | 0.852 | structurally blocked |
| **S105** | **S105** | **S105** | **S105** | | | | | GKMKDL[S105]PRWY | -0.218 | |
| | | | | T135 | | | | GIIWVA[T135]EGAL | 0.365 | structurally blocked |
| T141 | | | | | | | | TEGALN[T141]PKDH | -0.772 | |
| | | | | T165 | | | | LQLPQG[T165]TLPK | -0.63 | |
| T166 | | | | T166 | | | T166 | QLPQGT[T166]LPKG | 0.478 | potentially flexible loop |
| S176 | S176 | | | | | | | GFYAEG[S176]RGGS | -0.396 | |
| **S180** | **S180** | **S180** | **S180** | | **S180** | **S180** | **S180** | EGSRGG[S180]QASS | -0.134 | IDPR |
| S183 | S183 | | | | | | | RGGSQA[S183]SRSS | -0.583 | |
| S184 | S184 | | | | | | | GGSQAS[S184]RSSS | -0.49 | |
| | | | | | S186 | | | SQASSR[S186]SSRS | -0.518 | |
| | | S188 | S188 | | S188 | | S188 | ASSRSS[S188]RSRN | 0.295 | IDPR |
| | | | | | S193 | | | SSRSRN[S193]SRNS | -0.106 | |
| **S194** | **S194** | **S194** | **S194** | | **S194** | | **S194** | SRSRNS[S194]RNST | 0.273 | IDPR |
| | **S197** | **S197** | **S197** | **S197** | **S197** | | **S197** | RNSSRN[S197]TPGS | 0.363 | IDPR |
| **T198** | **T198** | **T198** | **T198** | | T198 | | T198 | NSSRNS[T198]PGSS | 0.03 | |
| S201 | S201 | | | | S201 | | | RNSTPG[S201]SRGT | -0.608 | |
| S202 | S202 | S202 | | | S202 | | | NSTPGS[S202]RGTS | -0.562 | |
| **T205** | **T205** | **T205** | **T205** | **T205** | **T205** | | **T205** | PGSSRG[T205]SPAR | 0.411 | IDPR |
| S206 | S206 | S206 | S206 | | S206 | | | GSSRGT[S206]PARM | -0.016 | |
| | | | | S235 | S235 | | | QLESKM[S235]GKGQ | -0.238 | |
| | | T263 | T263 | T263 | | | | KPRQKR[T263]ATKA | -0.237 | |
| | | T265 | T265 | T265 | T265 | T263 or T265* | T265 | RQKRTA[T265]KAYN | 0.604 | potentially flexible loop |
| | | | | T271 | | | | TKAYNV[T271]QAFG | -0.137 | |
| | | | | T282 | | | | RRGPEQ[T282]QGNF | -0.268 | |
| | | | | T325 | | | | RIGMEV[T325]PSGT | -0.588 | |
| | | | | T379 | | | | KKKADE[T379]QALP | -0.448 | |
| **T391** | | **T391** | **T391** | T391 | T391 | | T391 | RQKKQQ[T391]VTLL | 0.067 | unknown |
| | | | | T393 | T393 | | | KKQQTV[T393]LLPA | 0.172 | |
| | | | | | S404 | | | ADLDDF[S404]KQLQ | -0.385 | |
| | | | | S410 | S410 | | | SKQLQQ[S410]MSSA | -0.418 | |
| | | S413 | | | S413 | | | LQQSMS[S413]ADST | -0.44 | |

Moscow/York designates the place where the experiment was done
**bold black font** indicates the residues that are in vivo phosphorylated within SARS CoV-2 N, predicted to be phosphorylated by PKA, phosphorylated by PKA in E.coli co-expression system, and predicted as 14-3-3 binding sites
**bold blue font** in column names indicates the experimental data.
**bold brown-red font** in column names indicates the predictions or assumptions.
*ambiguity in phosphosite identification within the detected phosphopeptide(s)

**Supplementary Table 2.** Prediction of the 14-3-3-binding sites in all human coronavirus N proteins as the output of 14-3-3-Pred (Madeira et al 2015).

| Sequence | Identifier | Site | Peptide_[-6:4] | ANN | PSSM | SVM | Consensus | Manual inspection remark |
|---|---|---|---|---|---|---|---|---|
| NCAP_SARS Nucleoprotein OS=Severe acute respiratory syndrome coronavirus OX=694009 GN=N PE=1 SV=1 | | | | | | | | higher consesus score indicates more probable 14-3-3 binding |
| 1 | P59595 | 2 | ----M s DNGP | 0.33 | -0.143 | -0.5 | -0.104 |  |
| 1 | P59595 | 8 | SDNGPQ s NQRS | 0.037 | -0.412 | -1.807 | -0.727 | |
| 1 | P59595 | 12 | PQSNQR s APRI | 0.405 | 0.424 | -0.298 | 0.177 | |
| 1 | P59595 | 17 | RSAPRI t FGGP | 0.451 | 0.056 | -0.106 | 0.134 | |
| 1 | P59595 | 22 | ITFGGP t DSTD | 0.132 | -0.211 | -1.084 | -0.388 | |
| 1 | P59595 | 24 | FGGPTD s TDNN | 0.111 | -0.022 | -1.181 | -0.364 | |
| 1 | P59595 | 25 | GGPTDS t DNNQ | 0.135 | -0.148 | -1.322 | -0.445 | |
| 1 | P59595 | 50 | QGLPNN t ASWF | 0.131 | -0.112 | -0.879 | -0.287 | |
| 1 | P59595 | 52 | LPNNTA s WFTA | 0.106 | 0 | -0.849 | -0.248 | |
| 1 | P59595 | 55 | NTASWF t ALTQ | 0.154 | -0.094 | -1.04 | -0.327 | |
| 1 | P59595 | 58 | SWFTAL t QHGK | 0.139 | -0.161 | -1.293 | -0.438 | |
| 1 | P59595 | 77 | QGVPIN t NSGP | 0.07 | -0.295 | -1.268 | -0.498 | |
| 1 | P59595 | 79 | VPINTN s GPDD | 0.534 | 0.518 | 0.078 | 0.377 | |
| 1 | P59595 | 92 | GYYRRA t RRVR | 0.853 | 1.012 | 0.666 | 0.844 | questionable |
| 1 | P59595 | 106 | GKMKEL s PRWY | 0.28 | -0.199 | -0.814 | -0.244 | |
| 1 | P59595 | 116 | YFYYLG t GPEA | 0.36 | 0.343 | -0.117 | 0.195 | |
| 1 | P59595 | 121 | GTGPEA s LPYG | 0.718 | 0.467 | 0 | 0.395 | |
| 1 | P59595 | 136 | GIVWVA t EGAL | 0.638 | 0.329 | 0.073 | 0.347 | questionable |
| 1 | P59595 | 142 | TEGALN t PKDH | 0.051 | -0.518 | -1.848 | -0.772 | |
| 1 | P59595 | 149 | PKDHIG t RNPN | 0.184 | -0.165 | -0.785 | -0.255 | |
| 1 | P59595 | 158 | PNNNAA t VLQL | 0.203 | -0.006 | -0.629 | -0.144 | |
| 1 | P59595 | 166 | LQLPQG t TLPK | 0.035 | -0.37 | -1.556 | -0.63 | |
| 1 | P59595 | 167 | QLPQGT t LPKG | 0.615 | 0.497 | 0.321 | 0.478 | questionable |
| 1 | P59595 | 177 | GFYAEG s RGGS | 0.151 | -0.235 | -1.103 | -0.396 | |
| 1 | P59595 | 181 | EGSRGG s QASS | 0.091 | 0.442 | -0.934 | -0.134 | |
| 1 | P59595 | 184 | RGGSQA s SRSS | 0.057 | -0.25 | -1.556 | -0.583 | |
| 1 | P59595 | 185 | GGSQAS s RSSS | 0.089 | -0.257 | -1.301 | -0.49 | |
| 1 | P59595 | 187 | SQASSR s SSRS | 0.059 | -0.177 | -1.437 | -0.518 | |
| 1 | P59595 | 188 | QASSRS s SRSR | 0.078 | -0.25 | -1.266 | -0.479 | |

| | | | | | | | | |
|---|---|---|---|---|---|---|---|---|
| 1 | P59595 | 189 | **ASSRSS s RSRG** | 0.467 | 0.644 | -0.179 | 0.311 | |
| 1 | P59595 | 191 | **SRSSSR s RGNS** | 0.2 | 0.22 | -1.014 | -0.198 | |
| 1 | P59595 | 195 | **SRSRGN s RNST** | 0.32 | 0.805 | -0.255 | 0.29 | |
| 1 | P59595 | 198 | **RGNSRN s TPGS** | 0.697 | 0.526 | 0.033 | 0.419 | likely (by analogy with SARS2 N) |
| 1 | P59595 | 199 | **GNSRNS t PGSS** | 0.215 | 0.381 | -0.438 | 0.053 | |
| 1 | P59595 | 202 | **RNSTPG s SRGN** | 0.079 | -0.424 | -1.35 | -0.565 | |
| 1 | P59595 | 203 | **NSTPGS s RGNS** | 0.064 | -0.303 | -1.458 | -0.566 | |
| 1 | P59595 | 207 | **GSSRGN s PARM** | 0.317 | 0.338 | -0.567 | 0.029 | |
| 1 | P59595 | 213 | **SPARMA s GGGE** | 0.557 | 0.634 | 0.176 | 0.456 | questionable |
| 1 | P59595 | 218 | **ASGGGE t ALAL** | 0.057 | -0.158 | -1.458 | -0.52 | |
| 1 | P59595 | 233 | **RLNQLE s KVSG** | 0.15 | -0.053 | -1.051 | -0.318 | |
| 1 | P59595 | 236 | **QLESKV s GKGQ** | 0.176 | -0.244 | -0.794 | -0.287 | |
| 1 | P59595 | 246 | **QQQQGQ t VTKK** | 0.118 | -0.14 | -0.816 | -0.279 | |
| 1 | P59595 | 248 | **QQGQTV t KKSA** | 0.092 | -0.151 | -0.76 | -0.273 | |
| 1 | P59595 | 251 | **QTVTKK s AAEA** | 0.174 | -0.112 | -0.967 | -0.302 | |
| 1 | P59595 | 256 | **KSAAEA s KKPR** | 0.152 | -0.249 | -0.909 | -0.335 | |
| 1 | P59595 | 264 | **KPRQKR t ATKQ** | 0.188 | 0.023 | -1.163 | -0.317 | |
| 1 | P59595 | 266 | **RQKRTA t KQYN** | 0.672 | 0.963 | 0.549 | 0.728 | unlikely (by analogy with SARS2 N) |
| 1 | P59595 | 272 | **TKQYNV t QAFG** | 0.329 | -0.068 | -0.616 | -0.118 | |
| 1 | P59595 | 283 | **RRGPEQ t QGNF** | 0.166 | 0.137 | -1.106 | -0.268 | |
| 1 | P59595 | 297 | **DLIRQG t DYKH** | 0.445 | 0.755 | -0.51 | 0.23 | |
| 1 | P59595 | 311 | **IAQFAP s ASAF** | 0.266 | 0.03 | -0.65 | -0.118 | |
| 1 | P59595 | 313 | **QFAPSA s AFFG** | 0.36 | 0.155 | -0.221 | 0.098 | |
| 1 | P59595 | 319 | **SAFFGM s RIGM** | 0.108 | -0.181 | -1.136 | -0.403 | |
| 1 | P59595 | 326 | **RIGMEV t PSGT** | 0.086 | -0.467 | -1.384 | -0.588 | |
| 1 | P59595 | 328 | **GMEVTP s GTWL** | 0.102 | 0.025 | -0.991 | -0.288 | |
| 1 | P59595 | 330 | **EVTPSG t WLTY** | 0.139 | 0.111 | -0.766 | -0.172 | |
| 1 | P59595 | 333 | **PSGTWL t YHGA** | 0.104 | -0.218 | -1.046 | -0.387 | |
| 1 | P59595 | 363 | **HIDAYK t FPPT** | 0.694 | 0.583 | 0.571 | 0.616 | questionable |
| 1 | P59595 | 367 | **YKTFPP t EPKK** | 0.643 | 0.44 | 0.307 | 0.463 | questionable |
| 1 | P59595 | 377 | **KDKKKK t DEAQ** | 0.295 | -0.009 | -0.522 | -0.079 | |
| 1 | P59595 | 392 | **RQKKQP t VTLL** | 0.34 | 0.067 | -0.445 | -0.013 | |
| 1 | P59595 | 394 | **KKQPTV t LLPA** | 0.379 | 0.038 | -0.299 | 0.039 | |

| | | | | | | | |
|---|---|---|---|---|---|---|---|
| 1 | P59595 | 405 | **ADMDDF s RQLQ** | 0.083 | -0.22 | -1.376 | -0.504 |
| 1 | P59595 | 411 | **SRQLQN s MSGA** | 0.138 | 0.195 | -1.176 | -0.281 |
| 1 | P59595 | 413 | **QLQNSM s GASA** | 0.283 | 0.267 | -0.12 | 0.143 |
| 1 | P59595 | 416 | **NSMSGA s ADST** | 0.08 | -0.166 | -1.258 | -0.448 |
| 1 | P59595 | 419 | **SGASAD s TQA-** | 0.194 | -0.088 | -0.858 | -0.251 |
| 1 | P59595 | 420 | **GASADS t QA--** | 0.086 | -0.199 | -1.563 | -0.559 |
| **NCAP_CVHNL Nucleoprotein OS=Human coronavirus NL63 OX=277944 GN=N PE=1 SV=1** | | | | | | | |
| 2 | Q6Q1R8 | 3 | **---MA s VNWA** | 0.343 | -0.232 | -0.364 | -0.084 |
| 2 | Q6Q1R8 | 20 | **KKFPPP s FYMP** | 0.155 | -0.218 | -0.861 | -0.308 |
| 2 | Q6Q1R8 | 28 | **YMPLLV s SDKA** | 0.099 | -0.076 | -1.162 | -0.38 |
| 2 | Q6Q1R8 | 29 | **MPLLVS s DKAP** | 0.147 | -0.05 | -1.01 | -0.304 |
| 2 | Q6Q1R8 | 83 | **HFYYLG t GPHK** | 0.231 | 0.29 | -0.22 | 0.1 |
| 2 | Q6Q1R8 | 95 | **LKFRQR s DGVV** | 0.724 | 1.019 | 0.136 | 0.626 |
| 2 | Q6Q1R8 | 108 | **AKEGAK t VNTS** | 0.524 | 0.12 | 0.04 | 0.228 |
| 2 | Q6Q1R8 | 111 | **GAKTVN t SLGN** | 0.267 | -0.076 | -0.993 | -0.267 |
| 2 | Q6Q1R8 | 112 | **AKTVNT s LGNR** | 0.593 | 0.186 | 0.042 | 0.274 |
| 2 | Q6Q1R8 | 128 | **PLEPKF s IALP** | 0.105 | -0.143 | -0.884 | -0.307 |
| 2 | Q6Q1R8 | 136 | **ALPPEL s VVEF** | 0.091 | -0.165 | -1.263 | -0.446 |
| 2 | Q6Q1R8 | 144 | **VEFEDR s NNSS** | 0.039 | -0.249 | -1.876 | -0.695 |
| 2 | Q6Q1R8 | 147 | **EDRSNN s SRAS** | 0.255 | 0.042 | -0.624 | -0.109 |
| 2 | Q6Q1R8 | 148 | **DRSNNS s RASS** | 0.053 | -0.152 | -1.41 | -0.503 |
| 2 | Q6Q1R8 | 151 | **NNSSRA s SRSS** | 0.118 | -0.195 | -1.126 | -0.401 |
| 2 | Q6Q1R8 | 152 | **NSSRAS s RSST** | 0.226 | 0.511 | -0.487 | 0.083 |
| 2 | Q6Q1R8 | 154 | **SRASSR s STRN** | 0.221 | 0.163 | -0.969 | -0.195 |
| 2 | Q6Q1R8 | 155 | **RASSRS s TRNN** | 0.199 | -0.067 | -1.101 | -0.323 |
| 2 | Q6Q1R8 | 156 | **ASSRSS t RNNS** | 0.566 | 0.822 | 0.111 | 0.5 |
| 2 | Q6Q1R8 | 160 | **SSTRNN s RDSS** | 0.412 | 0.66 | -0.165 | 0.302 |
| 2 | Q6Q1R8 | 163 | **RNNSRD s SRST** | 0.09 | -0.285 | -1.41 | -0.535 |
| 2 | Q6Q1R8 | 164 | **NNSRDS s RSTS** | 0.169 | 0.416 | -0.66 | -0.025 |
| 2 | Q6Q1R8 | 166 | **SRDSSR s TSRQ** | 0.1 | 0.033 | -1.431 | -0.433 |
| 2 | Q6Q1R8 | 167 | **RDSSRS t SRQQ** | 0.116 | -0.268 | -1.228 | -0.46 |

| | | | | | | | |
|---|---|---|---|---|---|---|---|
| 2 | Q6Q1R8 | 168 | **DSSRST s RQQS** | 0.744 | 0.838 | 0.469 | 0.684 |
| 2 | Q6Q1R8 | 172 | **STSRQQ s RTRS** | 0.337 | 0.527 | -0.706 | 0.053 |
| 2 | Q6Q1R8 | 174 | **SRQQSR t RSDS** | 0.247 | 0.281 | -0.717 | -0.063 |
| 2 | Q6Q1R8 | 176 | **QQSRTR s DSNQ** | 0.257 | 0.754 | -0.527 | 0.161 |
| 2 | Q6Q1R8 | 178 | **SRTRSD s NQSS** | 0.552 | 1.052 | 0.095 | 0.566 |
| 2 | Q6Q1R8 | 181 | **RSDSNQ s SSDL** | 0.15 | -0.137 | -0.951 | -0.313 |
| 2 | Q6Q1R8 | 182 | **SDSNQS s SDLV** | 0.031 | -0.296 | -1.835 | -0.7 |
| 2 | Q6Q1R8 | 183 | **DSNQSS s DLVA** | 0.264 | 0.041 | -0.788 | -0.161 |
| 2 | Q6Q1R8 | 190 | **DLVAAV t LALK** | 0.676 | 0.111 | -0.041 | 0.249 |
| 2 | Q6Q1R8 | 202 | **LGFDNQ s KSPS** | 0.101 | -0.18 | -0.977 | -0.352 |
| 2 | Q6Q1R8 | 204 | **FDNQSK s PSSS** | 0.077 | -0.204 | -1.053 | -0.393 |
| 2 | Q6Q1R8 | 206 | **NQSKSP s SSGT** | 0.219 | -0.065 | -0.507 | -0.118 |
| 2 | Q6Q1R8 | 207 | **QSKSPS s SGTS** | 0.076 | -0.338 | -1.176 | -0.479 |
| 2 | Q6Q1R8 | 208 | **SKSPSS s GTST** | 0.124 | -0.12 | -0.937 | -0.311 |
| 2 | Q6Q1R8 | 210 | **SPSSSG t STPK** | 0.099 | -0.251 | -1.351 | -0.501 |
| 2 | Q6Q1R8 | 211 | **PSSSGT s TPKK** | 0.264 | 0.171 | -0.523 | -0.029 |
| 2 | Q6Q1R8 | 212 | **SSSGTS t PKKP** | 0.015 | -0.642 | -2.173 | -0.933 |
| 2 | Q6Q1R8 | 221 | **KPNKPL s QPRA** | 0.356 | 0.147 | -0.725 | -0.074 |
| 2 | Q6Q1R8 | 229 | **PRADKP s QLKK** | 0.094 | -0.058 | -1.344 | -0.436 |
| 2 | Q6Q1R8 | 241 | **RWKRVP t REEN** | 0.705 | 0.835 | 0.194 | 0.578 |
| 2 | Q6Q1R8 | 262 | **NHNMGD s DLVQ** | 0.067 | -0.271 | -1.953 | -0.719 |
| 2 | Q6Q1R8 | 291 | **AALFFD s EVST** | 0.091 | -0.223 | -1.421 | -0.518 |
| 2 | Q6Q1R8 | 294 | **FFDSEV s TDEV** | 0.39 | -0.008 | -0.787 | -0.135 |
| 2 | Q6Q1R8 | 295 | **FDSEVS t DEVG** | 0.073 | -0.121 | -1.455 | -0.501 |
| 2 | Q6Q1R8 | 305 | **GDNVQI t YTYK** | 0.068 | -0.297 | -1.643 | -0.624 |
| 2 | Q6Q1R8 | 307 | **NVQITY t YKML** | 0.078 | 0.061 | -0.892 | -0.251 |
| 2 | Q6Q1R8 | 327 | **KFIEQI s AFTK** | 0.118 | -0.084 | -1.13 | -0.365 |
| 2 | Q6Q1R8 | 330 | **EQISAF t KPSS** | 0.308 | 0.434 | -0.165 | 0.192 |
| 2 | Q6Q1R8 | 333 | **SAFTKP s SIKE** | 0.055 | -0.376 | -1.775 | -0.699 |
| 2 | Q6Q1R8 | 334 | **AFTKPS s IKEM** | 0.168 | -0.193 | -0.612 | -0.212 |
| 2 | Q6Q1R8 | 340 | **SIKEMQ s QSSH** | 0.033 | -0.335 | -1.883 | -0.728 |
| 2 | Q6Q1R8 | 342 | **KEMQSQ s SHVA** | 0.34 | 0.171 | -0.395 | 0.039 |
| 2 | Q6Q1R8 | 343 | **EMQSQS s HVAQ** | 0.062 | -0.001 | -1.279 | -0.406 |

| | | | | | | | | |
|---|---|---|---|---|---|---|---|---|
| 2 | Q6Q1R8 | 349 | **SHVAQN t VLNA** | 0.09 | -0.104 | -1.758 | -0.591 | |
| 2 | Q6Q1R8 | 354 | **NTVLNA s IPES** | 0.578 | 0.426 | -0.123 | 0.294 | |
| 2 | Q6Q1R8 | 358 | **NASIPE s KPLA** | 0.358 | 0.254 | -0.315 | 0.099 | |
| 2 | Q6Q1R8 | 366 | **PLADDD s AIIE** | 0.271 | -0.031 | -0.817 | -0.192 | |
| **NCAP_CVH22 Nucleoprotein OS=Human coronavirus 229E OX=11137 GN=N PE=1 SV=2** | | | | | | | | |
| 3 | P15130 | 3 | **---MA t VKWA** | 0.258 | -0.284 | -0.553 | -0.193 | |
| 3 | P15130 | 10 | **VKWADA s EPQR** | 0.755 | 0.646 | 0.476 | 0.626 | |
| 3 | P15130 | 23 | **QGRIPY s LYSP** | 0.2 | 0.179 | -0.281 | 0.033 | |
| 3 | P15130 | 26 | **IPYSLY s PLLV** | 0.038 | -0.419 | -1.747 | -0.709 | |
| 3 | P15130 | 32 | **SPLLVD s EQPW** | 0.102 | -0.182 | -1.376 | -0.485 | |
| 3 | P15130 | 66 | **VQKRFR t RKGK** | 0.413 | 0.589 | -0.072 | 0.31 | |
| 3 | P15130 | 75 | **GKRVDL s PKLH** | 0.117 | -0.195 | -1.196 | -0.425 | |
| 3 | P15130 | 85 | **HFYYLG t GPHK** | 0.231 | 0.29 | -0.22 | 0.1 | |
| 3 | P15130 | 110 | **AVDGAK t EPTG** | 0.703 | 0.591 | 0.392 | 0.562 | |
| 3 | P15130 | 113 | **GAKTEP t GYGV** | 0.184 | -0.219 | -1.185 | -0.407 | |
| 3 | P15130 | 122 | **GVRRKN s EPEI** | 0.972 | 1.796 | 1.844 | 1.537 | very likely, matches RXXpS/pTXP/G 1433-binding motif |
| 3 | P15130 | 138 | **KLPNGV t VVEE** | 0.19 | -0.101 | -0.825 | -0.245 | |
| 3 | P15130 | 145 | **VVEEPD s RAPS** | 0.04 | -0.503 | -1.821 | -0.761 | |
| 3 | P15130 | 149 | **PDSRAP s RSQS** | 0.365 | 0.555 | -0.229 | 0.23 | |
| 3 | P15130 | 151 | **SRAPSR s QSRS** | 0.101 | 0.057 | -1.341 | -0.394 | |
| 3 | P15130 | 153 | **APSRSQ s RSQS** | 0.529 | 0.769 | 0.098 | 0.465 | |
| 3 | P15130 | 155 | **SRSQSR s QSRG** | 0.18 | 0.12 | -1.098 | -0.266 | |
| 3 | P15130 | 157 | **SQSRSQ s RGRG** | 0.685 | 0.864 | 0.406 | 0.652 | |
| 3 | P15130 | 163 | **SRGRGE s KPQS** | 0.674 | 1.309 | 0.46 | 0.814 | very likely, matches RXXpS/pTXP/G 1433-binding motif |
| 3 | P15130 | 167 | **GESKPQ s RNPS** | 0.199 | -0.202 | -0.885 | -0.296 | |
| 3 | P15130 | 171 | **PQSRNP s SDRN** | 0.598 | 0.7 | 0.213 | 0.504 | |
| 3 | P15130 | 172 | **QSRNPS s DRNH** | 0.123 | -0.132 | -1.11 | -0.373 | |
| 3 | P15130 | 178 | **SDRNHN s QDDI** | 0.517 | 0.264 | -0.304 | 0.159 | |
| 3 | P15130 | 192 | **VAAALK s LGFD** | 0.365 | -0.039 | -0.213 | 0.038 | |

| | | | | | | | | |
|---|---|---|---|---|---|---|---|---|
| 3 | P15130 | 205 | **QEKDKK s AKTG** | 0.217 | -0.027 | -0.779 | -0.196 | |
| 3 | P15130 | 208 | **DKKSAK t GTPK** | 0.579 | -0.025 | -0.152 | 0.134 | |
| 3 | P15130 | 210 | **KSAKTG t PKPS** | 0.041 | -0.525 | -1.581 | -0.688 | |
| 3 | P15130 | 214 | **TGTPKP s RNQS** | 0.08 | -0.358 | -1.112 | -0.463 | |
| 3 | P15130 | 218 | **KPSRNQ s PASS** | 0.141 | 0.327 | -0.695 | -0.076 | |
| 3 | P15130 | 221 | **RNQSPA s SQTS** | 0.124 | -0.212 | -1.072 | -0.387 | |
| 3 | P15130 | 222 | **NQSPAS s QTSA** | 0.078 | -0.272 | -1.276 | -0.49 | |
| 3 | P15130 | 224 | **SPASSQ t SAKS** | 0.084 | -0.116 | -1.191 | -0.408 | |
| 3 | P15130 | 225 | **PASSQT s AKSL** | 0.067 | -0.153 | -1.352 | -0.479 | |
| 3 | P15130 | 228 | **SQTSAK s LARS** | 0.249 | -0.085 | -0.57 | -0.135 | |
| 3 | P15130 | 232 | **AKSLAR s QSSE** | 0.107 | -0.171 | -1.491 | -0.518 | |
| 3 | P15130 | 234 | **SLARSQ s SETK** | 0.708 | 0.991 | 0.649 | 0.783 | |
| 3 | P15130 | 235 | **LARSQS s ETKE** | 0.15 | 0.009 | -1.061 | -0.301 | |
| 3 | P15130 | 237 | **RSQSSE t KEQK** | 0.306 | 0.022 | -0.555 | -0.076 | |
| 3 | P15130 | 258 | **QPNDDV t SNVT** | 0.149 | -0.167 | -1.091 | -0.37 | |
| 3 | P15130 | 259 | **PNDDVT s NVTQ** | 0.108 | -0.247 | -1.403 | -0.514 | |
| 3 | P15130 | 262 | **DVTSNV t QCFG** | 0.156 | -0.306 | -1.25 | -0.467 | |
| 3 | P15130 | 276 | **LDHNFG s AGVV** | 0.062 | -0.17 | -1.568 | -0.559 | |
| 3 | P15130 | 298 | **FAELVP s TAAM** | 0.182 | -0.107 | -0.891 | -0.272 | |
| 3 | P15130 | 299 | **AELVPS t AAML** | 0.045 | -0.32 | -1.673 | -0.649 | |
| 3 | P15130 | 306 | **AAMLFD s HIVS** | 0.099 | -0.176 | -1.403 | -0.493 | |
| 3 | P15130 | 310 | **FDSHIV s KESG** | 0.124 | -0.152 | -0.819 | -0.282 | |
| 3 | P15130 | 313 | **HIVSKE s GNTV** | 0.099 | -0.112 | -1.23 | -0.414 | |
| 3 | P15130 | 316 | **SKESGN t VVLT** | 0.146 | -0.128 | -1.043 | -0.342 | |
| 3 | P15130 | 320 | **GNTVVL t FTTR** | 0.197 | -0.107 | -0.804 | -0.238 | |
| 3 | P15130 | 322 | **TVVLTF t TRVT** | 0.069 | -0.093 | -1.14 | -0.388 | |
| 3 | P15130 | 323 | **VVLTFT t RVTV** | 0.1 | -0.225 | -1.425 | -0.517 | |
| 3 | P15130 | 326 | **TFTTRV t VPKD** | 0.818 | 0.624 | 0.65 | 0.697 | likely |
| 3 | P15130 | 345 | **EELNAF t REMQ** | 0.098 | -0.173 | -1.122 | -0.399 | |
| 3 | P15130 | 357 | **HPLLNP s ALEF** | 0.118 | -0.077 | -1.042 | -0.334 | |
| 3 | P15130 | 364 | **ALEFNP s QTSP** | 0.165 | -0.209 | -1.146 | -0.397 | |
| 3 | P15130 | 366 | **EFNPSQ t SPAT** | 0.6 | 0.62 | 0.455 | 0.558 | |
| 3 | P15130 | 367 | **FNPSQT s PATA** | 0.029 | -0.574 | -1.778 | -0.774 | |

| | | | | | | | | |
|---|---|---|---|---|---|---|---|---|
| 3 | P15130 | 370 | **SQTSPA t AEPV** | 0.084 | -0.241 | -1.432 | -0.53 | |
| 3 | P15130 | 379 | **PVRDEV s IETD** | 0.464 | 0.165 | -0.383 | 0.082 | |
| 3 | P15130 | 382 | **DEVSIE t DIID** | 0.2 | -0.127 | -1.076 | -0.334 | |
| **NCAP_SARS2 Nucleoprotein OS=Severe acute respiratory syndrome coronavirus 2 OX=2697049 GN=N PE=1 SV=1** | | | | | | | | |
| 4 | P0DTC9 | 2 | **----M s DNGP** | 0.33 | -0.143 | -0.5 | -0.104 |  |
| 4 | P0DTC9 | 16 | **RNAPRI t FGGP** | 0.413 | 0.031 | -0.125 | 0.106 | |
| 4 | P0DTC9 | 21 | **ITFGGP s DSTG** | 0.114 | -0.232 | -1.206 | -0.441 | |
| 4 | P0DTC9 | 23 | **FGGPSD s TGSN** | 0.194 | 0.089 | -0.704 | -0.14 | |
| 4 | P0DTC9 | 24 | **GGPSDS t GSNQ** | 0.045 | -0.369 | -1.809 | -0.711 | |
| 4 | P0DTC9 | 26 | **PSDSTG s NQNG** | 0.075 | -0.202 | -1.54 | -0.556 | |
| 4 | P0DTC9 | 33 | **NQNGER s GARS** | 0.042 | -0.341 | -1.774 | -0.691 | |
| 4 | P0DTC9 | 37 | **ERSGAR s KQRR** | 0.219 | 0.14 | -0.665 | -0.102 | |
| 4 | P0DTC9 | 49 | **QGLPNN t ASWF** | 0.131 | -0.112 | -0.879 | -0.287 | |
| 4 | P0DTC9 | 51 | **LPNNTA s WFTA** | 0.106 | 0 | -0.849 | -0.248 | |
| 4 | P0DTC9 | 54 | **NTASWF t ALTQ** | 0.154 | -0.094 | -1.04 | -0.327 | |
| 4 | P0DTC9 | 57 | **SWFTAL t QHGK** | 0.139 | -0.161 | -1.293 | -0.438 | |
| 4 | P0DTC9 | 76 | **QGVPIN t NSSP** | 0.045 | -0.299 | -1.527 | -0.594 | |
| 4 | P0DTC9 | 78 | **VPINTN s SPDD** | 0.539 | 0.531 | 0.008 | 0.359 | |
| 4 | P0DTC9 | 79 | **PINTNS s PDDQ** | 0.133 | -0.282 | -1.115 | -0.421 | |
| 4 | P0DTC9 | 91 | **GYYRRA t RRIR** | 0.873 | 0.98 | 0.702 | 0.852 | unlikely |
| 4 | P0DTC9 | 105 | **GKMKDL s PRWY** | 0.268 | -0.197 | -0.726 | -0.218 | |
| 4 | P0DTC9 | 115 | **YFYYLG t GPEA** | 0.36 | 0.343 | -0.117 | 0.195 | |
| 4 | P0DTC9 | 135 | **GIIWVA t EGAL** | 0.649 | 0.356 | 0.089 | 0.365 | unlikely |
| 4 | P0DTC9 | 141 | **TEGALN t PKDH** | 0.051 | -0.518 | -1.848 | -0.772 | |
| 4 | P0DTC9 | 148 | **PKDHIG t RNPA** | 0.18 | -0.208 | -0.713 | -0.247 | |
| 4 | P0DTC9 | 165 | **LQLPQG t TLPK** | 0.035 | -0.37 | -1.556 | -0.63 | |
| 4 | P0DTC9 | 166 | **QLPQGT t LPKG** | 0.615 | 0.497 | 0.321 | 0.478 | unlikely |
| 4 | P0DTC9 | 176 | **GFYAEG s RGGS** | 0.151 | -0.235 | -1.103 | -0.396 | |
| 4 | P0DTC9 | 180 | **EGSRGG s QASS** | 0.091 | 0.442 | -0.934 | -0.134 | |
| 4 | P0DTC9 | 183 | **RGGSQA s SRSS** | 0.057 | -0.25 | -1.556 | -0.583 | |
| 4 | P0DTC9 | 184 | **GGSQAS s RSSS** | 0.089 | -0.257 | -1.301 | -0.49 | |

| | | | | | | | | |
|---|---|---|---|---|---|---|---|---|
| 4 | P0DTC9 | 186 | **SQASSR s SSRS** | 0.059 | -0.177 | -1.437 | -0.518 | |
| 4 | P0DTC9 | 187 | **QASSRS s SRSR** | 0.078 | -0.25 | -1.266 | -0.479 | |
| 4 | P0DTC9 | 188 | **ASSRSS s RSRN** | 0.458 | 0.659 | -0.232 | 0.295 | unlikely |
| 4 | P0DTC9 | 190 | **SRSSSR s RNSS** | 0.106 | 0.111 | -1.133 | -0.305 | |
| 4 | P0DTC9 | 193 | **SSRSRN s SRNS** | 0.364 | 0.152 | -0.835 | -0.106 | |
| 4 | P0DTC9 | 194 | **SRSRNS s RNST** | 0.277 | 0.751 | -0.208 | 0.273 | |
| 4 | P0DTC9 | 197 | **RNSSRN s TPGS** | 0.648 | 0.444 | -0.003 | 0.363 | most likely (this study) |
| 4 | P0DTC9 | 198 | **NSSRNS t PGSS** | 0.195 | 0.366 | -0.47 | 0.03 | |
| 4 | P0DTC9 | 201 | **RNSTPG s SRGT** | 0.059 | -0.495 | -1.388 | -0.608 | |
| 4 | P0DTC9 | 202 | **NSTPGS s RGTS** | 0.058 | -0.37 | -1.375 | -0.562 | |
| 4 | P0DTC9 | 205 | **PGSSRG t SPAR** | 0.597 | 0.49 | 0.145 | 0.411 | likely (this study) |
| 4 | P0DTC9 | 206 | **GSSRGT s PARM** | 0.239 | 0.287 | -0.575 | -0.016 | |
| 4 | P0DTC9 | 232 | **RLNQLE s KMSG** | 0.177 | -0.048 | -1.007 | -0.293 | |
| 4 | P0DTC9 | 235 | **QLESKM s GKGQ** | 0.157 | -0.118 | -0.754 | -0.238 | |
| 4 | P0DTC9 | 245 | **QQQQGQ t VTKK** | 0.118 | -0.14 | -0.816 | -0.279 | |
| 4 | P0DTC9 | 247 | **QQGQTV t KKSA** | 0.092 | -0.151 | -0.76 | -0.273 | |
| 4 | P0DTC9 | 250 | **QTVTKK s AAEA** | 0.174 | -0.112 | -0.967 | -0.302 | |
| 4 | P0DTC9 | 255 | **KSAAEA s KKPR** | 0.152 | -0.249 | -0.909 | -0.335 | |
| 4 | P0DTC9 | 263 | **KPRQKR t ATKA** | 0.219 | 0.048 | -0.977 | -0.237 | |
| 4 | P0DTC9 | 265 | **RQKRTA t KAYN** | 0.582 | 0.892 | 0.339 | 0.604 | unlikely (this study) |
| 4 | P0DTC9 | 271 | **TKAYNV t QAFG** | 0.295 | -0.126 | -0.581 | -0.137 | |
| 4 | P0DTC9 | 282 | **RRGPEQ t QGNF** | 0.166 | 0.137 | -1.106 | -0.268 | |
| 4 | P0DTC9 | 296 | **ELIRQG t DYKH** | 0.325 | 0.753 | -0.495 | 0.194 | |
| 4 | P0DTC9 | 310 | **IAQFAP s ASAF** | 0.266 | 0.03 | -0.65 | -0.118 | |
| 4 | P0DTC9 | 312 | **QFAPSA s AFFG** | 0.36 | 0.155 | -0.221 | 0.098 | |
| 4 | P0DTC9 | 318 | **SAFFGM s RIGM** | 0.108 | -0.181 | -1.136 | -0.403 | |
| 4 | P0DTC9 | 325 | **RIGMEV t PSGT** | 0.086 | -0.467 | -1.384 | -0.588 | |
| 4 | P0DTC9 | 327 | **GMEVTP s GTWL** | 0.102 | 0.025 | -0.991 | -0.288 | |
| 4 | P0DTC9 | 329 | **EVTPSG t WLTY** | 0.139 | 0.111 | -0.766 | -0.172 | |
| 4 | P0DTC9 | 332 | **PSGTWL t YTGA** | 0.105 | -0.254 | -1.079 | -0.409 | |
| 4 | P0DTC9 | 334 | **GTWLTY t GAIK** | 0.129 | 0.044 | -0.948 | -0.258 | |
| 4 | P0DTC9 | 362 | **HIDAYK t FPPT** | 0.694 | 0.583 | 0.571 | 0.616 | unlikely (this study) |
| 4 | P0DTC9 | 366 | **YKTFPP t EPKK** | 0.643 | 0.44 | 0.307 | 0.463 | unlikely (this study) |

| | | | | | | | |
|---|---|---|---|---|---|---|---|
| 4 | P0DTC9 | 379 | **KKKADE t QALP** | 0.145 | -0.235 | -1.253 | -0.448 |
| 4 | P0DTC9 | 391 | **RQKKQQ t VTLL** | 0.37 | 0.144 | -0.312 | 0.067 |
| 4 | P0DTC9 | 393 | **KKQQTV t LLPA** | 0.492 | 0.099 | -0.074 | 0.172 |
| 4 | P0DTC9 | 404 | **ADLDDF s KQLQ** | 0.113 | -0.181 | -1.087 | -0.385 |
| 4 | P0DTC9 | 410 | **SKQLQQ s MSSA** | 0.076 | 0.008 | -1.337 | -0.418 |
| 4 | P0DTC9 | 412 | **QLQQSM s SADS** | 0.6 | 0.402 | 0.166 | 0.389 |
| 4 | P0DTC9 | 413 | **LQQSMS s ADST** | 0.062 | -0.211 | -1.171 | -0.44 |
| 4 | P0DTC9 | 416 | **SMSSAD s TQA-** | 0.15 | 0.005 | -1.099 | -0.315 |
| 4 | P0DTC9 | 417 | **MSSADS t QA--** | 0.084 | -0.169 | -1.435 | -0.507 |
| **NCAP_CVHOC Nucleoprotein OS=Human coronavirus OC43 OX=31631 GN=N PE=1 SV=1** | | | | | | | |
| 5 | P33469 | 2 | **----M s FTPG** | 0.45 | -0.09 | -0.334 | 0.009 |
| 5 | P33469 | 4 | **--MSF t PGKQ** | 0.216 | -0.31 | -0.743 | -0.279 |
| 5 | P33469 | 9 | **FTPGKQ s SSRA** | 0.058 | -0.286 | -1.53 | -0.586 |
| 5 | P33469 | 10 | **TPGKQS s SRAS** | 0.075 | -0.214 | -1.336 | -0.492 |
| 5 | P33469 | 11 | **PGKQSS s RASS** | 0.149 | -0.055 | -0.702 | -0.203 |
| 5 | P33469 | 14 | **QSSSRA s SGNR** | 0.334 | 0.041 | -0.67 | -0.098 |
| 5 | P33469 | 15 | **SSSRAS s GNRS** | 0.364 | 0.57 | -0.279 | 0.218 |
| 5 | P33469 | 19 | **ASSGNR s GNGI** | 0.161 | -0.197 | -0.697 | -0.244 |
| 5 | P33469 | 30 | **LKWADQ s DQVR** | 0.339 | 0.178 | -0.38 | 0.046 |
| 5 | P33469 | 38 | **QVRNVQ t RGRR** | 0.315 | 0.097 | -0.482 | -0.023 |
| 5 | P33469 | 48 | **RAQPKQ t ATSQ** | 0.093 | -0.153 | -1.371 | -0.477 |
| 5 | P33469 | 50 | **QPKQTA t SQQP** | 0.157 | -0.103 | -0.768 | -0.238 |
| 5 | P33469 | 51 | **PKQTAT s QQPS** | 0.393 | 0.054 | -0.57 | -0.041 |
| 5 | P33469 | 55 | **ATSQQP s GGNV** | 0.172 | -0.159 | -1.279 | -0.422 |
| 5 | P33469 | 64 | **NVVPYY s WFSG** | 0.104 | 0.062 | -0.779 | -0.204 |
| 5 | P33469 | 67 | **PYYSWF s GITQ** | 0.151 | -0.072 | -0.738 | -0.22 |
| 5 | P33469 | 70 | **SWFSGI t QFQK** | 0.087 | -0.29 | -1.416 | -0.54 |
| 5 | P33469 | 95 | **APGVPA t EAKG** | 0.068 | -0.216 | -1.397 | -0.515 |
| 5 | P33469 | 108 | **YRHNRG s FKTA** | 0.222 | 0.224 | -0.402 | 0.015 |
| 5 | P33469 | 111 | **NRGSFK t ADGN** | 0.303 | 0.209 | -0.534 | -0.007 |
| 5 | P33469 | 130 | **YFYYLG t GPHA** | 0.233 | 0.262 | -0.217 | 0.093 |
| 5 | P33469 | 140 | **AKDQYG t DIDG** | 0.409 | 0.014 | -0.486 | -0.021 |

| | | | | | | | | |
|---|---|---|---|---|---|---|---|---|
| 5 | P33469 | 150 | **GVYWVA s NQAD** | 0.415 | 0.082 | -0.391 | 0.035 | |
| 5 | P33469 | 157 | **NQADVN t PADI** | 0.223 | -0.225 | -0.791 | -0.264 | |
| 5 | P33469 | 167 | **IVDRDP s SDEA** | 0.571 | 0.705 | 0.215 | 0.497 | |
| 5 | P33469 | 168 | **VDRDPS s DEAI** | 0.253 | 0.093 | -0.584 | -0.079 | |
| 5 | P33469 | 174 | **SDEAIP t RFPP** | 0.041 | -0.499 | -1.531 | -0.663 | |
| 5 | P33469 | 180 | **TRFPPG t VLPQ** | 0.057 | -0.166 | -1.391 | -0.5 | |
| 5 | P33469 | 191 | **GYYIEG s GRSA** | 0.081 | -0.246 | -1.355 | -0.507 | |
| 5 | P33469 | 194 | **IEGSGR s APNS** | 0.219 | 0.319 | -1.009 | -0.157 | |
| 5 | P33469 | 198 | **GRSAPN s RSTS** | 0.149 | -0.069 | -1.112 | -0.344 | |
| 5 | P33469 | 200 | **SAPNSR s TSRT** | 0.052 | -0.218 | -1.565 | -0.577 | |
| 5 | P33469 | 201 | **APNSRS t SRTS** | 0.092 | -0.261 | -1.454 | -0.541 | |
| 5 | P33469 | 202 | **PNSRST s RTSS** | 0.562 | 0.797 | 0.235 | 0.531 | |
| 5 | P33469 | 204 | **SRSTSR t SSRA** | 0.139 | 0.076 | -1.2 | -0.328 | |
| 5 | P33469 | 205 | **RSTSRT s SRAS** | 0.283 | -0.008 | -0.545 | -0.09 | |
| 5 | P33469 | 206 | **STSRTS s RASS** | 0.105 | 0.471 | -1.018 | -0.147 | |
| 5 | P33469 | 209 | **RTSSRA s SAGS** | 0.248 | -0.087 | -0.949 | -0.263 | |
| 5 | P33469 | 210 | **TSSRAS s AGSR** | 0.498 | 0.776 | 0.08 | 0.451 | |
| 5 | P33469 | 213 | **RASSAG s RSRA** | 0.093 | -0.389 | -1.574 | -0.623 | |
| 5 | P33469 | 215 | **SSAGSR s RANS** | 0.089 | -0.088 | -1.327 | -0.442 | |
| 5 | P33469 | 219 | **SRSRAN s GNRT** | 0.688 | 0.955 | 0.308 | 0.65 | |
| 5 | P33469 | 223 | **ANSGNR t PTSG** | 0.036 | -0.561 | -1.729 | -0.751 | |
| 5 | P33469 | 225 | **SGNRTP t SGVT** | 0.306 | 0.702 | -0.252 | 0.252 | |
| 5 | P33469 | 226 | **GNRTPT s GVTP** | 0.172 | -0.081 | -0.792 | -0.234 | |
| 5 | P33469 | 229 | **TPTSGV t PDMA** | 0.069 | -0.556 | -1.382 | -0.623 | |
| 5 | P33469 | 238 | **MADQIA s LVLA** | 0.444 | 0.101 | -0.102 | 0.148 | |
| 5 | P33469 | 249 | **KLGKDA t KPQQ** | 0.724 | 0.48 | 0.226 | 0.477 | likely, matches RXXpS/pTXP/G 1433-binding motif |
| 5 | P33469 | 255 | **TKPQQV t KHTA** | 0.348 | -0.083 | -0.492 | -0.076 | |
| 5 | P33469 | 258 | **QQVTKH t AKEV** | 0.148 | 0.02 | -1.125 | -0.319 | |
| 5 | P33469 | 275 | **KPRQKR s PNKQ** | 0.087 | -0.264 | -1.34 | -0.506 | |
| 5 | P33469 | 281 | **SPNKQC t VQQC** | 0.171 | -0.161 | -1.397 | -0.462 | |
| 5 | P33469 | 305 | **EMLKLG t SDPQ** | 0.077 | -0.125 | -1.269 | -0.439 | |
| 5 | P33469 | 306 | **MLKLGT s DPQF** | 0.73 | 0.606 | 0.367 | 0.568 | |

| | | | | | | | |
|---|---|---|---|---|---|---|---|
| 5 | P33469 | 319 | **LAELAP t AGAF** | 0.301 | 0.041 | -0.504 | -0.054 |
| 5 | P33469 | 327 | **GAFFFG s RLEL** | 0.143 | -0.155 | -1.39 | -0.467 |
| 5 | P33469 | 338 | **AKVQNL s GNPD** | 0.274 | -0.092 | -0.381 | -0.066 |
| 5 | P33469 | 361 | **GAIRFD s TLSG** | 0.355 | 0.667 | -0.564 | 0.153 |
| 5 | P33469 | 362 | **AIRFDS t LSGF** | 0.302 | 0.139 | -0.43 | 0.004 |
| 5 | P33469 | 364 | **RFDSTL s GFET** | 0.082 | -0.148 | -1.05 | -0.372 |
| 5 | P33469 | 368 | **TLSGFE t IMKV** | 0.123 | -0.14 | -1.343 | -0.453 |
| 5 | P33469 | 390 | **DGMMNM s PKPQ** | 0.107 | -0.357 | -1.079 | -0.443 |
| 5 | P33469 | 410 | **GENDNI s VAVP** | 0.157 | -0.107 | -1.178 | -0.376 |
| 5 | P33469 | 416 | **SVAVPK s RVQQ** | 0.06 | -0.3 | -1.217 | -0.486 |
| 5 | P33469 | 423 | **RVQQNK s RELT** | 0.258 | 0.005 | -0.442 | -0.06 |
| 5 | P33469 | 427 | **NKSREL t AEDI** | 0.783 | 0.959 | 0.533 | 0.758 |
| 5 | P33469 | 432 | **LTAEDI s LLKK** | 0.134 | -0.143 | -0.882 | -0.297 |
| 5 | P33469 | 442 | **KMDEPY t EDTS** | 0.08 | 0.037 | -0.992 | -0.292 |
| 5 | P33469 | 445 | **EPYTED t SEI-** | 0.067 | -0.269 | -1.684 | -0.629 |
| 5 | P33469 | 446 | **PYTEDT s EI--** | 0.176 | 0.186 | -0.613 | -0.084 |
| **NCAP_MERS1 Nucleoprotein OS=Middle East respiratory syndrome-related coronavirus (isolate United Kingdom/H123990006/2012) OX=1263720 GN=N PE=1 SV=1** | | | | | | | |
| 6 | K9N4V7 | 3 | **---MA s PAAP** | 0.096 | -0.613 | -1.209 | -0.575 |
| 6 | K9N4V7 | 11 | **AAPRAV s FADN** | 0.833 | 0.885 | 0.575 | 0.764 |
| 6 | K9N4V7 | 19 | **ADNNDI t NTNL** | 0.094 | -0.209 | -1.436 | -0.517 |
| 6 | K9N4V7 | 21 | **NNDITN t NLSR** | 0.053 | -0.168 | -1.566 | -0.56 |
| 6 | K9N4V7 | 24 | **ITNTNL s RGRG** | 0.206 | -0.255 | -1.132 | -0.394 |
| 6 | K9N4V7 | 40 | **RAAPNN t VSWY** | 0.25 | -0.041 | -0.686 | -0.159 |
| 6 | K9N4V7 | 42 | **APNNTV s WYTG** | 0.133 | -0.029 | -0.962 | -0.286 |
| 6 | K9N4V7 | 45 | **NTVSWY t GLTQ** | 0.098 | -0.152 | -1.132 | -0.395 |
| 6 | K9N4V7 | 48 | **SWYTGL t QHGK** | 0.048 | -0.328 | -1.723 | -0.668 |
| 6 | K9N4V7 | 56 | **HGKVPL t FPPG** | 0.394 | 0.409 | 0.149 | 0.317 |
| 6 | K9N4V7 | 69 | **VPLNAN s TPAQ** | 0.637 | 0.508 | 0.177 | 0.441 |
| 6 | K9N4V7 | 70 | **PLNANS t PAQN** | 0.102 | -0.387 | -1.204 | -0.496 |
| 6 | K9N4V7 | 87 | **QDRKIN t GNGI** | 0.567 | 0.203 | 0.211 | 0.327 |
| 6 | K9N4V7 | 103 | **RWYFYY t GTGP** | 0.199 | -0.018 | -0.67 | -0.163 |

| | | | | | | | | |
|---|---|---|---|---|---|---|---|---|
| 6 | K9N4V7 | 105 | **YFYYTG t GPEA** | 0.364 | 0.421 | 0.014 | 0.266 | |
| 6 | K9N4V7 | 129 | **VHEDGA t DAPS** | 0.095 | -0.231 | -1.645 | -0.594 | |
| 6 | K9N4V7 | 133 | **GATDAP s TFGT** | 0.255 | -0.114 | -0.697 | -0.185 | |
| 6 | K9N4V7 | 134 | **ATDAPS t FGTR** | 0.197 | -0.208 | -0.823 | -0.278 | |
| 6 | K9N4V7 | 137 | **APSTFG t RNPN** | 0.082 | -0.365 | -1.604 | -0.629 | |
| 6 | K9N4V7 | 144 | **RNPNND s AIVT** | 0.075 | -0.217 | -1.183 | -0.442 | |
| 6 | K9N4V7 | 148 | **NDSAIV t QFAP** | 0.156 | -0.216 | -0.844 | -0.301 | |
| 6 | K9N4V7 | 154 | **TQFAPG t KLPK** | 0.051 | -0.41 | -1.351 | -0.57 | |
| 6 | K9N4V7 | 165 | **NFHIEG t GGNS** | 0.084 | -0.146 | -1.39 | -0.484 | |
| 6 | K9N4V7 | 169 | **EGTGGN s QSSS** | 0.035 | -0.277 | -1.652 | -0.631 | |
| 6 | K9N4V7 | 171 | **TGGNSQ s SSRA** | 0.109 | -0.087 | -0.838 | -0.272 | |
| 6 | K9N4V7 | 172 | **GGNSQS s SRAS** | 0.078 | -0.229 | -1.387 | -0.513 | |
| 6 | K9N4V7 | 173 | **GNSQSS s RASS** | 0.104 | -0.172 | -1.076 | -0.381 | |
| 6 | K9N4V7 | 176 | **QSSSRA s SVSR** | 0.139 | -0.099 | -1.023 | -0.328 | |
| 6 | K9N4V7 | 177 | **SSSRAS s VSRN** | 0.343 | 0.593 | -0.446 | 0.163 | |
| 6 | K9N4V7 | 179 | **SRASSV s RNSS** | 0.237 | 0.168 | -0.57 | -0.055 | |
| 6 | K9N4V7 | 182 | **SSVSRN s SRSS** | 0.098 | -0.19 | -1.425 | -0.506 | |
| 6 | K9N4V7 | 183 | **SVSRNS s RSSS** | 0.102 | 0.46 | -0.885 | -0.108 | |
| 6 | K9N4V7 | 185 | **SRNSSR s SSQG** | 0.126 | 0.134 | -1.131 | -0.29 | |
| 6 | K9N4V7 | 186 | **RNSSRS s SQGS** | 0.126 | -0.186 | -1.044 | -0.368 | |
| 6 | K9N4V7 | 187 | **NSSRSS s QGSR** | 0.562 | 0.862 | 0.255 | 0.56 | very likely, matches RXXpS/pTXP/G 1433-binding motif |
| 6 | K9N4V7 | 190 | **RSSSQG s RSGN** | 0.054 | -0.418 | -1.801 | -0.722 | |
| 6 | K9N4V7 | 192 | **SSQGSR s GNST** | 0.091 | -0.024 | -0.98 | -0.304 | |
| 6 | K9N4V7 | 195 | **GSRSGN s TRGT** | 0.288 | -0.055 | -0.707 | -0.158 | |
| 6 | K9N4V7 | 196 | **SRSGNS t RGTS** | 0.094 | -0.068 | -1.147 | -0.374 | |
| 6 | K9N4V7 | 199 | **GNSTRG t SPGP** | 0.414 | 0.289 | -0.369 | 0.111 | |
| 6 | K9N4V7 | 200 | **NSTRGT s PGPS** | 0.248 | 0.4 | -0.265 | 0.128 | |
| 6 | K9N4V7 | 204 | **GTSPGP s GIGA** | 0.082 | -0.357 | -1.099 | -0.458 | |
| 6 | K9N4V7 | 227 | **RLQALE s GKVK** | 0.2 | -0.065 | -0.965 | -0.277 | |
| 6 | K9N4V7 | 233 | **SGKVKQ s QPKV** | 0.221 | 0.272 | -0.884 | -0.13 | |
| 6 | K9N4V7 | 239 | **SQPKVI t KKDA** | 0.231 | -0.116 | -0.65 | -0.178 | |
| 6 | K9N4V7 | 255 | **KMRHKR t STKS** | 0.1 | 0.081 | -1.325 | -0.381 | |

| | | | | | | | | |
|---|---|---|---|---|---|---|---|---|
| 6 | K9N4V7 | 256 | **MRHKRT s TKSF** | 0.292 | 0.335 | -0.406 | 0.074 | |
| 6 | K9N4V7 | 257 | **RHKRTS t KSFN** | 0.272 | 0.745 | -0.414 | 0.201 | |
| 6 | K9N4V7 | 259 | **KRTSTK s FNMV** | 0.329 | 0.243 | -0.395 | 0.059 | |
| 6 | K9N4V7 | 288 | **QLNKLG t EDPR** | 0.273 | 0.026 | -0.656 | -0.119 | |
| 6 | K9N4V7 | 302 | **IAELAP t ASAF** | 0.209 | -0.07 | -0.79 | -0.217 | |
| 6 | K9N4V7 | 304 | **ELAPTA s AFMG** | 0.221 | 0.053 | -0.646 | -0.124 | |
| 6 | K9N4V7 | 310 | **SAFMGM s QFKL** | 0.09 | -0.134 | -1.339 | -0.461 | |
| 6 | K9N4V7 | 315 | **MSQFKL t HQNN** | 0.216 | 0.057 | -0.904 | -0.21 | |
| 6 | K9N4V7 | 332 | **VYFLRY s GAIK** | 0.308 | 0.061 | -0.541 | -0.057 | |
| 6 | K9N4V7 | 360 | **NIDAYK t FPKK** | 0.745 | 0.558 | 0.464 | 0.589 | |
| 6 | K9N4V7 | 375 | **KAPKEE s TDQM** | 0.173 | -0.228 | -1.141 | -0.399 | |
| 6 | K9N4V7 | 376 | **APKEES t DQMS** | 0.084 | -0.252 | -1.592 | -0.587 | |
| 6 | K9N4V7 | 380 | **ESTDQM s EPPK** | 0.48 | 0.497 | -0.069 | 0.303 | |
| 6 | K9N4V7 | 391 | **EHRVQG t QRTR** | 0.092 | -0.068 | -1.535 | -0.504 | |
| 6 | K9N4V7 | 394 | **VQGTQR t RTRP** | 0.041 | -0.408 | -1.989 | -0.785 | |
| 6 | K9N4V7 | 396 | **GTQRTR t RPSV** | 0.761 | 1.302 | 0.241 | 0.768 | likely |
| 6 | K9N4V7 | 399 | **RTRTRP s VQPG** | 0.609 | 0.28 | -0.177 | 0.237 | |
| 6 | K9N4V7 | 410 | **PMIDVN t D---** | 0.326 | 0.356 | -0.874 | -0.064 | |
| **NCAP_CVHN1 Nucleoprotein OS=Human coronavirus HKU1 (isolate N1) OX=443239 GN=N PE=3 SV=1** | | | | | | | | |
| 7 | Q5MQC6 | 2 | **----M s YTPG** | 0.27 | -0.215 | -0.704 | -0.216 |  |
| 7 | Q5MQC6 | 4 | **--MSY t PGHY** | 0.29 | -0.219 | -0.396 | -0.108 | |
| 7 | Q5MQC6 | 11 | **PGHYAG s RSSS** | 0.048 | -0.268 | -1.513 | -0.578 | |
| 7 | Q5MQC6 | 13 | **HYAGSR s SSGN** | 0.188 | 0.143 | -0.56 | -0.076 | |
| 7 | Q5MQC6 | 14 | **YAGSRS s SGNR** | 0.151 | -0.017 | -1.004 | -0.29 | |
| 7 | Q5MQC6 | 15 | **AGSRSS s GNRS** | 0.576 | 0.782 | 0.283 | 0.547 | |
| 7 | Q5MQC6 | 19 | **SSSGNR s GILK** | 0.05 | -0.299 | -1.33 | -0.526 | |
| 7 | Q5MQC6 | 25 | **SGILKK t SWAD** | 0.22 | 0.026 | -0.669 | -0.141 | |
| 7 | Q5MQC6 | 26 | **GILKKT s WADQ** | 0.359 | 0.129 | -0.484 | 0.001 | |
| 7 | Q5MQC6 | 31 | **TSWADQ s ERNY** | 0.193 | 0.023 | -0.885 | -0.223 | |
| 7 | Q5MQC6 | 37 | **SERNYQ t FNRG** | 0.428 | 0.186 | -0.343 | 0.09 | |
| 7 | Q5MQC6 | 44 | **FNRGRK t QPKF** | 0.779 | 0.817 | 0.576 | 0.724 | |
| 7 | Q5MQC6 | 49 | **KTQPKF t VSTQ** | 0.079 | -0.224 | -1.327 | -0.491 | |

| | | | | | | | | |
|---|---|---|---|---|---|---|---|---|
| 7 | Q5MQC6 | 51 | **QPKFTV s TQPQ** | 0.194 | -0.103 | -0.791 | -0.233 | |
| 7 | Q5MQC6 | 52 | **PKFTVS t QPQG** | 0.658 | 0.412 | -0.127 | 0.314 | |
| 7 | Q5MQC6 | 58 | **TQPQGN t IPHY** | 0.291 | 0.276 | -0.269 | 0.099 | |
| 7 | Q5MQC6 | 63 | **NTIPHY s WFSG** | 0.164 | 0.1 | -0.753 | -0.163 | |
| 7 | Q5MQC6 | 66 | **PHYSWF s GITQ** | 0.093 | -0.132 | -1.132 | -0.39 | |
| 7 | Q5MQC6 | 69 | **SWFSGI t QFQK** | 0.087 | -0.29 | -1.416 | -0.54 | |
| 7 | Q5MQC6 | 80 | **GRDFKF s DGQG** | 0.431 | 0.283 | -0.39 | 0.108 | |
| 7 | Q5MQC6 | 94 | **AFGVPP s EAKG** | 0.063 | -0.246 | -1.256 | -0.48 | |
| 7 | Q5MQC6 | 104 | **GYWYRH s RRSF** | 0.332 | 0.272 | -0.456 | 0.049 | |
| 7 | Q5MQC6 | 107 | **YRHSRR s FKTA** | 0.246 | 0.238 | -0.551 | -0.022 | |
| 7 | Q5MQC6 | 110 | **SRRSFK t ADGQ** | 0.453 | 0.35 | -0.378 | 0.142 | |
| 7 | Q5MQC6 | 129 | **YFYYLG t GPYA** | 0.282 | 0.286 | -0.299 | 0.09 | |
| 7 | Q5MQC6 | 136 | **GPYANA s YGES** | 0.295 | -0.042 | -0.705 | -0.151 | |
| 7 | Q5MQC6 | 140 | **NASYGE s LEGV** | 0.163 | -0.149 | -0.988 | -0.325 | |
| 7 | Q5MQC6 | 154 | **ANHQAD t STPS** | 0.109 | -0.341 | -1.365 | -0.532 | |
| 7 | Q5MQC6 | 155 | **NHQADT s TPSD** | 0.405 | 0.4 | -0.387 | 0.139 | |
| 7 | Q5MQC6 | 156 | **HQADTS t PSDV** | 0.039 | -0.318 | -1.521 | -0.6 | |
| 7 | Q5MQC6 | 158 | **ADTSTP s DVSS** | 0.038 | -0.292 | -1.694 | -0.649 | |
| 7 | Q5MQC6 | 161 | **STPSDV s SRDP** | 0.109 | -0.372 | -1.423 | -0.562 | |
| 7 | Q5MQC6 | 162 | **TPSDVS s RDPT** | 0.073 | -0.362 | -1.529 | -0.606 | |
| 7 | Q5MQC6 | 166 | **VSSRDP t TQEA** | 0.507 | 0.603 | 0.077 | 0.396 | |
| 7 | Q5MQC6 | 167 | **SSRDPT t QEAI** | 0.274 | 0.083 | -0.639 | -0.094 | |
| 7 | Q5MQC6 | 173 | **TQEAIP t RFPP** | 0.066 | -0.393 | -1.099 | -0.475 | |
| 7 | Q5MQC6 | 179 | **TRFPPG t ILPQ** | 0.044 | -0.227 | -1.483 | -0.555 | |
| 7 | Q5MQC6 | 190 | **GYYVEG s GRSA** | 0.067 | -0.287 | -1.475 | -0.565 | |
| 7 | Q5MQC6 | 193 | **VEGSGR s ASNS** | 0.028 | -0.277 | -2.184 | -0.811 | |
| 7 | Q5MQC6 | 195 | **GSGRSA s NSRP** | 0.581 | 0.811 | -0.056 | 0.445 | |
| 7 | Q5MQC6 | 197 | **GRSASN s RPGS** | 0.815 | 0.783 | 0.47 | 0.689 | |
| 7 | Q5MQC6 | 201 | **SNSRPG s RSQS** | 0.117 | 0.299 | -0.864 | -0.149 | |
| 7 | Q5MQC6 | 203 | **SRPGSR s QSRG** | 0.097 | 0.08 | -1.422 | -0.415 | |
| 7 | Q5MQC6 | 205 | **PGSRSQ s RGPN** | 0.782 | 0.979 | 0.793 | 0.851 | very likely, matches RXXpS/pTXP/G 1433-binding motif |
| 7 | Q5MQC6 | 212 | **RGPNNR s LSRS** | 0.126 | -0.167 | -1.071 | -0.371 | |

| | | | | | | | | |
|---|---|---|---|---|---|---|---|---|
| 7 | Q5MQC6 | 214 | **PNNRSL s RSNS** | 0.405 | 0.804 | -0.188 | 0.34 | |
| 7 | Q5MQC6 | 216 | **NRSLSR s NSNF** | 0.167 | 0.276 | -1.028 | -0.195 | |
| 7 | Q5MQC6 | 218 | **SLSRSN s NFRH** | 0.758 | 0.899 | 0.166 | 0.608 | |
| 7 | Q5MQC6 | 223 | **NSNFRH s DSIV** | 0.436 | 0.171 | -0.836 | -0.076 | |
| 7 | Q5MQC6 | 225 | **NFRHSD s IVKP** | 0.319 | 0.222 | -0.339 | 0.067 | |
| 7 | Q5MQC6 | 247 | **AKLGKD s KPQQ** | 0.505 | 0.284 | -0.144 | 0.215 | |
| 7 | Q5MQC6 | 253 | **SKPQQV t KQNA** | 0.282 | -0.052 | -0.72 | -0.163 | |
| 7 | Q5MQC6 | 266 | **IRHKIL t KPRQ** | 0.575 | 0.525 | 0.063 | 0.388 | |
| 7 | Q5MQC6 | 273 | **KPRQKR t PNKH** | 0.068 | -0.335 | -1.522 | -0.596 | |
| 7 | Q5MQC6 | 290 | **FGKRGP s QNFG** | 0.497 | 0.718 | 0.089 | 0.435 | |
| 7 | Q5MQC6 | 303 | **EMLKLG t NDPQ** | 0.084 | -0.127 | -1.334 | -0.459 | |
| 7 | Q5MQC6 | 317 | **LAELAP t PGAF** | 0.143 | -0.291 | -0.857 | -0.335 | |
| 7 | Q5MQC6 | 325 | **GAFFFG s KLDL** | 0.221 | -0.033 | -0.985 | -0.266 | |
| 7 | Q5MQC6 | 334 | **DLVKRD s EADS** | 0.659 | 0.234 | -0.285 | 0.203 | |
| 7 | Q5MQC6 | 338 | **RDSEAD s PVKD** | 0.062 | -0.445 | -1.658 | -0.68 | |
| 7 | Q5MQC6 | 349 | **VFELHY s GSIR** | 0.208 | -0.042 | -0.615 | -0.15 | |
| 7 | Q5MQC6 | 351 | **ELHYSG s IRFD** | 0.186 | 0.03 | -0.502 | -0.095 | |
| 7 | Q5MQC6 | 356 | **GSIRFD s TLPG** | 0.435 | 0.644 | -0.394 | 0.228 | |
| 7 | Q5MQC6 | 357 | **SIRFDS t LPGF** | 0.688 | 0.679 | 0.393 | 0.587 | very likely, matches RXXXpS/pTXP/G 1433-binding motif |
| 7 | Q5MQC6 | 363 | **TLPGFE t IMKV** | 0.091 | -0.17 | -1.504 | -0.528 | |
| 7 | Q5MQC6 | 378 | **LNAYVN s NQNT** | 0.185 | -0.089 | -0.884 | -0.263 | |
| 7 | Q5MQC6 | 382 | **VNSNQN t DSDS** | 0.109 | -0.154 | -1.213 | -0.419 | |
| 7 | Q5MQC6 | 384 | **SNQNTD s DSLS** | 0.042 | -0.225 | -1.662 | -0.615 | |
| 7 | Q5MQC6 | 386 | **QNTDSD s LSSK** | 0.181 | -0.048 | -0.691 | -0.186 | |
| 7 | Q5MQC6 | 388 | **TDSDSL s SKPQ** | 0.066 | -0.178 | -1.258 | -0.457 | |
| 7 | Q5MQC6 | 389 | **DSDSLS s KPQR** | 0.315 | 0.157 | -0.579 | -0.036 | |
| 7 | Q5MQC6 | 406 | **LPEQFD s LNLS** | 0.285 | -0.08 | -0.558 | -0.118 | |
| 7 | Q5MQC6 | 410 | **FDSLNL s AGTQ** | 0.128 | -0.113 | -1.117 | -0.367 | |
| 7 | Q5MQC6 | 413 | **LNLSAG t QHIS** | 0.113 | -0.262 | -1.305 | -0.485 | |
| 7 | Q5MQC6 | 417 | **AGTQHI s NDFT** | 0.208 | -0.155 | -0.755 | -0.234 | |
| 7 | Q5MQC6 | 421 | **HISNDF t PEDH** | 0.073 | -0.292 | -1.147 | -0.455 | |
| 7 | Q5MQC6 | 426 | **FTPEDH s LLAT** | 0.213 | 0.119 | -0.692 | -0.12 | |

| | | | | | | | |
|---|---|---|---|---|---|---|---|
| 7 | Q5MQC6 | 430 | **DHSLLA t LDDP** | 0.375 | 0.032 | -0.821 | -0.138 |
| 7 | Q5MQC6 | 439 | **DPYVED s VA--** | 0.072 | -0.157 | -1.67 | -0.585 |
| **NCAP_CVHN2 Nucleoprotein OS=Human coronavirus HKU1 (isolate N2) OX=443240 GN=N PE=3 SV=1** | | | | | | | |
| 8 | Q14EA6 | 2 | **----M s YTPG** | 0.27 | -0.215 | -0.704 | -0.216 |
| 8 | Q14EA6 | 4 | **--MSY t PGHH** | 0.153 | -0.377 | -0.762 | -0.329 |
| 8 | Q14EA6 | 11 | **PGHHAG s RSSS** | 0.047 | -0.327 | -1.659 | -0.646 |
| 8 | Q14EA6 | 13 | **HHAGSR s SSGN** | 0.122 | 0.083 | -0.954 | -0.25 |
| 8 | Q14EA6 | 14 | **HAGSRS s SGNR** | 0.157 | 0.036 | -0.935 | -0.247 |
| 8 | Q14EA6 | 15 | **AGSRSS s GNRS** | 0.576 | 0.782 | 0.283 | 0.547 |
| 8 | Q14EA6 | 19 | **SSSGNR s GILK** | 0.05 | -0.299 | -1.33 | -0.526 |
| 8 | Q14EA6 | 25 | **SGILKK t SWVD** | 0.177 | -0.017 | -0.894 | -0.245 |
| 8 | Q14EA6 | 26 | **GILKKT s WVDQ** | 0.375 | 0.185 | -0.411 | 0.05 |
| 8 | Q14EA6 | 31 | **TSWVDQ s ERSH** | 0.039 | -0.219 | -1.527 | -0.569 |
| 8 | Q14EA6 | 34 | **VDQSER s HQTY** | 0.13 | -0.193 | -1.341 | -0.468 |
| 8 | Q14EA6 | 37 | **SERSHQ t YNRG** | 0.288 | 0.047 | -0.851 | -0.172 |
| 8 | Q14EA6 | 49 | **KPQPKF t VSTQ** | 0.064 | -0.243 | -1.392 | -0.524 |
| 8 | Q14EA6 | 51 | **QPKFTV s TQPQ** | 0.194 | -0.103 | -0.791 | -0.233 |
| 8 | Q14EA6 | 52 | **PKFTVS t QPQG** | 0.658 | 0.412 | -0.127 | 0.314 |
| 8 | Q14EA6 | 63 | **NPIPHY s WFSG** | 0.126 | 0.081 | -0.818 | -0.204 |
| 8 | Q14EA6 | 66 | **PHYSWF s GITQ** | 0.093 | -0.132 | -1.132 | -0.39 |
| 8 | Q14EA6 | 69 | **SWFSGI t QFQK** | 0.087 | -0.29 | -1.416 | -0.54 |
| 8 | Q14EA6 | 94 | **AYGIPP s EAKG** | 0.091 | -0.185 | -1.202 | -0.432 |
| 8 | Q14EA6 | 107 | **YKHNRR s FKTA** | 0.324 | 0.144 | -0.28 | 0.063 |
| 8 | Q14EA6 | 110 | **NRRSFK t ADGQ** | 0.561 | 0.414 | -0.099 | 0.292 |
| 8 | Q14EA6 | 129 | **YFYYLG t GPYA** | 0.282 | 0.286 | -0.299 | 0.09 |
| 8 | Q14EA6 | 134 | **GTGPYA s SSYG** | 0.138 | -0.18 | -1.302 | -0.448 |
| 8 | Q14EA6 | 135 | **TGPYAS s SYGD** | 0.121 | -0.156 | -0.839 | -0.291 |
| 8 | Q14EA6 | 136 | **GPYASS s YGDA** | 0.364 | 0.118 | -0.298 | 0.061 |
| 8 | Q14EA6 | 149 | **GIFWVA s HQAD** | 0.68 | 0.23 | 0.061 | 0.324 |
| 8 | Q14EA6 | 154 | **ASHQAD t SIPS** | 0.092 | -0.304 | -1.31 | -0.507 |
| 8 | Q14EA6 | 155 | **SHQADT s IPSD** | 0.267 | 0.31 | -0.645 | -0.023 |

| | | | | | | | | |
|---|---|---|---|---|---|---|---|---|
| 8 | Q14EA6 | 158 | **ADTSIP s DVSA** | 0.062 | -0.318 | -1.323 | -0.526 | |
| 8 | Q14EA6 | 161 | **SIPSDV s ARDP** | 0.164 | -0.191 | -1.184 | -0.404 | |
| 8 | Q14EA6 | 166 | **VSARDP t IQEA** | 0.488 | 0.608 | 0.235 | 0.444 | |
| 8 | Q14EA6 | 173 | **IQEAIP t RFSP** | 0.07 | -0.357 | -1.033 | -0.44 | |
| 8 | Q14EA6 | 176 | **AIPTRF s PGTI** | 0.275 | -0.09 | -0.653 | -0.156 | |
| 8 | Q14EA6 | 179 | **TRFSPG t ILPQ** | 0.046 | -0.232 | -1.54 | -0.575 | |
| 8 | Q14EA6 | 190 | **GYYVEG s GRSA** | 0.067 | -0.287 | -1.475 | -0.565 | |
| 8 | Q14EA6 | 193 | **VEGSGR s ASNS** | 0.028 | -0.277 | -2.184 | -0.811 | |
| 8 | Q14EA6 | 195 | **GSGRSA s NSRP** | 0.581 | 0.811 | -0.056 | 0.445 | |
| 8 | Q14EA6 | 197 | **GRSASN s RPGS** | 0.815 | 0.783 | 0.47 | 0.689 | |
| 8 | Q14EA6 | 201 | **SNSRPG s RSQS** | 0.117 | 0.299 | -0.864 | -0.149 | |
| 8 | Q14EA6 | 203 | **SRPGSR s QSRG** | 0.097 | 0.08 | -1.422 | -0.415 | |
| 8 | Q14EA6 | 205 | **PGSRSQ s RGPN** | 0.782 | 0.979 | 0.793 | 0.851 | very likely, matches RXXpS/pTXP/G 1433-binding motif |
| 8 | Q14EA6 | 212 | **RGPNNR s LSRS** | 0.126 | -0.167 | -1.071 | -0.371 | |
| 8 | Q14EA6 | 214 | **PNNRSL s RSNS** | 0.405 | 0.804 | -0.188 | 0.34 | |
| 8 | Q14EA6 | 216 | **NRSLSR s NSNF** | 0.167 | 0.276 | -1.028 | -0.195 | |
| 8 | Q14EA6 | 218 | **SLSRSN s NFRH** | 0.758 | 0.899 | 0.166 | 0.608 | |
| 8 | Q14EA6 | 223 | **NSNFRH s DSIV** | 0.436 | 0.171 | -0.836 | -0.076 | |
| 8 | Q14EA6 | 225 | **NFRHSD s IVKP** | 0.319 | 0.222 | -0.339 | 0.067 | |
| 8 | Q14EA6 | 237 | **MADEIA s LVLA** | 0.278 | 0.071 | -0.464 | -0.038 | |
| 8 | Q14EA6 | 247 | **AKLGKD s KPQQ** | 0.505 | 0.284 | -0.144 | 0.215 | |
| 8 | Q14EA6 | 253 | **SKPQQV t KQNA** | 0.282 | -0.052 | -0.72 | -0.163 | |
| 8 | Q14EA6 | 273 | **KPRQKR t PNKF** | 0.079 | -0.239 | -1.26 | -0.473 | |
| 8 | Q14EA6 | 296 | **LQNFGN s EMLK** | 0.157 | -0.019 | -0.868 | -0.243 | |
| 8 | Q14EA6 | 303 | **EMLKLG t NDPQ** | 0.084 | -0.127 | -1.334 | -0.459 | |
| 8 | Q14EA6 | 317 | **LAELAP t PGAF** | 0.143 | -0.291 | -0.857 | -0.335 | |
| 8 | Q14EA6 | 325 | **GAFFFG s KLEL** | 0.175 | -0.075 | -1.132 | -0.344 | |
| 8 | Q14EA6 | 334 | **ELFKRD s DADS** | 0.589 | 0.21 | -0.339 | 0.153 | |
| 8 | Q14EA6 | 338 | **RDSDAD s PSKD** | 0.066 | -0.475 | -1.689 | -0.699 | |
| 8 | Q14EA6 | 340 | **SDADSP s KDTF** | 0.16 | -0.041 | -0.704 | -0.195 | |
| 8 | Q14EA6 | 343 | **DSPSKD t FELR** | 0.112 | -0.276 | -1.359 | -0.508 | |
| 8 | Q14EA6 | 349 | **TFELRY s GSIR** | 0.273 | 0.065 | -0.422 | -0.028 | |

| | | | | | | | | |
|---|---|---|---|---|---|---|---|---|
| 8 | Q14EA6 | 351 | **ELRYSG s IRFD** | 0.488 | 0.322 | 0.106 | 0.305 | |
| 8 | Q14EA6 | 356 | **GSIRFD s TLPG** | 0.435 | 0.644 | -0.394 | 0.228 | |
| 8 | Q14EA6 | 357 | **SIRFDS t LPGF** | 0.688 | 0.679 | 0.393 | 0.587 | very likely, matches RXXXpS/pTXP/G 1433-binding motif |
| 8 | Q14EA6 | 363 | **TLPGFE t IMKV** | 0.091 | -0.17 | -1.504 | -0.528 | |
| 8 | Q14EA6 | 378 | **LDAYVN s NQNT** | 0.151 | -0.097 | -1.006 | -0.317 | |
| 8 | Q14EA6 | 382 | **VNSNQN t VSGS** | 0.061 | -0.28 | -1.312 | -0.51 | |
| 8 | Q14EA6 | 384 | **SNQNTV s GSLS** | 0.05 | -0.223 | -1.313 | -0.495 | |
| 8 | Q14EA6 | 386 | **QNTVSG s LSPK** | 0.131 | -0.094 | -0.683 | -0.215 | |
| 8 | Q14EA6 | 388 | **TVSGSL s PKPQ** | 0.045 | -0.343 | -1.285 | -0.528 | |
| 8 | Q14EA6 | 400 | **KRGVKQ s PESF** | 0.02 | -0.311 | -1.875 | -0.722 | |
| 8 | Q14EA6 | 403 | **VKQSPE s FDSL** | 0.231 | -0.019 | -0.605 | -0.131 | |
| 8 | Q14EA6 | 406 | **SPESFD s LNLS** | 0.163 | -0.199 | -1.126 | -0.387 | |
| 8 | Q14EA6 | 410 | **FDSLNL s ADTQ** | 0.116 | -0.122 | -1.151 | -0.386 | |
| 8 | Q14EA6 | 413 | **LNLSAD t QHIS** | 0.147 | -0.263 | -1.267 | -0.461 | |
| 8 | Q14EA6 | 417 | **ADTQHI s NDFT** | 0.142 | -0.23 | -1.01 | -0.366 | |
| 8 | Q14EA6 | 421 | **HISNDF t PEDH** | 0.073 | -0.292 | -1.147 | -0.455 | |
| 8 | Q14EA6 | 426 | **FTPEDH s LLAT** | 0.213 | 0.119 | -0.692 | -0.12 | |
| 8 | Q14EA6 | 430 | **DHSLLA t LDDP** | 0.375 | 0.032 | -0.821 | -0.138 | |
| 8 | Q14EA6 | 439 | **DPYVED s VA--** | 0.072 | -0.157 | -1.67 | -0.585 | |
| **NCAP_CVHN5 Nucleoprotein OS=Human coronavirus HKU1 (isolate N5) OX=443241 GN=N PE=3 SV=1** | | | | | | | | |
| 9 | Q0ZME3 | 2 | **----M s YTPG** | 0.27 | -0.215 | -0.704 | -0.216 |  |
| 9 | Q0ZME3 | 4 | **--MSY t PGHH** | 0.153 | -0.377 | -0.762 | -0.329 | |
| 9 | Q0ZME3 | 11 | **PGHHAG s RSSS** | 0.047 | -0.327 | -1.659 | -0.646 | |
| 9 | Q0ZME3 | 13 | **HHAGSR s SSGN** | 0.122 | 0.083 | -0.954 | -0.25 | |
| 9 | Q0ZME3 | 14 | **HAGSRS s SGNR** | 0.157 | 0.036 | -0.935 | -0.247 | |
| 9 | Q0ZME3 | 15 | **AGSRSS s GNRS** | 0.576 | 0.782 | 0.283 | 0.547 | |
| 9 | Q0ZME3 | 19 | **SSSGNR s GILK** | 0.05 | -0.299 | -1.33 | -0.526 | |
| 9 | Q0ZME3 | 25 | **SGILKK t SWVD** | 0.177 | -0.017 | -0.894 | -0.245 | |
| 9 | Q0ZME3 | 26 | **GILKKT s WVDQ** | 0.375 | 0.185 | -0.411 | 0.05 | |
| 9 | Q0ZME3 | 31 | **TSWVDQ s ERSH** | 0.039 | -0.219 | -1.527 | -0.569 | |

| | | | | | | | | |
|---|---|---|---|---|---|---|---|---|
| 9 | Q0ZME3 | 34 | **VDQSER s HQTY** | 0.13 | -0.193 | -1.341 | -0.468 | |
| 9 | Q0ZME3 | 37 | **SERSHQ t YNRG** | 0.288 | 0.047 | -0.851 | -0.172 | |
| 9 | Q0ZME3 | 49 | **KPQPKF t VSTQ** | 0.064 | -0.243 | -1.392 | -0.524 | |
| 9 | Q0ZME3 | 51 | **QPKFTV s TQPQ** | 0.194 | -0.103 | -0.791 | -0.233 | |
| 9 | Q0ZME3 | 52 | **PKFTVS t QPQG** | 0.658 | 0.412 | -0.127 | 0.314 | |
| 9 | Q0ZME3 | 63 | **NPIPHY s WFSG** | 0.126 | 0.081 | -0.818 | -0.204 | |
| 9 | Q0ZME3 | 66 | **PHYSWF s GITQ** | 0.093 | -0.132 | -1.132 | -0.39 | |
| 9 | Q0ZME3 | 69 | **SWFSGI t QFQK** | 0.087 | -0.29 | -1.416 | -0.54 | |
| 9 | Q0ZME3 | 94 | **AYGIPP s EAKG** | 0.091 | -0.185 | -1.202 | -0.432 | |
| 9 | Q0ZME3 | 107 | **YKHNRR s FKTA** | 0.324 | 0.144 | -0.28 | 0.063 | |
| 9 | Q0ZME3 | 110 | **NRRSFK t ADGQ** | 0.561 | 0.414 | -0.099 | 0.292 | |
| 9 | Q0ZME3 | 129 | **YFYYLG t GPYA** | 0.282 | 0.286 | -0.299 | 0.09 | |
| 9 | Q0ZME3 | 134 | **GTGPYA s SSYG** | 0.138 | -0.18 | -1.302 | -0.448 | |
| 9 | Q0ZME3 | 135 | **TGPYAS s SYGD** | 0.121 | -0.156 | -0.839 | -0.291 | |
| 9 | Q0ZME3 | 136 | **GPYASS s YGDA** | 0.364 | 0.118 | -0.298 | 0.061 | |
| 9 | Q0ZME3 | 149 | **GIFWVA s HQAD** | 0.68 | 0.23 | 0.061 | 0.324 | |
| 9 | Q0ZME3 | 154 | **ASHQAD t SIPS** | 0.092 | -0.304 | -1.31 | -0.507 | |
| 9 | Q0ZME3 | 155 | **SHQADT s IPSD** | 0.267 | 0.31 | -0.645 | -0.023 | |
| 9 | Q0ZME3 | 158 | **ADTSIP s DVSA** | 0.062 | -0.318 | -1.323 | -0.526 | |
| 9 | Q0ZME3 | 161 | **SIPSDV s ARDP** | 0.164 | -0.191 | -1.184 | -0.404 | |
| 9 | Q0ZME3 | 166 | **VSARDP t IQEA** | 0.488 | 0.608 | 0.235 | 0.444 | |
| 9 | Q0ZME3 | 173 | **IQEAIP t RFSP** | 0.07 | -0.357 | -1.033 | -0.44 | |
| 9 | Q0ZME3 | 176 | **AIPTRF s PGTI** | 0.275 | -0.09 | -0.653 | -0.156 | |
| 9 | Q0ZME3 | 179 | **TRFSPG t ILPQ** | 0.046 | -0.232 | -1.54 | -0.575 | |
| 9 | Q0ZME3 | 190 | **GYYVEG s GRSA** | 0.067 | -0.287 | -1.475 | -0.565 | |
| 9 | Q0ZME3 | 193 | **VEGSGR s ASNS** | 0.028 | -0.277 | -2.184 | -0.811 | |
| 9 | Q0ZME3 | 195 | **GSGRSA s NSRP** | 0.581 | 0.811 | -0.056 | 0.445 | |
| 9 | Q0ZME3 | 197 | **GRSASN s RPGS** | 0.815 | 0.783 | 0.47 | 0.689 | |
| 9 | Q0ZME3 | 201 | **SNSRPG s RSQS** | 0.117 | 0.299 | -0.864 | -0.149 | |
| 9 | Q0ZME3 | 203 | **SRPGSR s QSRG** | 0.097 | 0.08 | -1.422 | -0.415 | |
| 9 | Q0ZME3 | 205 | **PGSRSQ s RGPN** | 0.782 | 0.979 | 0.793 | 0.851 | very likely, matches RXXpS/pTXP/G 1433-binding motif |
| 9 | Q0ZME3 | 212 | **RGPNNR s LSRS** | 0.126 | -0.167 | -1.071 | -0.371 | |

| | | | | | | | | |
|---|---|---|---|---|---|---|---|---|
| 9 | Q0ZME3 | 214 | **PNNRSL s RSNS** | 0.405 | 0.804 | -0.188 | 0.34 | |
| 9 | Q0ZME3 | 216 | **NRSLSR s NSNF** | 0.167 | 0.276 | -1.028 | -0.195 | |
| 9 | Q0ZME3 | 218 | **SLSRSN s NFRH** | 0.758 | 0.899 | 0.166 | 0.608 | |
| 9 | Q0ZME3 | 223 | **NSNFRH s DSIV** | 0.436 | 0.171 | -0.836 | -0.076 | |
| 9 | Q0ZME3 | 225 | **NFRHSD s IVKP** | 0.319 | 0.222 | -0.339 | 0.067 | |
| 9 | Q0ZME3 | 237 | **MADEIA s LVLA** | 0.278 | 0.071 | -0.464 | -0.038 | |
| 9 | Q0ZME3 | 247 | **AKLGKD s KPQQ** | 0.505 | 0.284 | -0.144 | 0.215 | |
| 9 | Q0ZME3 | 253 | **SKPQQV t KQNA** | 0.282 | -0.052 | -0.72 | -0.163 | |
| 9 | Q0ZME3 | 273 | **KPRQKR t PNKF** | 0.079 | -0.239 | -1.26 | -0.473 | |
| 9 | Q0ZME3 | 296 | **LQNFGN s EMLK** | 0.157 | -0.019 | -0.868 | -0.243 | |
| 9 | Q0ZME3 | 303 | **EMLKLG t NDPQ** | 0.084 | -0.127 | -1.334 | -0.459 | |
| 9 | Q0ZME3 | 317 | **LAELAP t PGAF** | 0.143 | -0.291 | -0.857 | -0.335 | |
| 9 | Q0ZME3 | 325 | **GAFFFG s KLEL** | 0.175 | -0.075 | -1.132 | -0.344 | |
| 9 | Q0ZME3 | 334 | **ELFKRD s DADS** | 0.589 | 0.21 | -0.339 | 0.153 | |
| 9 | Q0ZME3 | 338 | **RDSDAD s PSKD** | 0.066 | -0.475 | -1.689 | -0.699 | |
| 9 | Q0ZME3 | 340 | **SDADSP s KDTF** | 0.16 | -0.041 | -0.704 | -0.195 | |
| 9 | Q0ZME3 | 343 | **DSPSKD t FELR** | 0.112 | -0.276 | -1.359 | -0.508 | |
| 9 | Q0ZME3 | 349 | **TFELRY s GSIR** | 0.273 | 0.065 | -0.422 | -0.028 | |
| 9 | Q0ZME3 | 351 | **ELRYSG s IRFD** | 0.488 | 0.322 | 0.106 | 0.305 | |
| 9 | Q0ZME3 | 356 | **GSIRFD s TLPG** | 0.435 | 0.644 | -0.394 | 0.228 | |
| 9 | Q0ZME3 | 357 | **SIRFDS t LPGF** | 0.688 | 0.679 | 0.393 | 0.587 | very likely, matches RXXXpS/pTXP/G 1433-binding motif |
| 9 | Q0ZME3 | 363 | **TLPGFE t IMKV** | 0.091 | -0.17 | -1.504 | -0.528 | |
| 9 | Q0ZME3 | 378 | **LDAYVN s NQNT** | 0.151 | -0.097 | -1.006 | -0.317 | |
| 9 | Q0ZME3 | 382 | **VNSNQN t VSGS** | 0.061 | -0.28 | -1.312 | -0.51 | |
| 9 | Q0ZME3 | 384 | **SNQNTV s GSLS** | 0.05 | -0.223 | -1.313 | -0.495 | |
| 9 | Q0ZME3 | 386 | **QNTVSG s LSPK** | 0.131 | -0.094 | -0.683 | -0.215 | |
| 9 | Q0ZME3 | 388 | **TVSGSL s PKPQ** | 0.045 | -0.343 | -1.285 | -0.528 | |
| 9 | Q0ZME3 | 400 | **KRGVKQ s PESF** | 0.02 | -0.311 | -1.875 | -0.722 | |
| 9 | Q0ZME3 | 403 | **VKQSPE s FDSL** | 0.231 | -0.019 | -0.605 | -0.131 | |
| 9 | Q0ZME3 | 406 | **SPESFD s LNLS** | 0.163 | -0.199 | -1.126 | -0.387 | |
| 9 | Q0ZME3 | 410 | **FDSLNL s ADTQ** | 0.116 | -0.122 | -1.151 | -0.386 | |
| 9 | Q0ZME3 | 413 | **LNLSAD t QHIS** | 0.147 | -0.263 | -1.267 | -0.461 | |

| | | | | | | | |
|---|---|---|---|---|---|---|---|
| 9 | Q0ZME3 | 417 | **ADTQHI s NDFT** | 0.142 | -0.23 | -1.01 | -0.366 |
| 9 | Q0ZME3 | 421 | **HISNDF t PEDH** | 0.073 | -0.292 | -1.147 | -0.455 |
| 9 | Q0ZME3 | 426 | **FTPEDH s LLAT** | 0.213 | 0.119 | -0.692 | -0.12 |
| 9 | Q0ZME3 | 430 | **DHSLLA t LDDP** | 0.375 | 0.032 | -0.821 | -0.138 |
| 9 | Q0ZME3 | 439 | **DPYVED s VA--** | 0.072 | -0.157 | -1.67 | -0.585 |

**Supplementary Table 3.** Prediction of the 14-3-3-binding sites in nine bat coronavirus N proteins as the output of 14-3-3-Pred (Madeira et al 2015).

| Sequence | Identifier | Site | Peptide_[-6:4] | ANN | PSSM | SVM | Consensus | Manual inspection - remark |
|---|---|---|---|---|---|---|---|---|
| NCAP_BCHK5 Nucleoprotein OS=Bat coronavirus HKU5 OX=694008 GN=N PE=3 SV=1 | | | | | | | | higher consesus score indicates more probable 14-3-3 binding |
| 1 | A3EXD7 | 3 | ---MA t PAPP | 0.065 | -0.743 | -1.499 | -0.726 | |
| 1 | A3EXD7 | 18 | FANDNE t PTNS | 0.087 | -0.354 | -1.751 | -0.673 | |
| 1 | A3EXD7 | 20 | NDNETP t NSQR | 0.037 | -0.322 | -1.746 | -0.677 | |
| 1 | A3EXD7 | 22 | NETPTN s QRSG | 0.097 | -0.179 | -1.348 | -0.477 | |
| 1 | A3EXD7 | 25 | PTNSQR s GRPR | 0.073 | -0.302 | -1.526 | -0.585 | |
| 1 | A3EXD7 | 30 | RSGRPR t KPRP | 0.67 | 1.053 | 0.36 | 0.694 | matches RXXpS/pTXP/G consensus |
| 1 | A3EXD7 | 38 | PRPAPN t TVSW | 0.095 | 0.033 | -1.106 | -0.326 | |
| 1 | A3EXD7 | 39 | RPAPNT t VSWF | 0.106 | -0.146 | -0.818 | -0.286 | |
| 1 | A3EXD7 | 41 | APNTTV s WFTG | 0.181 | -0.04 | -0.986 | -0.282 | |
| 1 | A3EXD7 | 44 | TTVSWF t GLTQ | 0.084 | -0.224 | -1.299 | -0.48 | |
| 1 | A3EXD7 | 47 | SWFTGL t QHGK | 0.06 | -0.306 | -1.687 | -0.644 | |
| 1 | A3EXD7 | 68 | VPLNAN s TPAQ | 0.637 | 0.508 | 0.177 | 0.441 | |
| 1 | A3EXD7 | 69 | PLNANS t PAQN | 0.102 | -0.387 | -1.204 | -0.496 | |
| 1 | A3EXD7 | 86 | QDRKIN t GNGT | 0.402 | 0.093 | -0.109 | 0.129 | |
| 1 | A3EXD7 | 90 | INTGNG t KPLA | 0.436 | 0.28 | 0.105 | 0.274 | |
| 1 | A3EXD7 | 102 | RWYFYY t GTGP | 0.199 | -0.018 | -0.67 | -0.163 | |
| 1 | A3EXD7 | 104 | YFYYTG t GPEA | 0.364 | 0.421 | 0.014 | 0.266 | |
| 1 | A3EXD7 | 114 | ANLPFR s VKDG | 0.13 | -0.191 | -1.066 | -0.376 | |
| 1 | A3EXD7 | 128 | VHENGA t DAPS | 0.071 | -0.247 | -1.517 | -0.564 | |
| 1 | A3EXD7 | 132 | GATDAP s VFGT | 0.282 | -0.079 | -0.586 | -0.128 | |
| 1 | A3EXD7 | 136 | APSVFG t RNPA | 0.057 | -0.414 | -1.626 | -0.661 | |
| 1 | A3EXD7 | 147 | NDPAIV t QFAP | 0.12 | -0.246 | -1.006 | -0.377 | |
| 1 | A3EXD7 | 153 | TQFAPG t TLPK | 0.045 | -0.422 | -1.485 | -0.621 | |
| 1 | A3EXD7 | 154 | QFAPGT t LPKN | 0.523 | 0.439 | 0.261 | 0.408 | |
| 1 | A3EXD7 | 164 | NFHIEG t GGNS | 0.084 | -0.146 | -1.39 | -0.484 | |
| 1 | A3EXD7 | 168 | EGTGGN s QSSS | 0.035 | -0.277 | -1.652 | -0.631 | |
| 1 | A3EXD7 | 170 | TGGNSQ s SSRA | 0.109 | -0.087 | -0.838 | -0.272 | |
| 1 | A3EXD7 | 171 | GGNSQS s SRAS | 0.078 | -0.229 | -1.387 | -0.513 | |
| 1 | A3EXD7 | 172 | GNSQSS s RASS | 0.104 | -0.172 | -1.076 | -0.381 | |

| | | | | | | | | |
|---|---|---|---|---|---|---|---|---|
| 1 | A3EXD7 | 175 | QSSSRA s SRSS | 0.118 | -0.193 | -1.157 | -0.411 | |
| 1 | A3EXD7 | 176 | SSSRAS s RSSS | 0.15 | 0.471 | -0.832 | -0.07 | |
| 1 | A3EXD7 | 178 | SRASSR s SSRS | 0.101 | 0.066 | -1.281 | -0.371 | |
| 1 | A3EXD7 | 179 | RASSRS s SRSS | 0.081 | -0.225 | -1.388 | -0.511 | |
| 1 | A3EXD7 | 180 | ASSRSS s RSSS | 0.26 | 0.639 | -0.384 | 0.172 | |
| 1 | A3EXD7 | 182 | SRSSSR s SSRN | 0.122 | 0.083 | -1.316 | -0.37 | |
| 1 | A3EXD7 | 183 | RSSSRS s SRNG | 0.183 | -0.117 | -1.086 | -0.34 | |
| 1 | A3EXD7 | 184 | SSSRSS s RNGR | 0.521 | 0.75 | 0.296 | 0.522 | |
| 1 | A3EXD7 | 189 | SSRNGR s NNSS | 0.051 | -0.089 | -1.458 | -0.499 | |
| 1 | A3EXD7 | 192 | NGRSNN s SRNA | 0.439 | 0.107 | -0.445 | 0.034 | |
| 1 | A3EXD7 | 193 | GRSNNS s RNAS | 0.198 | 0.11 | -0.545 | -0.079 | |
| 1 | A3EXD7 | 197 | NSSRNA s PAPH | 0.246 | 0.263 | -0.567 | -0.019 | |
| 1 | A3EXD7 | 211 | DVVGAG t LSVL | 0.198 | 0.049 | -0.769 | -0.174 | |
| 1 | A3EXD7 | 213 | VGAGTL s VLLD | 0.078 | -0.084 | -0.936 | -0.314 | |
| 1 | A3EXD7 | 238 | KQPKVI t KKDA | 0.283 | -0.09 | -0.485 | -0.097 | |
| 1 | A3EXD7 | 256 | RHKRVA t KGYN | 0.807 | 0.968 | 0.364 | 0.713 | matches RXXpS/pTXP/G consensus |
| 1 | A3EXD7 | 275 | GPGPLQ s NFGD | 0.118 | -0.196 | -1.119 | -0.399 | |
| 1 | A3EXD7 | 287 | QYNKLG t EDPR | 0.227 | -0.026 | -0.805 | -0.201 | |
| 1 | A3EXD7 | 301 | IAELAP s ASAF | 0.209 | -0.07 | -0.79 | -0.217 | |
| 1 | A3EXD7 | 303 | ELAPSA s AFMS | 0.401 | 0.212 | -0.25 | 0.121 | |
| 1 | A3EXD7 | 307 | SASAFM s TSQF | 0.089 | -0.226 | -1.349 | -0.495 | |
| 1 | A3EXD7 | 308 | ASAFMS t SQFK | 0.086 | -0.388 | -1.277 | -0.526 | |
| 1 | A3EXD7 | 309 | SAFMST s QFKV | 0.194 | -0.019 | -1.101 | -0.309 | |
| 1 | A3EXD7 | 314 | TSQFKV t HQSN | 0.201 | -0.094 | -0.876 | -0.256 | |
| 1 | A3EXD7 | 317 | FKVTHQ s NDEN | 0.517 | 0.207 | -0.434 | 0.097 | |
| 1 | A3EXD7 | 329 | EPVYFL s YSGA | 0.042 | -0.306 | -1.412 | -0.559 | |
| 1 | A3EXD7 | 331 | VYFLSY s GAIK | 0.334 | 0.128 | -0.356 | 0.035 | |
| 1 | A3EXD7 | 359 | NIDAYK s FPKK | 0.745 | 0.558 | 0.464 | 0.589 | |
| 1 | A3EXD7 | 370 | ERKQKP s GDDA | 0.302 | 0.127 | -0.209 | 0.073 | |
| 1 | A3EXD7 | 376 | SGDDAA t APAT | 0.605 | 0.532 | -0.063 | 0.358 | |
| 1 | A3EXD7 | 380 | AATAPA t SQME | 0.135 | -0.322 | -1.152 | -0.446 | |
| 1 | A3EXD7 | 381 | ATAPAT s QMED | 0.334 | -0.077 | -0.723 | -0.155 | |
| 1 | A3EXD7 | 403 | KRVVQG s IPQR | 0.244 | 0.47 | -0.517 | 0.066 | |

| | | | | | | | |
|---|---|---|---|---|---|---|---|
| 1 | A3EXD7 | 408 | GSIPQR s AGVP | 0.102 | -0.064 | -1.407 | -0.456 |
| 1 | A3EXD7 | 413 | RSAGVP s FEDV | 0.323 | -0.001 | -0.633 | -0.104 |
| 1 | A3EXD7 | 425 | DAIFPD s EA-- | 0.145 | -0.062 | -1.241 | -0.386 |
| NCAP_BCHK9 Nucleoprotein OS=Bat coronavirus HKU9 OX=694006 GN=N PE=3 SV=1 | | | | | | | |
| 2 | A3EXH0 | 2 | ----M s GRNR | 0.307 | -0.162 | -0.623 | -0.159 |
| 2 | A3EXH0 | 7 | MSGRNR s RSGT | 0.308 | 0.582 | -0.29 | 0.2 |
| 2 | A3EXH0 | 9 | GRNRSR s GTPS | 0.727 | 1.148 | 0.374 | 0.75 |
| 2 | A3EXH0 | 11 | NRSRSG t PSPK | 0.376 | 0.718 | -0.052 | 0.347 |
| 2 | A3EXH0 | 13 | SRSGTP s PKVT | 0.032 | -0.276 | -1.563 | -0.602 |
| 2 | A3EXH0 | 17 | TPSPKV t FKQE | 0.168 | -0.247 | -0.826 | -0.302 |
| 2 | A3EXH0 | 22 | VTFKQE s DGSD | 0.275 | -0.019 | -0.96 | -0.235 |
| 2 | A3EXH0 | 25 | KQESDG s DSES | 0.059 | -0.319 | -1.556 | -0.605 |
| 2 | A3EXH0 | 27 | ESDGSD s ESER | 0.198 | 0.032 | -0.707 | -0.159 |
| 2 | A3EXH0 | 29 | DGSDSE s ERRN | 0.421 | 0.045 | -0.706 | -0.08 |
| 2 | A3EXH0 | 46 | RPKNNN s RGSA | 0.247 | -0.063 | -0.623 | -0.146 |
| 2 | A3EXH0 | 49 | NNNSRG s APKP | 0.307 | 0.323 | -0.66 | -0.01 |
| 2 | A3EXH0 | 65 | APPQNV s WFAP | 0.232 | -0.058 | -0.591 | -0.139 |
| 2 | A3EXH0 | 73 | FAPLVQ t GKAE | 0.126 | -0.1 | -1.001 | -0.325 |
| 2 | A3EXH0 | 89 | GEGVPV s QGVD | 0.232 | -0.094 | -0.969 | -0.277 |
| 2 | A3EXH0 | 94 | VSQGVD s TYEH | 0.108 | -0.281 | -1.446 | -0.54 |
| 2 | A3EXH0 | 95 | SQGVDS t YEHG | 0.021 | -0.39 | -1.819 | -0.729 |
| 2 | A3EXH0 | 104 | HGYWLR t QRSF | 0.043 | -0.273 | -1.489 | -0.573 |
| 2 | A3EXH0 | 107 | WLRTQR s FQKG | 0.434 | 0.296 | -0.357 | 0.124 |
| 2 | A3EXH0 | 126 | RWYFYY t GTGR | 0.291 | 0.057 | -0.421 | -0.024 |
| 2 | A3EXH0 | 128 | YFYYTG t GRFG | 0.065 | -0.204 | -1.031 | -0.39 |
| 2 | A3EXH0 | 138 | GDLRFG t KNPD | 0.51 | 0.68 | 0.147 | 0.446 |
| 2 | A3EXH0 | 161 | RLGDMG t RNPS | 0.137 | -0.19 | -1.164 | -0.406 |
| 2 | A3EXH0 | 165 | MGTRNP s NDGA | 0.72 | 0.786 | 0.652 | 0.719 |
| 2 | A3EXH0 | 189 | YAEGRG s RGNS | 0.174 | -0.047 | -0.997 | -0.29 |
| 2 | A3EXH0 | 193 | RGSRGN s RSSS | 0.242 | 0.593 | -0.537 | 0.099 |
| 2 | A3EXH0 | 195 | SRGNSR s SSRN | 0.088 | 0.142 | -1.245 | -0.338 |
| 2 | A3EXH0 | 196 | RGNSRS s SRNS | 0.145 | -0.092 | -1.226 | -0.391 |

| | | | | | | | | |
|---|---|---|---|---|---|---|---|---|
| 2 | A3EXH0 | 197 | GNSRSS s RNSS | 0.463 | 0.789 | 0.144 | 0.465 | |
| 2 | A3EXH0 | 200 | RSSSRN s SRAS | 0.331 | 0.003 | -0.662 | -0.109 | |
| 2 | A3EXH0 | 201 | SSSRNS s RASS | 0.129 | 0.401 | -0.836 | -0.102 | |
| 2 | A3EXH0 | 204 | RNSSRA s SRGN | 0.286 | -0.106 | -0.732 | -0.184 | |
| 2 | A3EXH0 | 205 | NSSRAS s RGNS | 0.528 | 0.738 | -0.05 | 0.405 | |
| 2 | A3EXH0 | 209 | ASSRGN s RASS | 0.211 | 0.478 | -0.715 | -0.009 | |
| 2 | A3EXH0 | 212 | RGNSRA s SRGA | 0.321 | -0.068 | -0.625 | -0.124 | |
| 2 | A3EXH0 | 213 | GNSRAS s RGAS | 0.618 | 0.784 | 0.298 | 0.567 | matches RXXpS/pTXP/G consensus |
| 2 | A3EXH0 | 217 | ASSRGA s PGRP | 0.278 | 0.298 | -0.615 | -0.013 | |
| 2 | A3EXH0 | 226 | RPAANP s TEPW | 0.118 | -0.206 | -0.999 | -0.362 | |
| 2 | A3EXH0 | 227 | PAANPS t EPWM | 0.351 | 0.356 | 0.088 | 0.265 | |
| 2 | A3EXH0 | 243 | KLERLE s QVSG | 0.401 | 0.698 | -0.426 | 0.224 | |
| 2 | A3EXH0 | 246 | RLESQV s GTKP | 0.214 | -0.136 | -0.949 | -0.29 | |
| 2 | A3EXH0 | 248 | ESQVSG t KPAT | 0.491 | 0.569 | 0.214 | 0.425 | |
| 2 | A3EXH0 | 252 | SGTKPA t KNPV | 0.254 | -0.13 | -0.68 | -0.185 | |
| 2 | A3EXH0 | 259 | KNPVQV t KNEA | 0.117 | -0.23 | -1.01 | -0.374 | |
| 2 | A3EXH0 | 275 | KLRHKR t AHKG | 0.329 | 0.224 | -0.756 | -0.068 | |
| 2 | A3EXH0 | 280 | RTAHKG s GVTV | 0.081 | -0.256 | -1.455 | -0.543 | |
| 2 | A3EXH0 | 283 | HKGSGV t VNYG | 0.265 | -0.023 | -0.526 | -0.095 | |
| 2 | A3EXH0 | 308 | EMIKLG t DDPR | 0.146 | 0.035 | -1.029 | -0.283 | |
| 2 | A3EXH0 | 324 | QMAPNV s SFLF | 0.15 | -0.034 | -0.643 | -0.176 | |
| 2 | A3EXH0 | 325 | MAPNVS s FLFM | 0.071 | -0.187 | -1.228 | -0.448 | |
| 2 | A3EXH0 | 330 | SSFLFM s HLST | 0.087 | -0.147 | -1.465 | -0.508 | |
| 2 | A3EXH0 | 333 | LFMSHL s TRDE | 0.186 | -0.084 | -0.717 | -0.205 | |
| 2 | A3EXH0 | 334 | FMSHLS t RDED | 0.129 | -0.021 | -1.041 | -0.311 | |
| 2 | A3EXH0 | 362 | PNYEQW t KILA | 0.063 | -0.195 | -1.085 | -0.406 | |
| 2 | A3EXH0 | 378 | YKDFPP t EPKK | 0.588 | 0.398 | 0.109 | 0.365 | |
| 2 | A3EXH0 | 390 | KKKKEE t AQDT | 0.581 | 0.079 | -0.385 | 0.092 | |
| 2 | A3EXH0 | 394 | EETAQD t VIFE | 0.058 | -0.296 | -1.513 | -0.584 | |
| 2 | A3EXH0 | 401 | VIFEDA s TGTD | 0.267 | 0.025 | -0.702 | -0.137 | |
| 2 | A3EXH0 | 402 | IFEDAS t GTDQ | 0.34 | -0.07 | -0.54 | -0.09 | |
| 2 | A3EXH0 | 404 | EDASTG t DQTV | 0.052 | -0.233 | -1.536 | -0.572 | |
| 2 | A3EXH0 | 407 | STGTDQ t VVKV | 0.078 | -0.202 | -1.681 | -0.602 | |

| | | | | | | | |
|---|---|---|---|---|---|---|---|
| 2 | A3EXH0 | 420 | KDQDAQ t DDEW | 0.302 | 0.103 | -0.719 | -0.105 |
| 2 | A3EXH0 | 430 | WLGGDE t VYED | 0.207 | -0.014 | -0.674 | -0.16 |
| 2 | A3EXH0 | 441 | EDDRPK t QRRH | 0.275 | 0.373 | -0.55 | 0.033 |
| 2 | A3EXH0 | 450 | RHKKRG s TASR | 0.271 | 0.047 | -0.897 | -0.193 |
| 2 | A3EXH0 | 451 | HKKRGS t ASRV | 0.511 | 0.785 | -0.149 | 0.382 |
| 2 | A3EXH0 | 453 | KRGSTA s RVTI | 0.137 | 0.082 | -0.971 | -0.251 |
| 2 | A3EXH0 | 456 | STASRV t IADP | 0.238 | -0.12 | -0.796 | -0.226 |
| 2 | A3EXH0 | 461 | VTIADP t NAGA | 0.099 | -0.375 | -1.361 | -0.546 |
| 2 | A3EXH0 | 468 | NAGAER s ---- | 0.088 | -0.058 | -1.432 | -0.467 |
| **NCAP_BCHK4 Nucleoprotein OS=Bat coronavirus HKU4 OX=694007 GN=N PE=3 SV=1** | | | | | | | |
| 3 | A3EXA1 | 3 | ---MA t PAAP | 0.096 | -0.613 | -1.209 | -0.575 |
| 3 | A3EXA1 | 9 | TPAAPR t ISFA | 0.034 | -0.45 | -1.578 | -0.665 |
| 3 | A3EXA1 | 11 | AAPRTI s FADN | 0.6 | 0.826 | 0.177 | 0.534 |
| 3 | A3EXA1 | 40 | RPAPNN t VSWY | 0.243 | -0.034 | -0.707 | -0.166 |
| 3 | A3EXA1 | 42 | APNNTV s WYTG | 0.133 | -0.029 | -0.962 | -0.286 |
| 3 | A3EXA1 | 45 | NTVSWY t GLTQ | 0.098 | -0.152 | -1.132 | -0.395 |
| 3 | A3EXA1 | 48 | SWYTGL t QHGK | 0.048 | -0.328 | -1.723 | -0.668 |
| 3 | A3EXA1 | 69 | VPLNAN s TTAQ | 0.295 | -0.007 | -0.605 | -0.106 |
| 3 | A3EXA1 | 70 | PLNANS t TAQN | 0.197 | -0.114 | -0.853 | -0.257 |
| 3 | A3EXA1 | 71 | LNANST t AQNA | 0.29 | 0.154 | -0.22 | 0.075 |
| 3 | A3EXA1 | 87 | QDRKIN t GNGV | 0.474 | 0.145 | -0.235 | 0.128 |
| 3 | A3EXA1 | 103 | RWFFYY t GTGP | 0.248 | 0.004 | -0.634 | -0.127 |
| 3 | A3EXA1 | 105 | FFYYTG t GPEA | 0.435 | 0.495 | 0.066 | 0.332 |
| 3 | A3EXA1 | 115 | ANLPFR s VKDG | 0.13 | -0.191 | -1.066 | -0.376 |
| 3 | A3EXA1 | 129 | VYEEGA t DAPS | 0.098 | -0.204 | -1.371 | -0.492 |
| 3 | A3EXA1 | 133 | GATDAP s VFGT | 0.282 | -0.079 | -0.586 | -0.128 |
| 3 | A3EXA1 | 137 | APSVFG t RNPA | 0.057 | -0.414 | -1.626 | -0.661 |
| 3 | A3EXA1 | 154 | CQFAPG t LIPK | 0.142 | -0.208 | -0.548 | -0.205 |
| 3 | A3EXA1 | 165 | NFHIEG t GGNS | 0.084 | -0.146 | -1.39 | -0.484 |
| 3 | A3EXA1 | 169 | EGTGGN s QSSS | 0.035 | -0.277 | -1.652 | -0.631 |
| 3 | A3EXA1 | 171 | TGGNSQ s SSRA | 0.109 | -0.087 | -0.838 | -0.272 |
| 3 | A3EXA1 | 172 | GGNSQS s SRAS | 0.078 | -0.229 | -1.387 | -0.513 |

| | | | | | | | | |
|---|---|---|---|---|---|---|---|---|
| 3 | A3EXA1 | 173 | GNSQSS s RASS | 0.104 | -0.172 | -1.076 | -0.381 | |
| 3 | A3EXA1 | 176 | QSSSRA s SNSR | 0.184 | -0.068 | -0.789 | -0.224 | |
| 3 | A3EXA1 | 177 | SSSRAS s NSRN | 0.292 | 0.534 | -0.695 | 0.044 | |
| 3 | A3EXA1 | 179 | SRASSN s RNSS | 0.206 | 0.209 | -0.703 | -0.096 | |
| 3 | A3EXA1 | 182 | SSNSRN s SRSS | 0.096 | -0.204 | -1.494 | -0.534 | |
| 3 | A3EXA1 | 183 | SNSRNS s RSSS | 0.093 | 0.369 | -0.933 | -0.157 | |
| 3 | A3EXA1 | 185 | SRNSSR s SSRG | 0.118 | 0.083 | -1.36 | -0.386 | |
| 3 | A3EXA1 | 186 | RNSSRS s SRGG | 0.154 | -0.227 | -0.956 | -0.343 | |
| 3 | A3EXA1 | 187 | NSSRSS s RGGR | 0.666 | 0.834 | 0.583 | 0.694 | matches RXXpS/pTXP/G consensus |
| 3 | A3EXA1 | 192 | SSRGGR s TSNS | 0.054 | -0.066 | -1.642 | -0.551 | |
| 3 | A3EXA1 | 193 | SRGGRS t SNSR | 0.086 | 0.135 | -1.015 | -0.265 | |
| 3 | A3EXA1 | 194 | RGGRST s NSRG | 0.626 | 0.889 | 0.223 | 0.579 | |
| 3 | A3EXA1 | 196 | GRSTSN s RGTS | 0.564 | 0.361 | -0.266 | 0.22 | |
| 3 | A3EXA1 | 199 | TSNSRG t SPVS | 0.321 | 0.327 | -0.642 | 0.002 | |
| 3 | A3EXA1 | 200 | SNSRGT s PVSH | 0.049 | 0.138 | -1.258 | -0.357 | |
| 3 | A3EXA1 | 203 | RGTSPV s HGVG | 0.414 | -0.044 | -0.317 | 0.018 | |
| 3 | A3EXA1 | 208 | VSHGVG s AESL | 0.051 | -0.245 | -1.677 | -0.624 | |
| 3 | A3EXA1 | 211 | GVGSAE s LAAL | 0.303 | 0.11 | -0.477 | -0.021 | |
| 3 | A3EXA1 | 230 | RLADLE s GKSK | 0.103 | -0.158 | -1.207 | -0.421 | |
| 3 | A3EXA1 | 233 | DLESGK s KQPK | 0.283 | -0.128 | -0.516 | -0.12 | |
| 3 | A3EXA1 | 240 | KQPKVV t KKDA | 0.379 | -0.077 | -0.361 | -0.02 | |
| 3 | A3EXA1 | 258 | RHKRVA t KGFN | 0.756 | 0.929 | 0.291 | 0.659 | |
| 3 | A3EXA1 | 264 | TKGFNV t QAFG | 0.239 | -0.157 | -0.908 | -0.275 | |
| 3 | A3EXA1 | 289 | NYNKFG t EDPR | 0.331 | 0.024 | -0.662 | -0.102 | |
| 3 | A3EXA1 | 303 | MAELAP s ASAF | 0.228 | 0.021 | -0.76 | -0.17 | |
| 3 | A3EXA1 | 305 | ELAPSA s AFMS | 0.401 | 0.212 | -0.25 | 0.121 | |
| 3 | A3EXA1 | 309 | SASAFM s MSQF | 0.095 | -0.101 | -1.323 | -0.443 | |
| 3 | A3EXA1 | 311 | SAFMSM s QFKL | 0.277 | 0.156 | -0.664 | -0.077 | |
| 3 | A3EXA1 | 316 | MSQFKL t HQSN | 0.131 | -0.032 | -1.014 | -0.305 | |
| 3 | A3EXA1 | 319 | FKLTHQ s NDDK | 0.597 | 0.18 | -0.169 | 0.203 | |
| 3 | A3EXA1 | 331 | DPIYFL s YSGA | 0.071 | -0.278 | -1.411 | -0.539 | |
| 3 | A3EXA1 | 333 | IYFLSY s GAIK | 0.422 | 0.16 | -0.172 | 0.137 | |
| 3 | A3EXA1 | 354 | WLELLE s NIDA | 0.211 | -0.089 | -0.843 | -0.24 | |

| | | | | | | | | |
|---|---|---|---|---|---|---|---|---|
| 3 | A3EXA1 | 361 | NIDAYK t FPKK | 0.745 | 0.558 | 0.464 | 0.589 | |
| 3 | A3EXA1 | 371 | KERKPK t TEDG | 0.61 | 0.233 | -0.023 | 0.273 | |
| 3 | A3EXA1 | 372 | ERKPKT t EDGA | 0.249 | 0.155 | -0.264 | 0.047 | |
| 3 | A3EXA1 | 379 | EDGAVA s SSAS | 0.054 | -0.232 | -1.421 | -0.533 | |
| 3 | A3EXA1 | 380 | DGAVAS s SASQ | 0.057 | -0.285 | -1.511 | -0.58 | |
| 3 | A3EXA1 | 381 | GAVASS s ASQM | 0.214 | 0.01 | -0.796 | -0.191 | |
| 3 | A3EXA1 | 383 | VASSSA s QMED | 0.369 | 0.041 | -0.623 | -0.071 | |
| 3 | A3EXA1 | 398 | PQRKPK s RVAG | 0.583 | 0.249 | 0.232 | 0.355 | |
| 3 | A3EXA1 | 403 | KSRVAG s ITMR | 0.21 | -0.027 | -0.876 | -0.231 | |
| 3 | A3EXA1 | 405 | RVAGSI t MRSG | 0.269 | 0.227 | -0.35 | 0.049 | |
| 3 | A3EXA1 | 408 | GSITMR s GSLP | 0.063 | -0.294 | -1.582 | -0.604 | |
| 3 | A3EXA1 | 410 | ITMRSG s LPAL | 0.962 | 1.708 | 2.063 | 1.578 | matches RXXpS/pTXP/G consensus |
| 3 | A3EXA1 | 418 | PALQDV t FDSE | 0.341 | -0.056 | -0.376 | -0.03 | |
| 3 | A3EXA1 | 421 | QDVTFD s EA-- | 0.077 | -0.127 | -1.525 | -0.525 | |
| **NCAP_BCHK3 Nucleoprotein OS=Bat coronavirus HKU3 OX=442736 GN=N PE=3 SV=1** | | | | | | | | |
| 4 | Q3LZX4 | 2 | ----M s DNGP | 0.33 | -0.143 | -0.5 | -0.104 |  |
| 4 | Q3LZX4 | 8 | SDNGPQ s QRSA | 0.026 | -0.467 | -1.803 | -0.748 | |
| 4 | Q3LZX4 | 11 | GPQSQR s APRI | 0.511 | 0.455 | -0.528 | 0.146 | |
| 4 | Q3LZX4 | 16 | RSAPRI t FGGP | 0.451 | 0.056 | -0.106 | 0.134 | |
| 4 | Q3LZX4 | 23 | FGGPAD s NDNN | 0.203 | 0.001 | -1.046 | -0.281 | |
| 4 | Q3LZX4 | 33 | NQDGGR s GARP | 0.026 | -0.423 | -1.823 | -0.74 | |
| 4 | Q3LZX4 | 49 | QGLPNN t ASWF | 0.131 | -0.112 | -0.879 | -0.287 | |
| 4 | Q3LZX4 | 51 | LPNNTA s WFTA | 0.106 | 0 | -0.849 | -0.248 | |
| 4 | Q3LZX4 | 54 | NTASWF t ALTQ | 0.154 | -0.094 | -1.04 | -0.327 | |
| 4 | Q3LZX4 | 57 | SWFTAL t QHGK | 0.139 | -0.161 | -1.293 | -0.438 | |
| 4 | Q3LZX4 | 76 | QGVPIN t NSGK | 0.093 | -0.279 | -1.118 | -0.435 | |
| 4 | Q3LZX4 | 78 | VPINTN s GKDD | 0.17 | -0.004 | -0.718 | -0.184 | |
| 4 | Q3LZX4 | 91 | GYYRRA t RRVR | 0.853 | 1.012 | 0.666 | 0.844 | unlikely, by analogy with SARS-CoV-2 |
| 4 | Q3LZX4 | 105 | GKMKEL s PRWY | 0.28 | -0.199 | -0.814 | -0.244 | |
| 4 | Q3LZX4 | 115 | YFYYLG t GPEA | 0.36 | 0.343 | -0.117 | 0.195 | |
| 4 | Q3LZX4 | 120 | GTGPEA s LPYG | 0.718 | 0.467 | 0 | 0.395 | |

| | | | | | | | | |
|---|---|---|---|---|---|---|---|---|
| 4 | Q3LZX4 | 135 | GIVWVA t EGAL | 0.638 | 0.329 | 0.073 | 0.347 | |
| 4 | Q3LZX4 | 141 | TEGALN t PKDH | 0.051 | -0.518 | -1.848 | -0.772 | |
| 4 | Q3LZX4 | 148 | PKDHIG t RNPN | 0.184 | -0.165 | -0.785 | -0.255 | |
| 4 | Q3LZX4 | 165 | LQLPQG t TLPK | 0.035 | -0.37 | -1.556 | -0.63 | |
| 4 | Q3LZX4 | 166 | QLPQGT t LPKG | 0.615 | 0.497 | 0.321 | 0.478 | |
| 4 | Q3LZX4 | 176 | GFYAEG s RGGS | 0.151 | -0.235 | -1.103 | -0.396 | |
| 4 | Q3LZX4 | 180 | EGSRGG s QSSS | 0.078 | 0.435 | -1.012 | -0.166 | |
| 4 | Q3LZX4 | 182 | SRGGSQ s SSRS | 0.097 | 0.129 | -1.109 | -0.294 | |
| 4 | Q3LZX4 | 183 | RGGSQS s SRSS | 0.03 | -0.356 | -1.833 | -0.72 | |
| 4 | Q3LZX4 | 184 | GGSQSS s RSSS | 0.115 | -0.112 | -1.021 | -0.339 | |
| 4 | Q3LZX4 | 186 | SQSSSR s SSRS | 0.054 | -0.207 | -1.575 | -0.576 | |
| 4 | Q3LZX4 | 187 | QSSSRS s SRSR | 0.08 | -0.264 | -1.231 | -0.472 | |
| 4 | Q3LZX4 | 188 | SSSRSS s RSRG | 0.367 | 0.621 | -0.347 | 0.214 | |
| 4 | Q3LZX4 | 190 | SRSSSR s RGNS | 0.2 | 0.22 | -1.014 | -0.198 | |
| 4 | Q3LZX4 | 194 | SRSRGN s RNST | 0.32 | 0.805 | -0.255 | 0.29 | |
| 4 | Q3LZX4 | 197 | RGNSRN s TPGS | 0.697 | 0.526 | 0.033 | 0.419 | likely, by analogy with SARS-CoV-2 |
| 4 | Q3LZX4 | 198 | GNSRNS t PGSS | 0.215 | 0.381 | -0.438 | 0.053 | |
| 4 | Q3LZX4 | 201 | RNSTPG s SRGS | 0.06 | -0.471 | -1.453 | -0.621 | |
| 4 | Q3LZX4 | 202 | NSTPGS s RGSS | 0.041 | -0.392 | -1.568 | -0.64 | |
| 4 | Q3LZX4 | 205 | PGSSRG s SPAR | 0.597 | 0.49 | 0.145 | 0.411 | |
| 4 | Q3LZX4 | 206 | GSSRGS s PARL | 0.162 | 0.272 | -0.803 | -0.123 | |
| 4 | Q3LZX4 | 212 | SPARLA s GGGE | 0.467 | 0.643 | -0.048 | 0.354 | |
| 4 | Q3LZX4 | 217 | ASGGGE t ALAL | 0.057 | -0.158 | -1.458 | -0.52 | |
| 4 | Q3LZX4 | 232 | RLNQLE s KVSG | 0.15 | -0.053 | -1.051 | -0.318 | |
| 4 | Q3LZX4 | 235 | QLESKV s GKGQ | 0.176 | -0.244 | -0.794 | -0.287 | |
| 4 | Q3LZX4 | 245 | QQQPGQ t VTKK | 0.078 | -0.201 | -1.041 | -0.388 | |
| 4 | Q3LZX4 | 247 | QPGQTV t KKSA | 0.094 | -0.175 | -0.915 | -0.332 | |
| 4 | Q3LZX4 | 250 | QTVTKK s AAEA | 0.174 | -0.112 | -0.967 | -0.302 | |
| 4 | Q3LZX4 | 255 | KSAAEA s KKPR | 0.152 | -0.249 | -0.909 | -0.335 | |
| 4 | Q3LZX4 | 263 | KPRQKR t ATKQ | 0.188 | 0.023 | -1.163 | -0.317 | |
| 4 | Q3LZX4 | 265 | RQKRTA t KQYN | 0.672 | 0.963 | 0.549 | 0.728 | unlikely, by analogy with SARS-CoV-2 |
| 4 | Q3LZX4 | 271 | TKQYNV t QAFG | 0.329 | -0.068 | -0.616 | -0.118 | |
| 4 | Q3LZX4 | 282 | RRGPEQ t QGNF | 0.166 | 0.137 | -1.106 | -0.268 | |

| | | | | | | | |
|---|---|---|---|---|---|---|---|
| 4 | Q3LZX4 | 310 | IAQFAP s ASAF | 0.266 | 0.03 | -0.65 | -0.118 |
| 4 | Q3LZX4 | 312 | QFAPSA s AFFG | 0.36 | 0.155 | -0.221 | 0.098 |
| 4 | Q3LZX4 | 318 | SAFFGM s RIGM | 0.108 | -0.181 | -1.136 | -0.403 |
| 4 | Q3LZX4 | 325 | RIGMEV t PSGT | 0.086 | -0.467 | -1.384 | -0.588 |
| 4 | Q3LZX4 | 327 | GMEVTP s GTWL | 0.102 | 0.025 | -0.991 | -0.288 |
| 4 | Q3LZX4 | 329 | EVTPSG t WLTY | 0.139 | 0.111 | -0.766 | -0.172 |
| 4 | Q3LZX4 | 332 | PSGTWL t YHGA | 0.104 | -0.218 | -1.046 | -0.387 |
| 4 | Q3LZX4 | 362 | HIDAYK t FPPT | 0.694 | 0.583 | 0.571 | 0.616 |
| 4 | Q3LZX4 | 366 | YKTFPP t EPKK | 0.643 | 0.44 | 0.307 | 0.463 |
| 4 | Q3LZX4 | 376 | KDKKKK t DEAQ | 0.295 | -0.009 | -0.522 | -0.079 |
| 4 | Q3LZX4 | 391 | RQKKQP t VTLL | 0.34 | 0.067 | -0.445 | -0.013 |
| 4 | Q3LZX4 | 393 | KKQPTV t LLPA | 0.379 | 0.038 | -0.299 | 0.039 |
| 4 | Q3LZX4 | 404 | ADMDDF s RQLQ | 0.083 | -0.22 | -1.376 | -0.504 |
| 4 | Q3LZX4 | 410 | SRQLQH s MSGA | 0.154 | 0.275 | -1.051 | -0.207 |
| 4 | Q3LZX4 | 412 | QLQHSM s GASA | 0.319 | 0.252 | -0.172 | 0.133 |
| 4 | Q3LZX4 | 415 | HSMSGA s ADST | 0.076 | -0.109 | -1.163 | -0.399 |
| 4 | Q3LZX4 | 418 | SGASAD s TQA- | 0.194 | -0.088 | -0.858 | -0.251 |
| 4 | Q3LZX4 | 419 | GASADS t QA-- | 0.086 | -0.199 | -1.563 | -0.559 |
| **NCAP_BCRP3 Nucleoprotein OS=Bat coronavirus Rp3/2004 OX=349344 GN=N PE=3 SV=1** | | | | | | | |
| 5 | Q3I5I7 | 2 | ----M s DNGP | 0.33 | -0.143 | -0.5 | -0.104 |
| 5 | Q3I5I7 | 11 | GPQNQR s APRI | 0.517 | 0.507 | -0.36 | 0.221 |
| 5 | Q3I5I7 | 16 | RSAPRI t FGGP | 0.451 | 0.056 | -0.106 | 0.134 |
| 5 | Q3I5I7 | 21 | ITFGGP t DSTD | 0.132 | -0.211 | -1.084 | -0.388 |
| 5 | Q3I5I7 | 23 | FGGPTD s TDNN | 0.111 | -0.022 | -1.181 | -0.364 |
| 5 | Q3I5I7 | 24 | GGPTDS t DNNQ | 0.135 | -0.148 | -1.322 | -0.445 |
| 5 | Q3I5I7 | 33 | NQDGGR s GARP | 0.026 | -0.423 | -1.823 | -0.74 |
| 5 | Q3I5I7 | 49 | QGLPNN t ASWF | 0.131 | -0.112 | -0.879 | -0.287 |
| 5 | Q3I5I7 | 51 | LPNNTA s WFTA | 0.106 | 0 | -0.849 | -0.248 |
| 5 | Q3I5I7 | 54 | NTASWF t ALTQ | 0.154 | -0.094 | -1.04 | -0.327 |
| 5 | Q3I5I7 | 57 | SWFTAL t QHGK | 0.139 | -0.161 | -1.293 | -0.438 |
| 5 | Q3I5I7 | 76 | QGVPIN t NSGK | 0.093 | -0.279 | -1.118 | -0.435 |
| 5 | Q3I5I7 | 78 | VPINTN s GKDD | 0.17 | -0.004 | -0.718 | -0.184 |

| | | | | | | | | |
|---|---|---|---|---|---|---|---|---|
| 5 | Q3I5I7 | 91 | GYYRRA t RRVR | 0.853 | 1.012 | 0.666 | 0.844 | unlikely, by analogy with SARS-CoV-2 |
| 5 | Q3I5I7 | 105 | GKMKEL s PRWY | 0.28 | -0.199 | -0.814 | -0.244 | |
| 5 | Q3I5I7 | 115 | YFYYLG t GPEA | 0.36 | 0.343 | -0.117 | 0.195 | |
| 5 | Q3I5I7 | 120 | GTGPEA s LPYG | 0.718 | 0.467 | 0 | 0.395 | |
| 5 | Q3I5I7 | 135 | GIVWVA t EGAL | 0.638 | 0.329 | 0.073 | 0.347 | |
| 5 | Q3I5I7 | 141 | TEGALN t PKDH | 0.051 | -0.518 | -1.848 | -0.772 | |
| 5 | Q3I5I7 | 148 | PKDHIG t RNPN | 0.184 | -0.165 | -0.785 | -0.255 | |
| 5 | Q3I5I7 | 165 | LQLPQG t TLPK | 0.035 | -0.37 | -1.556 | -0.63 | |
| 5 | Q3I5I7 | 166 | QLPQGT t LPKG | 0.615 | 0.497 | 0.321 | 0.478 | |
| 5 | Q3I5I7 | 176 | GFYAEG s RGGS | 0.151 | -0.235 | -1.103 | -0.396 | |
| 5 | Q3I5I7 | 180 | EGSRGG s QASS | 0.091 | 0.442 | -0.934 | -0.134 | |
| 5 | Q3I5I7 | 183 | RGGSQA s SRSS | 0.057 | -0.25 | -1.556 | -0.583 | |
| 5 | Q3I5I7 | 184 | GGSQAS s RSSS | 0.089 | -0.257 | -1.301 | -0.49 | |
| 5 | Q3I5I7 | 186 | SQASSR s SSRS | 0.059 | -0.177 | -1.437 | -0.518 | |
| 5 | Q3I5I7 | 187 | QASSRS s SRSR | 0.078 | -0.25 | -1.266 | -0.479 | |
| 5 | Q3I5I7 | 188 | ASSRSS s RSRG | 0.467 | 0.644 | -0.179 | 0.311 | |
| 5 | Q3I5I7 | 190 | SRSSSR s RGNS | 0.2 | 0.22 | -1.014 | -0.198 | |
| 5 | Q3I5I7 | 194 | SRSRGN s RNST | 0.32 | 0.805 | -0.255 | 0.29 | |
| 5 | Q3I5I7 | 197 | RGNSRN s TPGS | 0.697 | 0.526 | 0.033 | 0.419 | likely, by analogy with SARS-CoV-2 |
| 5 | Q3I5I7 | 198 | GNSRNS t PGSS | 0.215 | 0.381 | -0.438 | 0.053 | |
| 5 | Q3I5I7 | 201 | RNSTPG s SRGN | 0.079 | -0.424 | -1.35 | -0.565 | |
| 5 | Q3I5I7 | 202 | NSTPGS s RGNS | 0.064 | -0.303 | -1.458 | -0.566 | |
| 5 | Q3I5I7 | 206 | GSSRGN s PARM | 0.317 | 0.338 | -0.567 | 0.029 | |
| 5 | Q3I5I7 | 212 | SPARMA s GGGE | 0.557 | 0.634 | 0.176 | 0.456 | |
| 5 | Q3I5I7 | 217 | ASGGGE t ALAL | 0.057 | -0.158 | -1.458 | -0.52 | |
| 5 | Q3I5I7 | 232 | RLNQLE s KVSG | 0.15 | -0.053 | -1.051 | -0.318 | |
| 5 | Q3I5I7 | 235 | QLESKV s GRSQ | 0.129 | -0.285 | -1.029 | -0.395 | |
| 5 | Q3I5I7 | 238 | SKVSGR s QQQQ | 0.072 | -0.278 | -1.631 | -0.612 | |
| 5 | Q3I5I7 | 245 | QQQQGQ t VTKK | 0.118 | -0.14 | -0.816 | -0.279 | |
| 5 | Q3I5I7 | 247 | QQGQTV t KKSA | 0.092 | -0.151 | -0.76 | -0.273 | |
| 5 | Q3I5I7 | 250 | QTVTKK s AAEA | 0.174 | -0.112 | -0.967 | -0.302 | |
| 5 | Q3I5I7 | 255 | KSAAEA s KKPR | 0.152 | -0.249 | -0.909 | -0.335 | |
| 5 | Q3I5I7 | 263 | KPRQKR t ATKQ | 0.188 | 0.023 | -1.163 | -0.317 | |

| | | | | | | | | |
|---|---|---|---|---|---|---|---|---|
| 5 | Q3I5I7 | 265 | RQKRTA t KQYN | 0.672 | 0.963 | 0.549 | 0.728 | unlikely, by analogy with SARS-CoV-2 |
| 5 | Q3I5I7 | 271 | TKQYNV t QAFG | 0.329 | -0.068 | -0.616 | -0.118 | |
| 5 | Q3I5I7 | 282 | RRGPEQ t QGNF | 0.166 | 0.137 | -1.106 | -0.268 | |
| 5 | Q3I5I7 | 296 | ELIRQG t DYKH | 0.325 | 0.753 | -0.495 | 0.194 | |
| 5 | Q3I5I7 | 310 | IAQFAP s ASAF | 0.266 | 0.03 | -0.65 | -0.118 | |
| 5 | Q3I5I7 | 312 | QFAPSA s AFFG | 0.36 | 0.155 | -0.221 | 0.098 | |
| 5 | Q3I5I7 | 318 | SAFFGM s RIGM | 0.108 | -0.181 | -1.136 | -0.403 | |
| 5 | Q3I5I7 | 325 | RIGMEV t PSGT | 0.086 | -0.467 | -1.384 | -0.588 | |
| 5 | Q3I5I7 | 327 | GMEVTP s GTWL | 0.102 | 0.025 | -0.991 | -0.288 | |
| 5 | Q3I5I7 | 329 | EVTPSG t WLTY | 0.139 | 0.111 | -0.766 | -0.172 | |
| 5 | Q3I5I7 | 332 | PSGTWL t YHGA | 0.104 | -0.218 | -1.046 | -0.387 | |
| 5 | Q3I5I7 | 366 | YKIFPP t EPKK | 0.613 | 0.456 | 0.171 | 0.413 | |
| 5 | Q3I5I7 | 376 | KDKKKK t DEAQ | 0.295 | -0.009 | -0.522 | -0.079 | |
| 5 | Q3I5I7 | 391 | RQKKQP t VTLL | 0.34 | 0.067 | -0.445 | -0.013 | |
| 5 | Q3I5I7 | 393 | KKQPTV t LLPA | 0.379 | 0.038 | -0.299 | 0.039 | |
| 5 | Q3I5I7 | 404 | ADMDDF s RQLQ | 0.083 | -0.22 | -1.376 | -0.504 | |
| 5 | Q3I5I7 | 410 | SRQLQN s MSGA | 0.138 | 0.195 | -1.176 | -0.281 | |
| 5 | Q3I5I7 | 412 | QLQNSM s GASA | 0.283 | 0.267 | -0.12 | 0.143 | |
| 5 | Q3I5I7 | 415 | NSMSGA s ADST | 0.08 | -0.166 | -1.258 | -0.448 | |
| 5 | Q3I5I7 | 418 | SGASAD s TQA- | 0.194 | -0.088 | -0.858 | -0.251 | |
| 5 | Q3I5I7 | 419 | GASADS t QA-- | 0.086 | -0.199 | -1.563 | -0.559 | |
| **NCAP_BC133 Nucleoprotein OS=Bat coronavirus 133/2005 OX=389230 GN=N PE=3 SV=1** | | | | | | | | |
| 6 | Q0Q4E6 | 3 | ---MA t PAAP | 0.096 | -0.613 | -1.209 | -0.575 | |
| 6 | Q0Q4E6 | 9 | TPAAPR t ISFA | 0.034 | -0.45 | -1.578 | -0.665 | |
| 6 | Q0Q4E6 | 11 | AAPRTI s FADN | 0.6 | 0.826 | 0.177 | 0.534 | |
| 6 | Q0Q4E6 | 40 | RPAPNN t VSWY | 0.243 | -0.034 | -0.707 | -0.166 | |
| 6 | Q0Q4E6 | 42 | APNNTV s WYTG | 0.133 | -0.029 | -0.962 | -0.286 | |
| 6 | Q0Q4E6 | 45 | NTVSWY t GLTQ | 0.098 | -0.152 | -1.132 | -0.395 | |
| 6 | Q0Q4E6 | 48 | SWYTGL t QHGK | 0.048 | -0.328 | -1.723 | -0.668 | |
| 6 | Q0Q4E6 | 69 | VPLNAN s TTAQ | 0.295 | -0.007 | -0.605 | -0.106 | |
| 6 | Q0Q4E6 | 70 | PLNANS t TAQN | 0.197 | -0.114 | -0.853 | -0.257 | |
| 6 | Q0Q4E6 | 71 | LNANST t AQNA | 0.29 | 0.154 | -0.22 | 0.075 | |

| | | | | | | | | |
|---|---|---|---|---|---|---|---|---|
| 6 | Q0Q4E6 | 87 | **QDRKIN t GNGV** | 0.474 | 0.145 | -0.235 | 0.128 | |
| 6 | Q0Q4E6 | 103 | **RWFFYY t GTGP** | 0.248 | 0.004 | -0.634 | -0.127 | |
| 6 | Q0Q4E6 | 105 | **FFYYTG t GPEA** | 0.435 | 0.495 | 0.066 | 0.332 | |
| 6 | Q0Q4E6 | 115 | **ANLPFR s VKDG** | 0.13 | -0.191 | -1.066 | -0.376 | |
| 6 | Q0Q4E6 | 129 | **VYEEGA t DAPS** | 0.098 | -0.204 | -1.371 | -0.492 | |
| 6 | Q0Q4E6 | 133 | **GATDAP s VFGT** | 0.282 | -0.079 | -0.586 | -0.128 | |
| 6 | Q0Q4E6 | 137 | **APSVFG t RNPA** | 0.057 | -0.414 | -1.626 | -0.661 | |
| 6 | Q0Q4E6 | 154 | **CQFAPG t LIPK** | 0.142 | -0.208 | -0.548 | -0.205 | |
| 6 | Q0Q4E6 | 165 | **NFHIEG t GGNS** | 0.084 | -0.146 | -1.39 | -0.484 | |
| 6 | Q0Q4E6 | 169 | **EGTGGN s QSSS** | 0.035 | -0.277 | -1.652 | -0.631 | |
| 6 | Q0Q4E6 | 171 | **TGGNSQ s SSRA** | 0.109 | -0.087 | -0.838 | -0.272 | |
| 6 | Q0Q4E6 | 172 | **GGNSQS s SRAS** | 0.078 | -0.229 | -1.387 | -0.513 | |
| 6 | Q0Q4E6 | 173 | **GNSQSS s RASS** | 0.104 | -0.172 | -1.076 | -0.381 | |
| 6 | Q0Q4E6 | 176 | **QSSSRA s SNSR** | 0.184 | -0.068 | -0.789 | -0.224 | |
| 6 | Q0Q4E6 | 177 | **SSSRAS s NSRN** | 0.292 | 0.534 | -0.695 | 0.044 | |
| 6 | Q0Q4E6 | 179 | **SRASSN s RNSS** | 0.206 | 0.209 | -0.703 | -0.096 | |
| 6 | Q0Q4E6 | 182 | **SSNSRN s SRSN** | 0.123 | -0.158 | -1.391 | -0.475 | |
| 6 | Q0Q4E6 | 183 | **SNSRNS s RSNS** | 0.158 | 0.458 | -0.823 | -0.069 | |
| 6 | Q0Q4E6 | 185 | **SRNSSR s NSRG** | 0.122 | 0.081 | -1.424 | -0.407 | |
| 6 | Q0Q4E6 | 187 | **NSSRSN s RGGR** | 0.83 | 0.956 | 0.848 | 0.878 | matches RXXpS/pTXP/G consensus |
| 6 | Q0Q4E6 | 192 | **NSRGGR s TSNS** | 0.082 | -0.002 | -1.363 | -0.428 | |
| 6 | Q0Q4E6 | 193 | **SRGGRS t SNSR** | 0.086 | 0.135 | -1.015 | -0.265 | |
| 6 | Q0Q4E6 | 194 | **RGGRST s NSRG** | 0.626 | 0.889 | 0.223 | 0.579 | |
| 6 | Q0Q4E6 | 196 | **GRSTSN s RGTS** | 0.564 | 0.361 | -0.266 | 0.22 | |
| 6 | Q0Q4E6 | 199 | **TSNSRG t SPVS** | 0.321 | 0.327 | -0.642 | 0.002 | |
| 6 | Q0Q4E6 | 200 | **SNSRGT s PVSH** | 0.049 | 0.138 | -1.258 | -0.357 | |
| 6 | Q0Q4E6 | 203 | **RGTSPV s HGVG** | 0.414 | -0.044 | -0.317 | 0.018 | |
| 6 | Q0Q4E6 | 208 | **VSHGVG s AESL** | 0.051 | -0.245 | -1.677 | -0.624 | |
| 6 | Q0Q4E6 | 211 | **GVGSAE s LAAL** | 0.303 | 0.11 | -0.477 | -0.021 | |
| 6 | Q0Q4E6 | 230 | **RLADLE s GKSK** | 0.103 | -0.158 | -1.207 | -0.421 | |
| 6 | Q0Q4E6 | 233 | **DLESGK s KQPK** | 0.283 | -0.128 | -0.516 | -0.12 | |
| 6 | Q0Q4E6 | 240 | **KQPKVV t KKDA** | 0.379 | -0.077 | -0.361 | -0.02 | |
| 6 | Q0Q4E6 | 258 | **RHKRVA t KGFN** | 0.756 | 0.929 | 0.291 | 0.659 | matches RXXpS/pTXP/G consensus |

| | | | | | | | | |
|---|---|---|---|---|---|---|---|---|
| 6 | Q0Q4E6 | 264 | TKGFNV t QAFG | 0.239 | -0.157 | -0.908 | -0.275 | |
| 6 | Q0Q4E6 | 289 | NYNKFG t EDPR | 0.331 | 0.024 | -0.662 | -0.102 | |
| 6 | Q0Q4E6 | 303 | MAELAP s ASAF | 0.228 | 0.021 | -0.76 | -0.17 | |
| 6 | Q0Q4E6 | 305 | ELAPSA s AFMS | 0.401 | 0.212 | -0.25 | 0.121 | |
| 6 | Q0Q4E6 | 309 | SASAFM s MSQF | 0.095 | -0.101 | -1.323 | -0.443 | |
| 6 | Q0Q4E6 | 311 | SAFMSM s QFKL | 0.277 | 0.156 | -0.664 | -0.077 | |
| 6 | Q0Q4E6 | 316 | MSQFKL t HQSN | 0.131 | -0.032 | -1.014 | -0.305 | |
| 6 | Q0Q4E6 | 319 | FKLTHQ s NDDK | 0.597 | 0.18 | -0.169 | 0.203 | |
| 6 | Q0Q4E6 | 331 | DPIYFL s YSGA | 0.071 | -0.278 | -1.411 | -0.539 | |
| 6 | Q0Q4E6 | 333 | IYFLSY s GAIK | 0.422 | 0.16 | -0.172 | 0.137 | |
| 6 | Q0Q4E6 | 361 | NIDAYK t FPKK | 0.745 | 0.558 | 0.464 | 0.589 | |
| 6 | Q0Q4E6 | 371 | KERKPK t TEDG | 0.61 | 0.233 | -0.023 | 0.273 | |
| 6 | Q0Q4E6 | 372 | ERKPKT t EDGA | 0.249 | 0.155 | -0.264 | 0.047 | |
| 6 | Q0Q4E6 | 380 | DGAVVA s SSAS | 0.126 | -0.126 | -1.112 | -0.371 | |
| 6 | Q0Q4E6 | 381 | GAVVAS s SASQ | 0.051 | -0.261 | -1.694 | -0.635 | |
| 6 | Q0Q4E6 | 382 | AVVASS s ASQM | 0.148 | -0.016 | -0.894 | -0.254 | |
| 6 | Q0Q4E6 | 384 | VASSSA s QMED | 0.369 | 0.041 | -0.623 | -0.071 | |
| 6 | Q0Q4E6 | 399 | PQRKPK s RVAG | 0.583 | 0.249 | 0.232 | 0.355 | |
| 6 | Q0Q4E6 | 404 | KSRVAG s ITMR | 0.21 | -0.027 | -0.876 | -0.231 | |
| 6 | Q0Q4E6 | 406 | RVAGSI t MRSG | 0.269 | 0.227 | -0.35 | 0.049 | |
| 6 | Q0Q4E6 | 409 | GSITMR s GSSP | 0.04 | -0.358 | -1.799 | -0.706 | |
| 6 | Q0Q4E6 | 411 | ITMRSG s SPAL | 0.928 | 1.569 | 1.527 | 1.341 | matches RXXpS/pTXP/G consensus |
| 6 | Q0Q4E6 | 412 | TMRSGS s PALQ | 0.034 | -0.293 | -1.712 | -0.657 | |
| 6 | Q0Q4E6 | 419 | PALQDV t FDSE | 0.341 | -0.056 | -0.376 | -0.03 | |
| 6 | Q0Q4E6 | 422 | QDVTFD s EA-- | 0.077 | -0.127 | -1.525 | -0.525 | |
| **NCAP_BC279 Nucleoprotein OS=Bat coronavirus 279/2005 OX=389167 GN=N PE=3 SV=1** | | | | | | | | |
| 7 | Q0Q468 | 2 | ----M s DNGP | 0.33 | -0.143 | -0.5 | -0.104 |  |
| 7 | Q0Q468 | 11 | GPQNQR s APRI | 0.517 | 0.507 | -0.36 | 0.221 | |
| 7 | Q0Q468 | 16 | RSAPRI t FGGP | 0.451 | 0.056 | -0.106 | 0.134 | |
| 7 | Q0Q468 | 21 | ITFGGP s DSTD | 0.132 | -0.211 | -1.084 | -0.388 | |
| 7 | Q0Q468 | 23 | FGGPSD s TDNN | 0.282 | 0.169 | -0.628 | -0.059 | |

| | | | | | | | | |
|---|---|---|---|---|---|---|---|---|
| 7 | Q0Q468 | 24 | GGPSDS t DNNQ | 0.113 | -0.185 | -1.366 | -0.479 | |
| 7 | Q0Q468 | 33 | NQDGGR s GARP | 0.026 | -0.423 | -1.823 | -0.74 | |
| 7 | Q0Q468 | 49 | QGLPNN t ASWF | 0.131 | -0.112 | -0.879 | -0.287 | |
| 7 | Q0Q468 | 51 | LPNNTA s WFTA | 0.106 | 0 | -0.849 | -0.248 | |
| 7 | Q0Q468 | 54 | NTASWF t ALTQ | 0.154 | -0.094 | -1.04 | -0.327 | |
| 7 | Q0Q468 | 57 | SWFTAL t QHGK | 0.139 | -0.161 | -1.293 | -0.438 | |
| 7 | Q0Q468 | 76 | QGVPIN t NSGK | 0.093 | -0.279 | -1.118 | -0.435 | |
| 7 | Q0Q468 | 78 | VPINTN s GKDD | 0.17 | -0.004 | -0.718 | -0.184 | |
| 7 | Q0Q468 | 91 | GYYRRA t RRVR | 0.853 | 1.012 | 0.666 | 0.844 | unlikely, by analogy with SARS-CoV-2 |
| 7 | Q0Q468 | 105 | GKMKKL s PRWY | 0.214 | -0.211 | -0.807 | -0.268 | |
| 7 | Q0Q468 | 115 | YFYYLG t GPEA | 0.36 | 0.343 | -0.117 | 0.195 | |
| 7 | Q0Q468 | 120 | GTGPEA s LPYG | 0.718 | 0.467 | 0 | 0.395 | |
| 7 | Q0Q468 | 135 | GIVWVA t EGAL | 0.638 | 0.329 | 0.073 | 0.347 | |
| 7 | Q0Q468 | 141 | TEGALN t PKDH | 0.051 | -0.518 | -1.848 | -0.772 | |
| 7 | Q0Q468 | 148 | PKDHIG t RNPN | 0.184 | -0.165 | -0.785 | -0.255 | |
| 7 | Q0Q468 | 165 | LQLPQG t TLPK | 0.035 | -0.37 | -1.556 | -0.63 | |
| 7 | Q0Q468 | 166 | QLPQGT t LPKG | 0.615 | 0.497 | 0.321 | 0.478 | |
| 7 | Q0Q468 | 176 | GFYAEG s RGGS | 0.151 | -0.235 | -1.103 | -0.396 | |
| 7 | Q0Q468 | 180 | EGSRGG s QASS | 0.091 | 0.442 | -0.934 | -0.134 | |
| 7 | Q0Q468 | 183 | RGGSQA s SRSS | 0.057 | -0.25 | -1.556 | -0.583 | |
| 7 | Q0Q468 | 184 | GGSQAS s RSSS | 0.089 | -0.257 | -1.301 | -0.49 | |
| 7 | Q0Q468 | 186 | SQASSR s SSRS | 0.059 | -0.177 | -1.437 | -0.518 | |
| 7 | Q0Q468 | 187 | QASSRS s SRSR | 0.078 | -0.25 | -1.266 | -0.479 | |
| 7 | Q0Q468 | 188 | ASSRSS s RSRG | 0.467 | 0.644 | -0.179 | 0.311 | |
| 7 | Q0Q468 | 190 | SRSSSR s RGNS | 0.2 | 0.22 | -1.014 | -0.198 | |
| 7 | Q0Q468 | 194 | SRSRGN s RNST | 0.32 | 0.805 | -0.255 | 0.29 | |
| 7 | Q0Q468 | 197 | RGNSRN s TPGS | 0.697 | 0.526 | 0.033 | 0.419 | likely, by analogy with SARS-CoV-2 |
| 7 | Q0Q468 | 198 | GNSRNS t PGSS | 0.215 | 0.381 | -0.438 | 0.053 | |
| 7 | Q0Q468 | 201 | RNSTPG s SRGN | 0.079 | -0.424 | -1.35 | -0.565 | |
| 7 | Q0Q468 | 202 | NSTPGS s RGNS | 0.064 | -0.303 | -1.458 | -0.566 | |
| 7 | Q0Q468 | 206 | GSSRGN s PARM | 0.317 | 0.338 | -0.567 | 0.029 | |
| 7 | Q0Q468 | 212 | SPARMA s GSGE | 0.35 | 0.52 | -0.218 | 0.217 | |
| 7 | Q0Q468 | 214 | ARMASG s GETA | 0.217 | 0.183 | -0.567 | -0.056 | |

| | | | | | | | | |
|---|---|---|---|---|---|---|---|---|
| 7 | Q0Q468 | 217 | ASGSGE t ALAL | 0.056 | -0.214 | -1.579 | -0.579 | |
| 7 | Q0Q468 | 232 | RLNQLE s KVSG | 0.15 | -0.053 | -1.051 | -0.318 | |
| 7 | Q0Q468 | 235 | QLESKV s GKGQ | 0.176 | -0.244 | -0.794 | -0.287 | |
| 7 | Q0Q468 | 245 | QQQQGQ t VTKK | 0.118 | -0.14 | -0.816 | -0.279 | |
| 7 | Q0Q468 | 247 | QQGQTV t KKSA | 0.092 | -0.151 | -0.76 | -0.273 | |
| 7 | Q0Q468 | 250 | QTVTKK s AAEA | 0.174 | -0.112 | -0.967 | -0.302 | |
| 7 | Q0Q468 | 255 | KSAAEA s KKPR | 0.152 | -0.249 | -0.909 | -0.335 | |
| 7 | Q0Q468 | 263 | KPRQKR t ATKS | 0.18 | 0.045 | -1.152 | -0.309 | |
| 7 | Q0Q468 | 265 | RQKRTA t KSYN | 0.534 | 0.885 | 0.261 | 0.56 | unlikely, by analogy with SARS-CoV-2 |
| 7 | Q0Q468 | 267 | KRTATK s YNVT | 0.206 | 0.183 | -0.397 | -0.003 | |
| 7 | Q0Q468 | 271 | TKSYNV t QAFG | 0.273 | -0.156 | -0.719 | -0.201 | |
| 7 | Q0Q468 | 282 | RRGPEQ t QGNF | 0.166 | 0.137 | -1.106 | -0.268 | |
| 7 | Q0Q468 | 296 | DLIRQG t DYKY | 0.638 | 0.913 | -0.145 | 0.469 | |
| 7 | Q0Q468 | 310 | IAQFAP s ASAF | 0.266 | 0.03 | -0.65 | -0.118 | |
| 7 | Q0Q468 | 312 | QFAPSA s AFFG | 0.36 | 0.155 | -0.221 | 0.098 | |
| 7 | Q0Q468 | 318 | SAFFGM s RIGM | 0.108 | -0.181 | -1.136 | -0.403 | |
| 7 | Q0Q468 | 325 | RIGMEV t PLGT | 0.092 | -0.432 | -1.361 | -0.567 | |
| 7 | Q0Q468 | 329 | EVTPLG t WLTY | 0.05 | -0.158 | -1.45 | -0.519 | |
| 7 | Q0Q468 | 332 | PLGTWL t YHGA | 0.191 | -0.057 | -0.729 | -0.198 | |
| 7 | Q0Q468 | 366 | YKAFPP t EPKK | 0.635 | 0.431 | 0.32 | 0.462 | |
| 7 | Q0Q468 | 376 | KDKKKK t DEAQ | 0.295 | -0.009 | -0.522 | -0.079 | |
| 7 | Q0Q468 | 390 | QRKKQP t VTLL | 0.378 | 0.249 | -0.37 | 0.086 | |
| 7 | Q0Q468 | 392 | KKQPTV t LLPA | 0.379 | 0.038 | -0.299 | 0.039 | |
| 7 | Q0Q468 | 403 | ADMDDF s RQLQ | 0.083 | -0.22 | -1.376 | -0.504 | |
| 7 | Q0Q468 | 409 | SRQLQN s MSGA | 0.138 | 0.195 | -1.176 | -0.281 | |
| 7 | Q0Q468 | 411 | QLQNSM s GASA | 0.283 | 0.267 | -0.12 | 0.143 | |
| 7 | Q0Q468 | 414 | NSMSGA s ADST | 0.08 | -0.166 | -1.258 | -0.448 | |
| 7 | Q0Q468 | 417 | SGASAD s TQA- | 0.194 | -0.088 | -0.858 | -0.251 | |
| 7 | Q0Q468 | 418 | GASADS t QA-- | 0.086 | -0.199 | -1.563 | -0.559 | |
| **NCAP_BC512 Nucleoprotein OS=Bat coronavirus 512/2005 OX=693999 GN=N PE=3 SV=1** | | | | | | | | |
| 8 | Q0Q462 | 3 | ---MA s VKFQ | 0.156 | -0.401 | -1.006 | -0.417 | |

| | | | | | | | |
|---|---|---|---|---|---|---|---|
| 8 | Q0Q462 | 12 | FQPRGR s KGRV | 0.404 | 0.732 | -0.318 | 0.273 |
| 8 | Q0Q462 | 19 | KGRVPL s LFAP | 0.233 | 0.12 | -0.302 | 0.017 |
| 8 | Q0Q462 | 27 | FAPLRV t DEKP | 0.239 | 0.031 | -0.971 | -0.234 |
| 8 | Q0Q462 | 73 | DRVDLP s NWHF | 0.128 | -0.004 | -1.32 | -0.399 |
| 8 | Q0Q462 | 82 | HFYFLG t GPHS | 0.204 | 0.303 | -0.414 | 0.031 |
| 8 | Q0Q462 | 86 | LGTGPH s DLPF | 0.12 | -0.087 | -0.834 | -0.267 |
| 8 | Q0Q462 | 94 | LPFRKR t DGVF | 0.453 | 0.833 | -0.109 | 0.392 |
| 8 | Q0Q462 | 107 | AIDGAK t QPTG | 0.704 | 0.532 | 0.154 | 0.463 |
| 8 | Q0Q462 | 110 | GAKTQP t GLGV | 0.121 | -0.23 | -1.389 | -0.499 |
| 8 | Q0Q462 | 117 | GLGVRK s SEKP | 0.223 | 0.128 | -0.683 | -0.111 |
| 8 | Q0Q462 | 118 | LGVRKS s EKPL | 0.291 | 0.73 | 0.019 | 0.347 |
| 8 | Q0Q462 | 140 | VEIVEP t TPNN | 0.299 | 0.335 | -0.877 | -0.081 |
| 8 | Q0Q462 | 141 | EIVEPT t PNNS | 0.039 | -0.33 | -1.469 | -0.587 |
| 8 | Q0Q462 | 145 | PTTPNN s RANS | 0.184 | -0.16 | -1.125 | -0.367 |
| 8 | Q0Q462 | 149 | NNSRAN s RSRS | 0.496 | 0.606 | -0.258 | 0.281 |
| 8 | Q0Q462 | 151 | SRANSR s RSRG | 0.133 | 0.105 | -1.007 | -0.256 |
| 8 | Q0Q462 | 153 | ANSRSR s RGGQ | 0.605 | 0.765 | 0.212 | 0.527 |
| 8 | Q0Q462 | 158 | RSRGGQ s NSRG | 0.163 | -0.042 | -1.089 | -0.323 |
| 8 | Q0Q462 | 160 | RGGQSN s RGNS | 0.452 | 0.22 | -0.37 | 0.101 |
| 8 | Q0Q462 | 164 | SNSRGN s QNRG | 0.401 | 0.554 | -0.458 | 0.166 |
| 8 | Q0Q462 | 171 | QNRGDK s RNQS | 0.267 | 0.022 | -0.342 | -0.018 |
| 8 | Q0Q462 | 175 | DKSRNQ s RNRS | 0.758 | 0.735 | 0.336 | 0.61 |
| 8 | Q0Q462 | 179 | NQSRNR s QSND | 0.418 | 0.731 | -0.242 | 0.302 |
| 8 | Q0Q462 | 181 | SRNRSQ s NDRG | 0.77 | 1.156 | 0.417 | 0.781 |
| 8 | Q0Q462 | 186 | QSNDRG s DSRD | 0.179 | -0.13 | -1.301 | -0.417 |
| 8 | Q0Q462 | 188 | NDRGSD s RDDL | 0.513 | 0.317 | -0.073 | 0.252 |
| 8 | Q0Q462 | 214 | AKPKGK t QSGK | 0.217 | -0.213 | -0.882 | -0.293 |
| 8 | Q0Q462 | 216 | PKGKTQ s GKNT | 0.232 | 0.06 | -0.59 | -0.099 |
| 8 | Q0Q462 | 220 | TQSGKN t PKNK | 0.047 | -0.445 | -1.531 | -0.643 |
| 8 | Q0Q462 | 225 | NTPKNK s RSGS | 0.202 | -0.215 | -0.843 | -0.285 |
| 8 | Q0Q462 | 227 | PKNKSR s GSVQ | 0.394 | 0.176 | -0.504 | 0.022 |
| 8 | Q0Q462 | 229 | NKSRSG s VQRA | 0.832 | 0.967 | 0.786 | 0.862 |
| 8 | Q0Q462 | 244 | KPEWRR t PSGD | 0.109 | -0.387 | -1.196 | -0.491 |

| | | | | | | | |
|---|---|---|---|---|---|---|---|
| 8 | Q0Q462 | 246 | EWRRTP s GDES | 0.667 | 1.065 | 0.519 | 0.75 |
| 8 | Q0Q462 | 250 | TPSGDE s VEVC | 0.061 | -0.274 | -1.621 | -0.611 |
| 8 | Q0Q462 | 261 | FGPRGG t RNFG | 0.228 | 0.569 | -0.408 | 0.13 |
| 8 | Q0Q462 | 266 | GTRNFG s SEFV | 0.167 | -0.001 | -1.281 | -0.372 |
| 8 | Q0Q462 | 267 | TRNFGS s EFVA | 0.136 | 0.123 | -0.916 | -0.219 |
| 8 | Q0Q462 | 284 | GYAQAA s LVPG | 0.667 | 0.217 | 0.196 | 0.36 |
| 8 | Q0Q462 | 300 | FGGNVA t KEMA | 0.149 | -0.078 | -0.801 | -0.243 |
| 8 | Q0Q462 | 310 | ADGVEI t YTYK | 0.049 | -0.412 | -1.773 | -0.712 |
| 8 | Q0Q462 | 312 | GVEITY t YKML | 0.097 | 0.028 | -0.856 | -0.244 |
| 8 | Q0Q462 | 348 | QRKVKR s RTPT | 0.078 | -0.1 | -1.338 | -0.453 |
| 8 | Q0Q462 | 350 | KVKRSR t PTPK | 0.51 | 0.644 | 0.066 | 0.407 |
| 8 | Q0Q462 | 352 | KRSRTP t PKPA | 0.224 | 0.575 | -0.281 | 0.173 |
| 8 | Q0Q462 | 357 | PTPKPA t EPVY | 0.775 | 0.559 | 0.229 | 0.521 |
| 8 | Q0Q462 | 369 | DVAADP t YANL | 0.098 | -0.203 | -1.245 | -0.45 |
| 8 | Q0Q462 | 377 | ANLEWD t TVED | 0.131 | -0.197 | -1.135 | -0.4 |
| 8 | Q0Q462 | 378 | NLEWDT t VEDG | 0.456 | 0.062 | -0.165 | 0.118 |
| 8 | Q0Q462 | 392 | INEVFD t QN-- | 0.115 | -0.17 | -1.294 | -0.45 |
| SARS Nucleoprotein OS=Bat coronavirus RaTG13 OX=2709072 GN=N PE=3 SV=1 | | | | | | | |
| 9 | A0A6B9WI45 | 2 | ----M s DNGP | 0.33 | -0.143 | -0.5 | -0.104 |
| 9 | A0A6B9WI45 | 16 | RNAPRI t FGGP | 0.413 | 0.031 | -0.125 | 0.106 |
| 9 | A0A6B9WI45 | 21 | ITFGGP s DSTG | 0.114 | -0.232 | -1.206 | -0.441 |
| 9 | A0A6B9WI45 | 23 | FGGPSD s TGSN | 0.194 | 0.089 | -0.704 | -0.14 |
| 9 | A0A6B9WI45 | 24 | GGPSDS t GSNQ | 0.045 | -0.369 | -1.809 | -0.711 |
| 9 | A0A6B9WI45 | 26 | PSDSTG s NQNG | 0.075 | -0.202 | -1.54 | -0.556 |
| 9 | A0A6B9WI45 | 33 | NQNGER s GARP | 0.038 | -0.379 | -1.821 | -0.721 |
| 9 | A0A6B9WI45 | 49 | QGLPNN t ASWF | 0.131 | -0.112 | -0.879 | -0.287 |
| 9 | A0A6B9WI45 | 51 | LPNNTA s WFTA | 0.106 | 0 | -0.849 | -0.247 |
| 9 | A0A6B9WI45 | 54 | NTASWF t ALTQ | 0.154 | -0.094 | -1.04 | -0.327 |
| 9 | A0A6B9WI45 | 57 | SWFTAL t QHGK | 0.139 | -0.161 | -1.293 | -0.438 |
| 9 | A0A6B9WI45 | 76 | QGVPIN t NSSP | 0.045 | -0.299 | -1.527 | -0.594 |
| 9 | A0A6B9WI45 | 78 | VPINTN s SPDD | 0.539 | 0.531 | 0.008 | 0.36 |

| | | | | | | | | |
|---|---|---|---|---|---|---|---|---|
| 9 | A0A6B9WI45 | 79 | PINTNS s PDDQ | 0.133 | -0.282 | -1.115 | -0.421 | |
| 9 | A0A6B9WI45 | 91 | GYYRRA t RRIR | 0.873 | 0.98 | 0.702 | 0.852 | unlikely, by analogy with SARS-CoV-2 |
| 9 | A0A6B9WI45 | 105 | GKMKDL s PRWY | 0.268 | -0.197 | -0.726 | -0.219 | |
| 9 | A0A6B9WI45 | 115 | YFYYLG t GPEA | 0.36 | 0.343 | -0.117 | 0.195 | |
| 9 | A0A6B9WI45 | 135 | GIIWVA t EGAL | 0.649 | 0.356 | 0.089 | 0.365 | |
| 9 | A0A6B9WI45 | 141 | TEGALN t PKDH | 0.051 | -0.518 | -1.848 | -0.772 | |
| 9 | A0A6B9WI45 | 148 | PKDHIG t RNPA | 0.18 | -0.208 | -0.713 | -0.247 | |
| 9 | A0A6B9WI45 | 165 | LQLPQG t TLPK | 0.035 | -0.37 | -1.556 | -0.631 | |
| 9 | A0A6B9WI45 | 166 | QLPQGT t LPKG | 0.615 | 0.497 | 0.321 | 0.478 | unlikely, by analogy with SARS-CoV-2 |
| 9 | A0A6B9WI45 | 176 | GFYAEG s RGGS | 0.151 | -0.235 | -1.103 | -0.396 | |
| 9 | A0A6B9WI45 | 180 | EGSRGG s QASS | 0.091 | 0.442 | -0.934 | -0.134 | |
| 9 | A0A6B9WI45 | 183 | RGGSQA s SRSS | 0.057 | -0.25 | -1.556 | -0.583 | |
| 9 | A0A6B9WI45 | 184 | GGSQAS s RSSS | 0.089 | -0.257 | -1.301 | -0.49 | |
| 9 | A0A6B9WI45 | 186 | SQASSR s SSRS | 0.059 | -0.177 | -1.437 | -0.518 | |
| 9 | A0A6B9WI45 | 187 | QASSRS s SRSR | 0.078 | -0.25 | -1.266 | -0.479 | |
| 9 | A0A6B9WI45 | 188 | ASSRSS s RSRN | 0.458 | 0.659 | -0.232 | 0.295 | |
| 9 | A0A6B9WI45 | 190 | SRSSSR s RNSS | 0.106 | 0.111 | -1.133 | -0.305 | |
| 9 | A0A6B9WI45 | 193 | SSRSRN s SRNS | 0.364 | 0.152 | -0.835 | -0.107 | |
| 9 | A0A6B9WI45 | 194 | SRSRNS s RNST | 0.277 | 0.751 | -0.208 | 0.274 | |
| 9 | A0A6B9WI45 | 197 | RNSSRN s TPGS | 0.648 | 0.444 | -0.003 | 0.363 | likely, by analogy with SARS-CoV-2 |
| 9 | A0A6B9WI45 | 198 | NSSRNS t PGSS | 0.195 | 0.366 | -0.47 | 0.03 | |
| 9 | A0A6B9WI45 | 201 | RNSTPG s SRGT | 0.059 | -0.495 | -1.388 | -0.608 | |
| 9 | A0A6B9WI45 | 202 | NSTPGS s RGTS | 0.058 | -0.37 | -1.375 | -0.562 | |
| 9 | A0A6B9WI45 | 205 | PGSSRG t SPAR | 0.597 | 0.49 | 0.145 | 0.411 | |
| 9 | A0A6B9WI45 | 206 | GSSRGT s PARM | 0.239 | 0.287 | -0.575 | -0.016 | |
| 9 | A0A6B9WI45 | 215 | RMAGNG s DAAL | 0.164 | 0.115 | -0.698 | -0.14 | |
| 9 | A0A6B9WI45 | 232 | RLNQLE s KMSG | 0.177 | -0.048 | -1.007 | -0.293 | |
| 9 | A0A6B9WI45 | 235 | QLESKM s GKGQ | 0.157 | -0.118 | -0.754 | -0.238 | |
| 9 | A0A6B9WI45 | 243 | KGQQQQ s QTVT | 0.133 | -0.119 | -1.098 | -0.361 | |
| 9 | A0A6B9WI45 | 245 | QQQQSQ t VTKK | 0.345 | 0.15 | -0.141 | 0.118 | |
| 9 | A0A6B9WI45 | 247 | QQSQTV t KKSA | 0.111 | -0.159 | -0.662 | -0.237 | |
| 9 | A0A6B9WI45 | 250 | QTVTKK s AAEA | 0.174 | -0.112 | -0.967 | -0.302 | |
| 9 | A0A6B9WI45 | 255 | KSAAEA s KKPR | 0.152 | -0.249 | -0.909 | -0.335 | |

| | | | | | | | | |
|---|---|---|---|---|---|---|---|---|
| 9 | A0A6B9WI45 | 263 | KPRQKR t ATKQ | 0.188 | 0.023 | -1.163 | -0.317 | |
| 9 | A0A6B9WI45 | 265 | RQKRTA t KQYN | 0.672 | 0.963 | 0.549 | 0.728 | unlikely, by analogy with SARS-CoV-2 |
| 9 | A0A6B9WI45 | 271 | TKQYNV t QAFG | 0.329 | -0.068 | -0.616 | -0.119 | |
| 9 | A0A6B9WI45 | 282 | RRGPEQ t QGNF | 0.166 | 0.137 | -1.106 | -0.268 | |
| 9 | A0A6B9WI45 | 296 | ELIRQG t DYKH | 0.325 | 0.753 | -0.495 | 0.194 | |
| 9 | A0A6B9WI45 | 310 | IAQFAP s ASAF | 0.266 | 0.03 | -0.65 | -0.118 | |
| 9 | A0A6B9WI45 | 312 | QFAPSA s AFFG | 0.36 | 0.155 | -0.221 | 0.098 | |
| 9 | A0A6B9WI45 | 318 | SAFFGM s RIGM | 0.108 | -0.181 | -1.136 | -0.403 | |
| 9 | A0A6B9WI45 | 325 | RIGMEV t PSGT | 0.086 | -0.467 | -1.384 | -0.588 | |
| 9 | A0A6B9WI45 | 327 | GMEVTP s GTWL | 0.102 | 0.025 | -0.991 | -0.288 | |
| 9 | A0A6B9WI45 | 329 | EVTPSG t WLTY | 0.139 | 0.111 | -0.766 | -0.172 | |
| 9 | A0A6B9WI45 | 332 | PSGTWL t YTGA | 0.105 | -0.254 | -1.079 | -0.409 | |
| 9 | A0A6B9WI45 | 334 | GTWLTY t GAIK | 0.129 | 0.044 | -0.948 | -0.258 | |
| 9 | A0A6B9WI45 | 362 | HIDAYK t FPPT | 0.694 | 0.583 | 0.571 | 0.616 | |
| 9 | A0A6B9WI45 | 366 | YKTFPP t EPKK | 0.643 | 0.44 | 0.307 | 0.463 | |
| 9 | A0A6B9WI45 | 379 | KKKADE t QALP | 0.145 | -0.235 | -1.253 | -0.448 | |
| 9 | A0A6B9WI45 | 391 | RQKKQQ t VTLL | 0.37 | 0.144 | -0.312 | 0.068 | |
| 9 | A0A6B9WI45 | 393 | KKQQTV t LLPA | 0.492 | 0.099 | -0.074 | 0.172 | |
| 9 | A0A6B9WI45 | 404 | ADLDDF s KQLQ | 0.113 | -0.181 | -1.087 | -0.385 | |
| 9 | A0A6B9WI45 | 410 | SKQLQQ s MSSA | 0.076 | 0.008 | -1.337 | -0.418 | |
| 9 | A0A6B9WI45 | 412 | QLQQSM s SADS | 0.6 | 0.402 | 0.166 | 0.389 | |
| 9 | A0A6B9WI45 | 413 | LQQSMS s ADST | 0.062 | -0.211 | -1.171 | -0.44 | |
| 9 | A0A6B9WI45 | 416 | SMSSAD s TQA- | 0.15 | 0.005 | -1.099 | -0.315 | |
| 9 | A0A6B9WI45 | 417 | MSSADS t QA-- | 0.084 | -0.169 | -1.435 | -0.507 | |

**Supplementary table 4. Primer sequences used in this study.**

| **Primer Name** | **Primer sequence (5’ – 3’)** |
|---|---|
| N.Full_F | CCAGGGACCAGCAATGTCTGATAATGGACCCCAA |
| N.Full_R | GAGGAGAAGGCGCGTTAGGCCTGAGTTGAGTCA |
| N.1-211_R | GAGGAGAAGGCGCGTTAAGCCATTCTAGCAGGAGA |
| N.1-238_R | GAGGAGAAGGCGCGTTAGCCTTTACCAGACATTTTGC |
| N.212-419_F | CCAGGGACCAGCAATGGGCAATGGCGGTGATG |
| N.1-179_BamHI_rev | TATATGGATCCTTAGCCGCCTCTGCTCCCTTC |
| N.1-196_BamHI_rev | TATATGGATCCTTAATTTCTTGAACTGTTGCGAC |
| N.1-204_BamHI_rev | TATATGGATCCTTATCCCCTACTGCTGCCTGGAG |

**Supplementary table 5. Properties of the studied proteins.**

| Protein | Mw, Da | A, ml/(mg*cm) | ε, 1/(M*cm) | pI |
|---|---|---|---|---|
| N_His$_6$_Full-length (1-419) | 48064 | 0.91 | 43890 | 10.03 |
| N_Full-length (1-419) | 45851 | 0.93 | 42530 | 10.07 |
| N_His$_6$_NTD (1-238) | 27766 | 0.94 | 26030 | 10.43 |
| N_NTD (1-238) | 25620 | 1.02 | 26030 | 10.55 |
| N_His$_6$_NTD (1-211) | 25042 | 1.04 | 26030 | 10.63 |
| N_NTD (1-211) | 22896 | 1.14 | 26030 | 10.77 |
| N_His$_6$_NTD (1-204) | 24328 | 1.07 | 26030 | 10.50 |
| N_NTD (1-204) | 22181 | 1.17 | 26030 | 10.63 |
| N_His$_6$_NTD (1-196) | 23598 | 1.10 | 26030 | 10.38 |
| N_NTD (1-196) | 21452 | 1.21 | 26030 | 10.50 |
| N_His$_6$_NTD (1-179) | 21762 | 1.20 | 26030 | 9.93 |
| N_NTD (1-179) | 19744 | 1.32 | 26030 | 10.04 |
| N_His$_6$_CTD (212-419) | 25476 | 0.65 | 16500 | 9.54 |
| N_CTD (212-419) | 23329 | 0.71 | 16500 | 9.62 |
| N_His$_6$_CTD (247-364) | 15309 | 1.08 | 16500 | 9.65 |
| human 14-3-3γ (1-247) | 28303 | 1.13 | 31860 | 4.80 |
| human His$_6$-14-3-3ε (1-255) | 32548 | 0.93 | 30370 | 4.85 |
| human 14-3-3ζ (1-245) | 27745 | 0.99 | 27390 | 4.73 |
| human His$_6$-14-3-3γΔC (2-238) | 30669 | 1.23 | 37820 | 5.19 |
| human 14-3-3βΔC (1-232) | 27084 | 1.01 | 27390 | 5.06 |
| human 14-3-3τΔC (1-230) | 26718 | 1.03 | 27390 | 4.95 |
| human 14-3-3σΔC (1-231) | 26106 | 0.99 | 25900 | 4.85 |
| human His$_6$-14-3-3ηΔC (1-235) | 30286 | 1.15 | 34840 | 5.20 |
| human 14-3-3εΔC (1-232) | 26670 | 1.08 | 28880 | 4.92 |
| human 14-3-3ζm-S58E ($^{12}$LAE$^{14}$→QQR) | 27886 | 0.98 | 27390 | 4.78 |

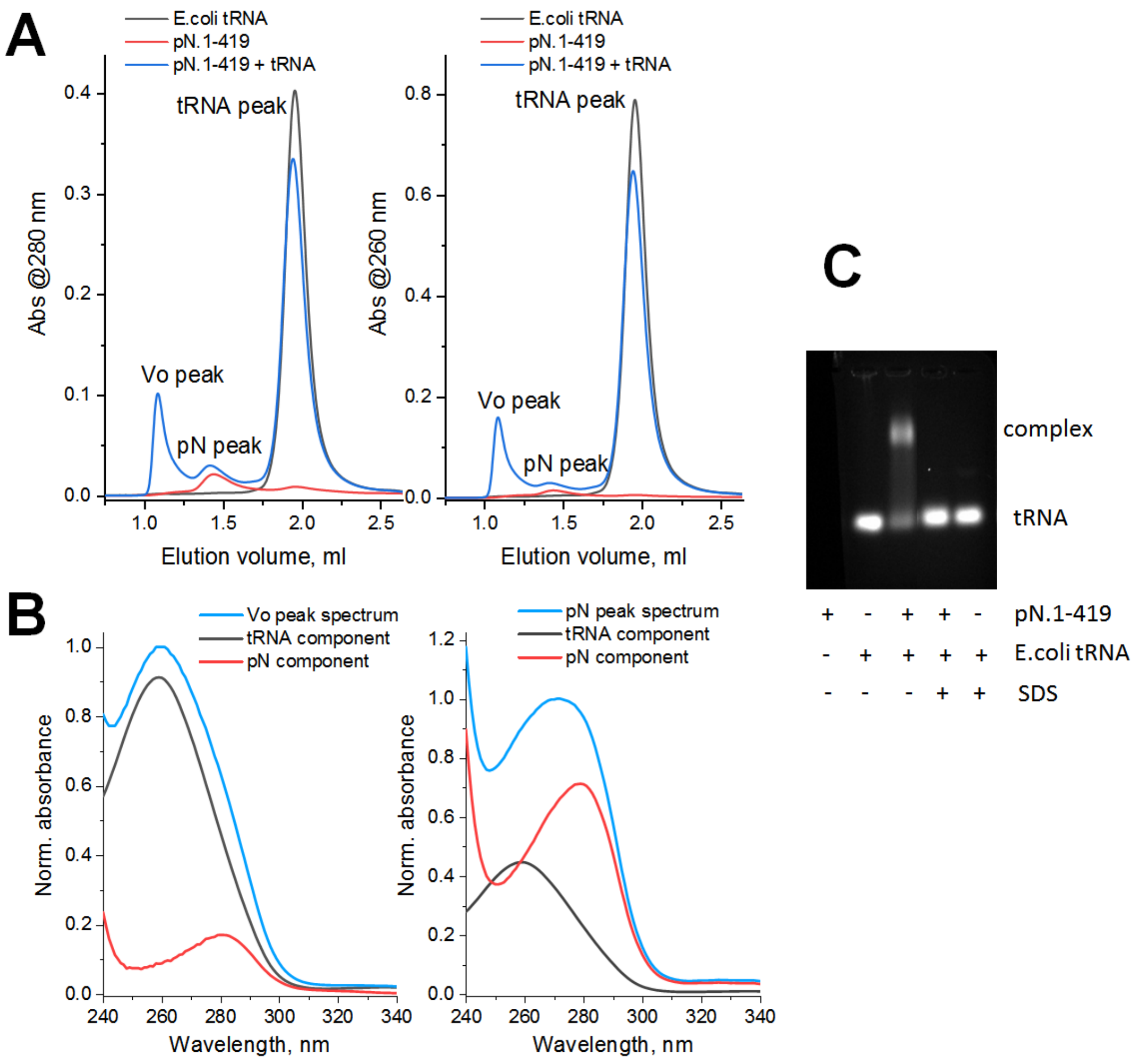

**Supplementary Fig. 1**. Phosphorylated N binds *E.coli* tRNA and forms oligomers containing both tRNA and pN. A. SEC profiles run at 200 mM NaCl and followed by 260 and 280-nm detection show the interaction between pN and tRNA. B. Absorbance spectra of the Vo or pN fractions as retrieved from the diode array detection data obtained during the SEC runs. C. 1% agarose gel showing the interaction of pN with *E.coli* tRNA. Note that the addition of SDS denatures the protein and releases tRNA. The experiment was performed twice and the most typical results are presented.

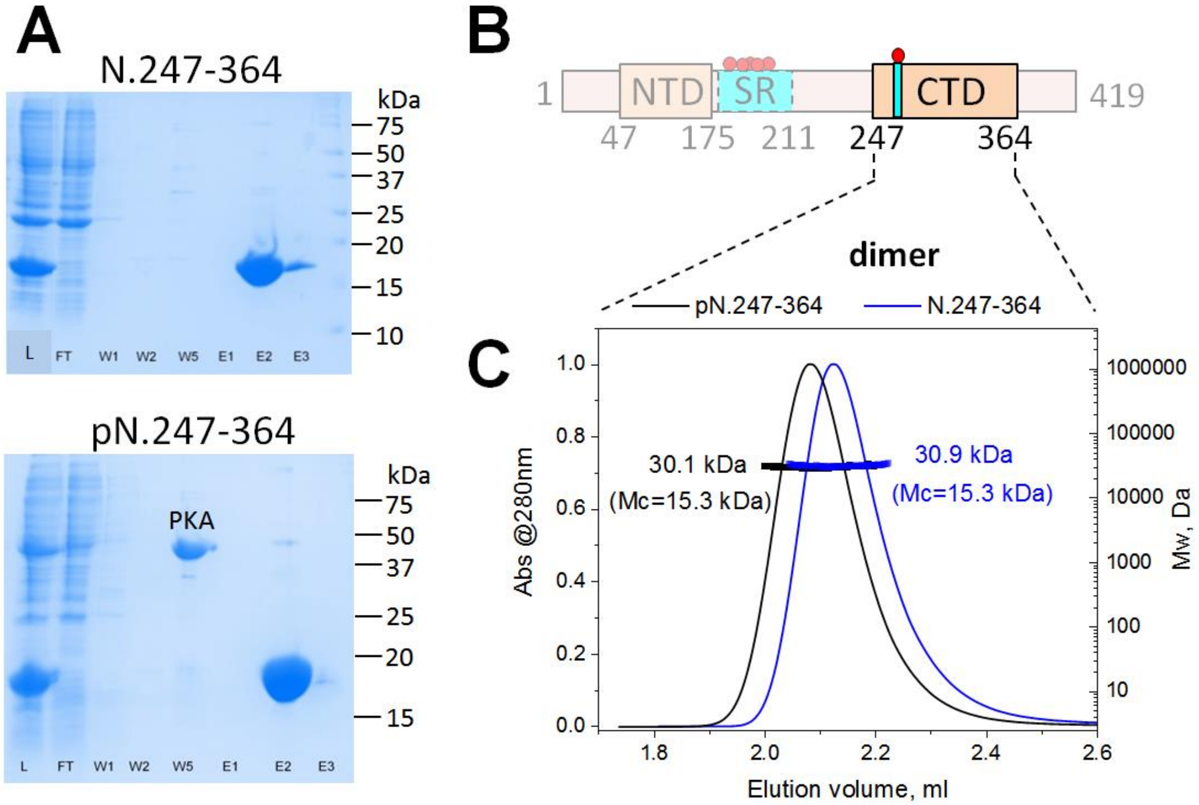

**Supplementary Fig. 2**. Purification and SEC-MALS analysis of the unphosphorylated and phosphorylated versions of the N.247-364 construct. A. Immobilized metal-affinity chromatography of N.247-364 and pN.247-364 proteins. L - loaded fraction, FL - flowthrough, W1, W2, W5 - wash at 10, 20 and 50 mM imidazole, respectively, E1-E3 - eluted fractions. Mw markers are shown on the right in kDa. Note the presence of PKA in the W5 fraction of only pN.247-364. B. Schematic representation of the N protein sequence with the main domains/regions highlighted. C. SEC-MALS of N.247-364 and pN.247-364 proteins using a Superdex 200 Increase 5/150 column operated at a 0.45 ml/min flow rate. Note the significant shift of the peak of pN.247-364 due to phosphorylation. Given the same mass corresponding to the dimeric form of the protein, this shift indicates the increase of the apparent size. The experiment was repeated twice and the most typical results are presented.

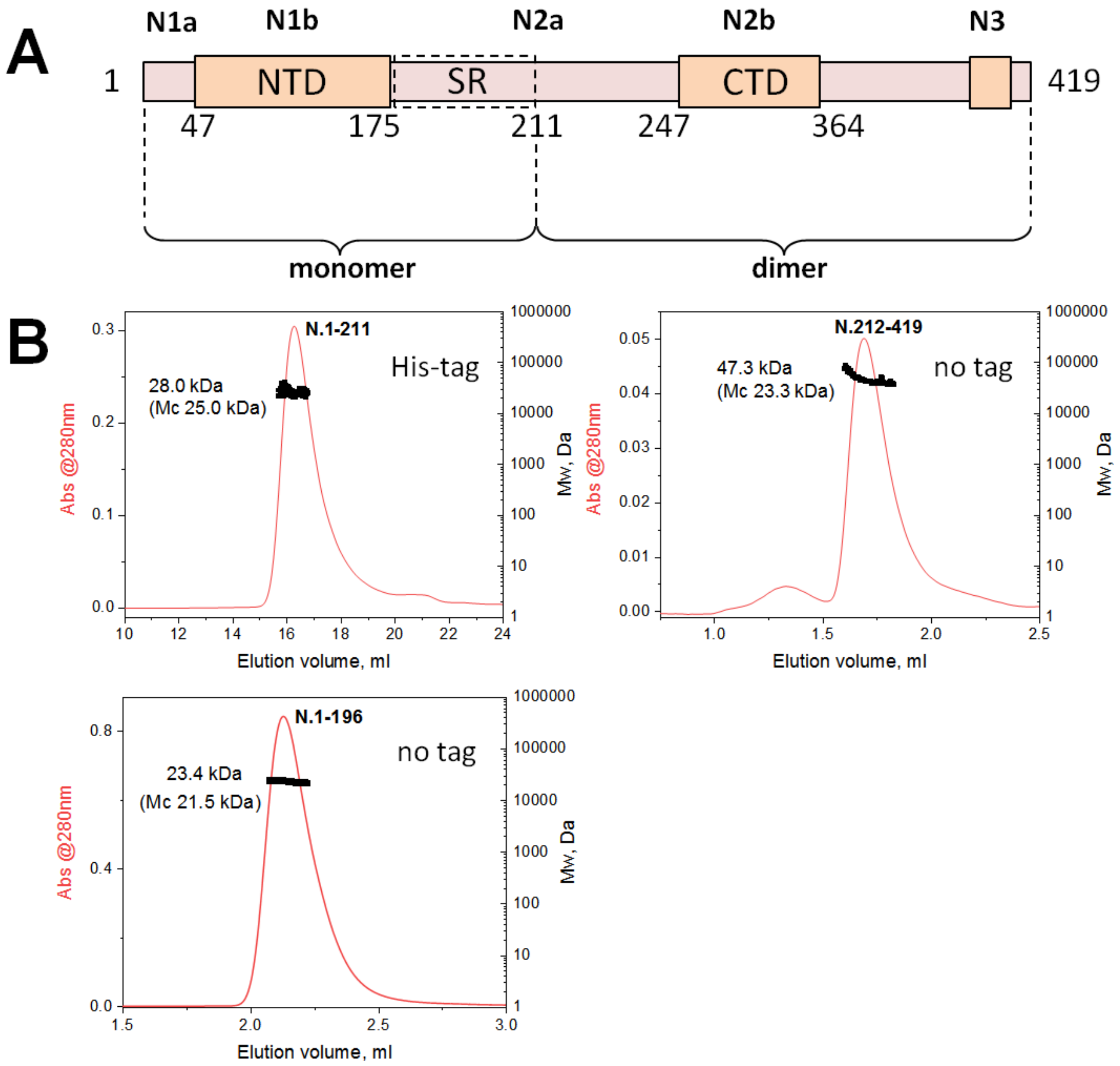

**Supplementary Fig. 3**. Oligomeric state of the N- and C-terminal halves of N studied by SEC-MALS. A. Schematic representation of the N protein sequence with the main domains/regions highlighted. B. SEC-MALS of N.1-211, N.1-196 and N.212-419 proteins using a Superdex 200 Increase 10/300 column operated at a 0.8 ml/min flow rate (N.1-211) or a Superdex 200 Increase 5/150 column operated at a 0.45 ml/min flow rate (N.1-196 and N.212-419). Mw distribution across the protein peaks are shown with the average Mw value. For comparison, Mw calculated from protein sequence is also indicated (Mc). The experiment was repeated twice and the most typical results are presented.
